# Synthetic antifreeze glycoproteins with potent ice-binding activity

**DOI:** 10.1101/2023.09.07.556704

**Authors:** Anna C. Deleray, Simranpreet S. Saini, Alexander C. Wallberg, Jessica R. Kramer

## Abstract

Antifreeze glycoproteins (AFGPs) are produced by extremophiles to defend against tissue damage in freezing climates. Cumbersome isolation from polar fish has limited probing AFGP molecular mechanisms of action and limited developing bioinspired cryoprotectants for application in agriculture, foods, coatings, and biomedicine. Here, we present a rapid, scalable, and tunable route to synthetic AFGPs (sAFGPs) using *N*-carboxyanhydride polymerization. Our materials are the first mimics to harness the molecular size, chemical motifs, and long-range conformation of native AFGPs. We found that ice-shaping and ice-recrystallization inhibition activity increases with chain length and Ala is a key residue. Glycan structure had only minor effects and all glycans examined displayed antifreeze activity. The sAFGPs are biodegradable, non-toxic, and internalized into endocytosing cells. sAFGPs were found to be bystanders in cryopreservation of human red blood cells. Overall, our sAFGPs functioned as surrogates for bona fide AFGPs, solving a long-standing challenge in access to natural antifreeze materials.

## Introduction

Extremophile organisms in cold climates produce specialized antifreeze proteins to protect their tissues from freezing damage.^1–4^ These proteins bind to ice to lower plasma’s freezing point and modify ice crystal shape and size to prevent mechanical damage to tissues.^5–8^ Antifreeze proteins are of great interest for applications in food technology, agriculture, fisheries, coatings, and in the petroleum industry.^9,10^ They have even come to market as texture-improving ice cream additives.^11^ A particularly impactful potential application is to improve post-thaw tissue viability and function in biomedical cryopreservation.^12,13^

To date, five classes of antifreeze proteins have been identified, including a subgroup called antifreeze glycoproteins (AFGPs) which are the most potent ice-shaping molecules ever discovered.^14,15^ Identified as the major serum protein of certain Antarctic and Arctic fish, AFGPs are a fascinating and rare example of convergent evolution.^16^ AFGPs consist of a highly conserved Ala-Ala-Thr repeat, with Thr bearing the disaccharide βGal(1→3)αGalNAc (Fig. 1A).^7^ Isoforms from 2.6–33.7 kDa, classified as AFGP8– AFGP1, are encoded within polyprotein genes.^16^ AFGPs adopt extended polyproline type II (PPII) helical conformations^17,18^, similar to Pro-rich proteins like collagen and mucins^19^ (Fig. 1A). Compared to the well-known right-handed α-helix with 3.6 residues/turn, PPII-helix is left-handed and has 3 residues/turn.^17,18^

**Figure 1.**
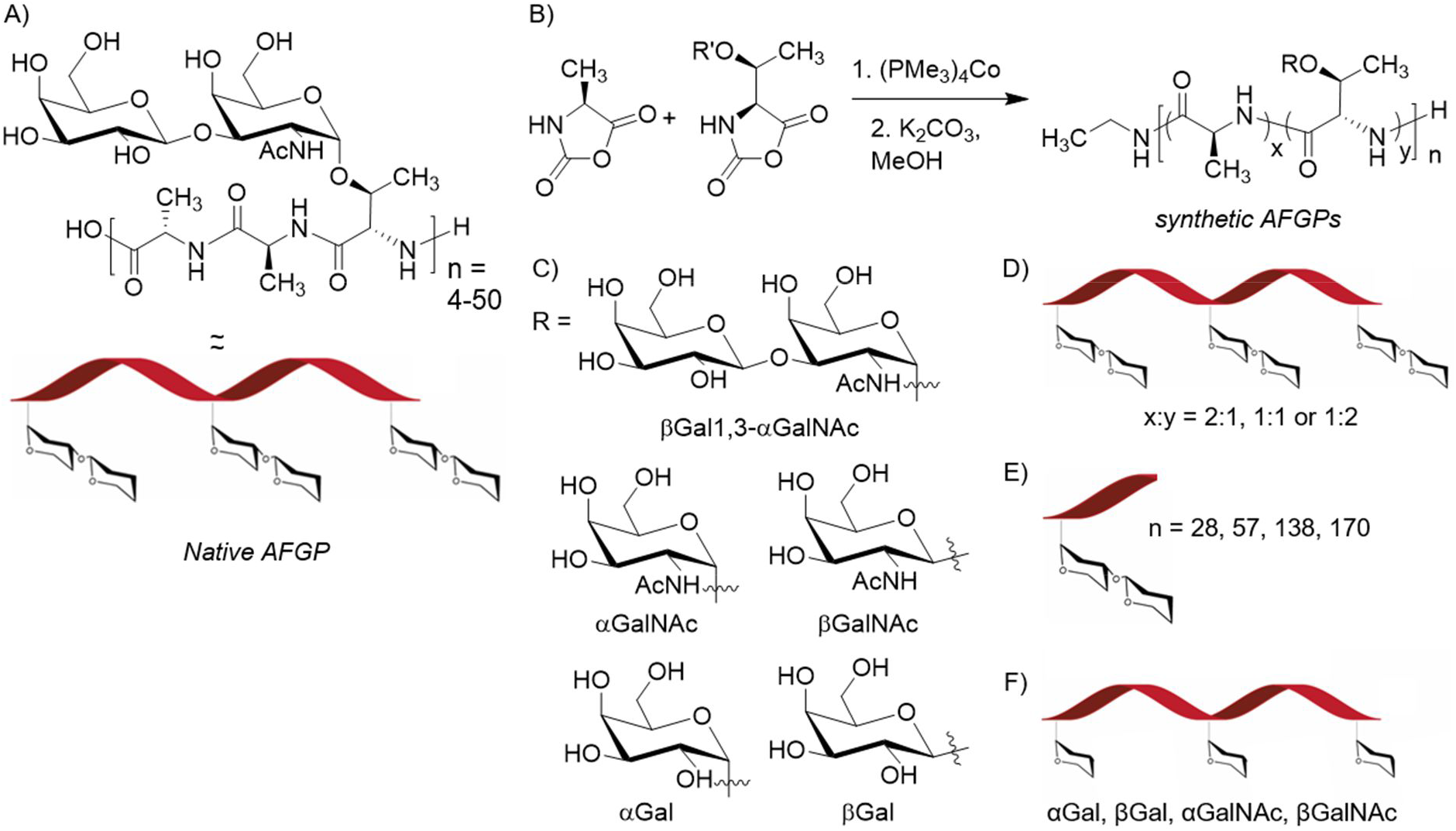
Structure of native AFGPs and preparation of sAFGP panel. A) Chemical structure of the AFGP tripeptide repeat and native PPII helical conformation. B) Preparation of tunable sAFGPs by NCA polymerization using transition metal catalysis. C) Structures of the five glycans utilized in the sAFGP panel. sAFGPs were prepared with varying D) ratios of Ala:glyco-Thr, E) molecular weights, and F) glycans.

AFGPs possess two well-established properties: ice recrystallization inhibition (IRI) and thermal hysteresis (TH). IRI prevents the growth of large ice crystals, while TH produces a non-colligative freezing point depression and gap between solution freezing and melting points.^6,8^ Effects are observable at concentrations 300–100,000x lower than dissolved salts and sugars.^20^ Overall, these properties render AFGPs highly attractive for cryopreservation applications in biomedical, agricultural, and food sectors.^9–13^

Despite their potential, there are no commercial AFGP sources. Isolating the proteins requires a polar fishing trip followed by labor-intensive fractionation that still yields heterogeneous mixtures of isoforms and potential contamination with strong allergens.^15^ Additionally, high molecular weight (MW) native AFGPs (50–150 residues, 10.5–34 kDa) are much more active than low MW AFGPs (12–38 residues, 2.6–7.9 kDa) but are of low abundance.^6,8,15,21^ The proteins have so far resisted recombinant production.^9,10,22^ Synthetic approaches using solid-phase synthesis or step-growth polymerizations have been beautifully explored, but are laborious and limited to low-activity, low-MW AFGPs.^10,23–35^ Additionally, many structures utilized non-native residues/glycans that could affect biocompatibility. Poly(vinyl alcohol) (PVA) has been investigated as a polyhydroxylated surrogate^36^ while polyproline has been utilized to capture the PPII conformation^37^. These structures are reportedly orders of magnitude less active than even low MW AFGPs. PVA has shown mixed results in combination with traditional cryoprotectants.^36,38,39^ Various other mimics have been investigated but have shown little or no IRI activity.^22,32,40–44^

A need remains for an accessible route to AFGPs, and that allows molecular tuning to probe the characteristics that drive ice-binding. Here, we present a rapid, scalable, and tunable route to AFGPs (sAFGPs) using amino acid *N*-carboxyanhydride (NCA) polymerization (Fig. 1B–F). Our building blocks were prepared on gram-scale and the NCA method is used commercially.^45^ The approach allows precise control over polypeptide backbone MW, composition, and glycosylation.^46,47^ We tuned these factors to optimize activity and to reveal long-debated molecular drivers of antifreeze activity. Finally, we probed the interaction of sAFGPs with model biological systems.

## Results and Discussion

### Design and synthesis of sAFGP panel

Despite nearly 70 years of study, the molecular details of AFGP ice-binding are debated. The generally accepted mechanism involves a protein ice-binding face and a non-binding face.^48–50^ After adsorption to an embryonic ice crystal, the non-binding face disorders approaching liquid water molecules, resulting in inhibited crystal growth, ice shaping, and lowered melting point. However, the roles of hydrophobic Ala versus hydrophilic sugars in these faces are unclear.

Molecular dynamics modeling of a 14-residue AFGP predicted the PPII structure is crucial and that Ala methyls nest into ice-surface cavities driven by the entropy of dehydration.^51^ An alternative model proposed amphipathic binding where Ala methyls and glycan hydroxyls cooperatively bind ice.^52^ Ice-binding via the hydroxyls alone has also been proposed based on observation of fluorescently-labeled AFGPs on ice crystal surfaces and because activity was disrupted by sugar-complexing borate.^5,6,53,54^ Additionally, AFGPs adsorb more efficiently to hydrophilic surfaces^55,56^ and monolayers of βGal(1→3)αGalNAc alone shape ice.^57^

Prior work indicated loss of activity due to oxidation, alkylation, and borylation of AFGP disaccharides.^14,58,59^ Tachibana et al. conducted the most comprehensive study of glycan structure to date.^34^ They compared 6–9 residue AFGP fragments bearing the α vs β-linked native disaccharide, αGal vs αGalNAc, and other glycans. They observed significant differences in TH and IRI behavior and identified the α-linkage and the C2 NHAc group as particularly important for activity. However, their peptides could only make 2–3 PPII-helical turns, and this assumes end-group participation. Experimental evidence using native AFGPs indicates activity is strongly dependent upon MW. However, data is convoluted since, due to purification challenges, experiments were conducted on pooled fractions.

To explore AFGP structure-function, we synthesized sAFGPs via NCA polymerization using our established methods.^46,47^ We varied chain length, glycosylation pattern, and hydrophobicity (Fig. 1B–F). To explore the role of MW, 28–170mers were prepared (Fig. 1E). To probe the role of Ala methyls vs. glycan hydroxyls, we prepared glycopolypeptides with varied densities of the two residues. The native amino acid ratio of 2:1 Ala:Thr was increased to 1:1 or 1:2 (Fig. 1D).

To explore glycosylation’s role in PPII structure and sAFGPs antifreeze activity, we prepared glyco-Thr conjugates bearing αGal, βGal, αGalNAc, βGalNAc, or native disaccharide βGal(1→3)αGalNAc (abbreviated as βGalαGalNAc) (Fig. 1C, F). These glycans might differently affect hydrogen-bonding or protein secondary structure.^47,60,61^ NMR studies on short αGalNAc-Thr/Ser peptides suggested an intramolecular hydrogen bond between the sugar *N*-Ac and Thr carbonyl is a stabilizing force for the PPII structure. However, our recent CD studies of high MW glycosylated polyThr indicated that the PPII conformation is adopted in structures lacking the *N*-Ac group (i.e. Gal instead of GalNAc).

Ala NCA was prepared from Ala in one step by treatment with phosgene in tetrahydrofuran (THF).^46^ Peracetylated glyco-Thr conjugates were prepared using literature protocols (See Supplemental Information (SI)).^26,47,62^ All conjugates were converted to NCAs from their tert-butyloxycarbonyl-forms by treatment with triphosgene and triethylamine in THF, via our previously optimized conditions.^47^ Direct phosgenation of glyco-Thrs in the same manner as Ala results in poor yields. In all cases, NCAs were isolated by anhydrous chromatography to give crystalline monomer.^63^

Glyco-Thr and Ala NCAs were converted to sAFGPs using (PMe_3_)_4_Co catalyst in THF (Fig. 1B). NCA:catalyst ratios were varied to tune degree of polymerization (DP) and monomer feed ratios varied to tune composition. Complete monomer consumption was evidenced by infrared spectroscopy. NCA carbonyl stretches at ∼1850 and 1790 cm^-1^ disappeared, while peptide carbonyl stretches at ∼1650 and 1540 cm^-1^ emerged (See SI). Peracetylated sAFGPs were characterized by ^1^H NMR and size exclusion chromatography coupled to multi-angle light scattering and refractive index (SEC/MALS/RI) (Fig.2A, Table 1, see SI). MW and DP correlated well with expected values.

**Table 1.** Representative data for characterization of sAFGP polymers prepared using (PMe_3_)_4_Co in THF. [a] Sample name and amino acid composition. [b,c] M_n_, and Đ as determined by SEC/MALS/RI in DMF with 0.1M LiBr at 60 °C. All polymers were analyzed in their peracetylated forms. -indicates samples which were insoluble in the SEC mobile phase and were therefore analyzed by ^1^H NMR and Đ was not determined. [d] Observed degree of polymerization (DP).

| sAFGP<sup>[a]</sup> | $M_n$<sup>[b]</sup> | $\bar{D}$<sup>[b]</sup> | DP<sup>[c]</sup> |
| --- | --- | --- | --- |
| $(\beta\text{GalT}_{0.33}\text{-s-A}_{0.66})_n$ | 17,680 | - | 93 |
| $(\alpha\text{GalT}_{0.33}\text{-s-A}_{0.66})_n$ | 17,680 | - | 93 |
| $(\beta\text{GalNAcT}_{0.33}\text{-s-A}_{0.66})_n$ | 17,649 | - | 93 |
| $(\alpha\text{GalNAcT}_{0.33}\text{-s-A}_{0.66})_n$ | 17,649 | - | 93 |
| $(\beta\text{Gal}\alpha\text{GalNAcT}_{0.33}\text{-s-A}_{0.66})_n$ | 8,134 | 1.81 | 28 |
| $(\beta\text{Gal}\alpha\text{GalNAcT}_{0.33}\text{-s-A}_{0.66})_n$ | 16,460 | 1.27 | 57 |
| $(\beta\text{Gal}\alpha\text{GalNAcT}_{0.33}\text{-s-A}_{0.66})_n$ | 39,680 | 1.17 | 138 |
| $(\beta\text{Gal}\alpha\text{GalNAcT}_{0.33}\text{-s-A}_{0.66})_n$ | 48,830 | 1.39 | 170 |
| $(\beta\text{Gal}\alpha\text{GalNAcT}_{0.5}\text{-s-A}_{0.5})_n$ | 20,490 | 1.47 | 52 |
| $(\beta\text{Gal}\alpha\text{GalNAcT}_{0.66}\text{-s-A}_{0.33})_n$ | 23,140 | 1.31 | 46 |

### Characterization of sAFGP structure

Circular dichroism (CD) was used to characterize sAFGP secondary structures. Distinct signatures are observed for the η→π* and π →π* transitions of PPII, disordered, sheet, or α-helical conformations.^64–67^ Homopolymers of βGalNAc- or βGalαGalNAc-Thr have not been previously prepared; however, our previous work on αGalNAc-, αGal-, and βGal-bearing polyThr revealed they adopt extended PPII conformations.^47^ The αGalNAc amide positive ellipticity between 190–200 nm overlaps with the peptide π →π* transition.^47^

For sAFGPs with native 1:2 Thr:Ala and βGalαGalNAc, MW-dependent secondary structures were observed (Fig. 2B). The 28mer (equivalent to AFGP7–AFGP6), exhibited a combination of disordered and PPII conformations as evidenced by minor absorbance at ∼217 nm in the PPII region, and spectral alignment with denatured collagen and intrinsically disordered proteins.^66,67^ This was expected considering the 28mer can only make ∼9 PPII-helical turns. Helical propensity increases with chain length,^19,46^ which explains the classic PPII structure observed for 57- and 138-mers (maxima at ∼217 nm, minima at ∼197 nm). Our spectra correlate nicely with those of native AFGP5– AFGP2.^17,26,59^ Spectra obtained in water and phosphate buffered saline (PBS) were identical and sAFGP conformation was stable from 4–50°C (see SI).

**Figure 2.**
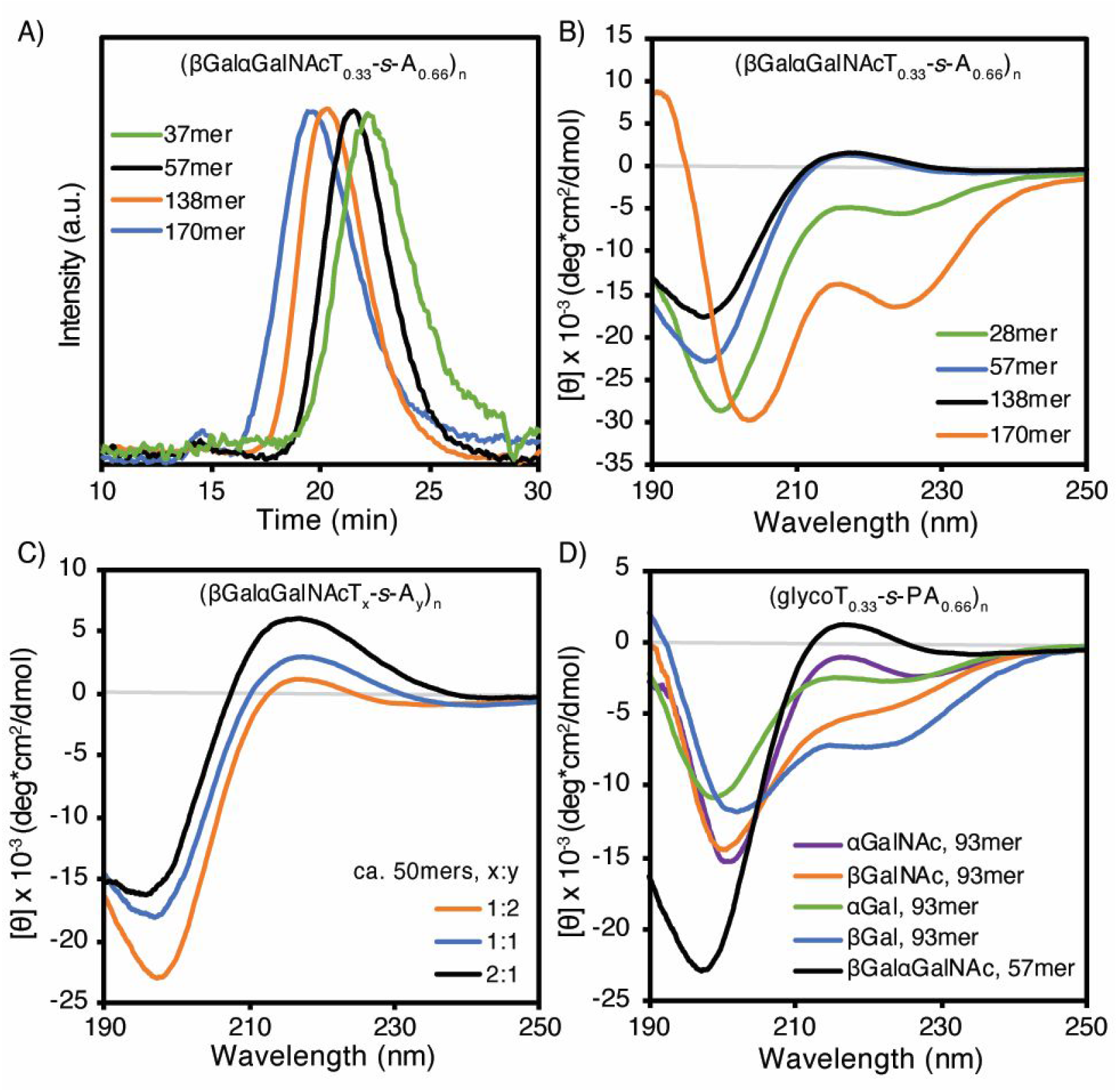
Characterization of sAFGP molar masses and conformations. A) SEC/MALS/RI in dimethylformamide with 0.1M LiBr, indicating differing elution times for peracetylated chains of increasing lengths. B–D) Aqueous CD spectra of deacetylated sAFGPs where B) are structures with increasing molecular weights, C) are structures with increasing βGalαGalNAcThr content, and D) are structures bearing glycans of differing identity and anomeric orientation.

The disaccharide-sAFGP 170mer adopted a mix of PPII-and presumably α-helices (minimum shift to 203nm, new minimum at 223nm), observed in multiple batches and spectral runs. PolyAla is a known α-helix former^68^, so we hypothesize larger MWs may facilitate Ala-rich microdomains. Though it is surprising the 32 additional residues could have this effect. However, compared to PPII, α-helical conformations have higher intensity absorbances at identical protein concentrations and therefore could be of low relative abundance. In any case, antifreeze activity was not affected (vide infra). Overall, we found that low MW sAFGPs are far less ordered than high MW AFGPs, correlating well with observations of native AFGPs.

Glycan identity and density also play a role in sAFGP conformation. Increasing βGalαGalNAcThr content, while decreasing Ala, led to proportional increases in extended PPII structure (maxima at ∼217nm, Fig. 2C). Clearly, βGalαGalNAc-Thr drives the PPII conformation. Truncation to αGalNAc in native 1:2 Thr:Ala structures resulted in reduced PPII propensity (reduced absorbance at 217 nm, Fig. 2D). αGal, βGal, and βGalNAc polymers adopt predominantly disordered conformations, indicated by the absence or reduction of the 217 nm band. These data suggest that the αGalNAc group partially orients the peptide into PPII helices, which is further stabilized by βGal(1→3) glycan extension.

### sAFGP antifreeze activity assays

Native AFGPs bind irreversibly to embryonic ice crystals, inhibiting prism-face growth and influencing overall shape and size. This results in a reduction in mean grain size (MGS), i.e. IRI, and shaping of single crystals into characteristic hexagonal and bipyramidal structures (Fig. 3A).^5^ We used cryostage microscopy and cooling splat assays to observe ice shaping and IRI activity for sAFGP ∼50mers with varied βGalαGalNAcThr:Ala ratios.^15^ Our controls were PBS and common cryoprotectant dimethylsulfoxide (DMSO).^69^

**Figure 3.**
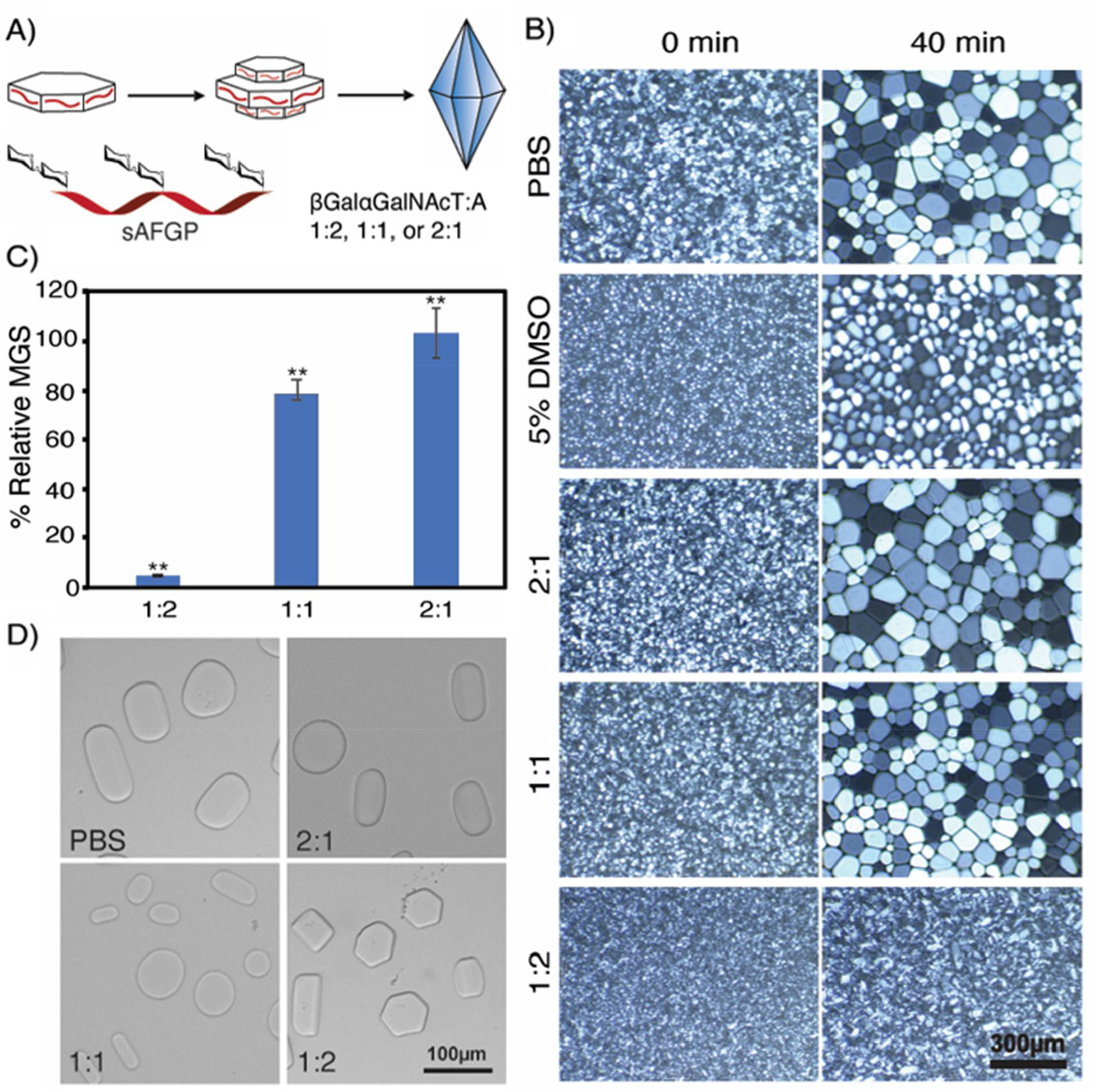
Ice binding properties of sAFGPs with varying amino acid compositions. A) Cartoon illustration of ice binding and shaping in the presence of sAFGPs composed of 1:2, 1:1, or 2:1 βGalαGalNAcThr:Ala ca. 50mers. B) Images of cooling splat assays and IRI activity for 71 µM sAFGP or 5 wt.% DMSO in PBS, or PBS alone. C) Quantified IRI data as % MGS relative to PBS; mean and standard deviation, ** indicates p < 0.01. D) Ice shaping experiments with 71 µM sAFGP in PBS.

Increasing Ala content correlated with higher IRI activity and stronger ice-shaping properties (Fig. 3B–D). Polymers with 33% Ala displayed no IRI activity and had no effect on crystal shape. Those with 50% Ala showed minor IRI activity (22% MGS reduction) but had little impact on crystal shape. By contrast, sAFGPs with native 66% Ala and 33% βGalαGalNAcThr exhibited remarkably potent IRI activity analogous to that of native AFGPs (95% MGS reduction).^5–8,44^ They also displayed identical hexagon-inducing ice-shaping properties as native AFGPs^5,6^. These findings highlight the importance of Ala for the ice-binding properties of (s)AFGPs.

IRI and ice-shaping also depended upon MW. Native ratio 1:2 βGalαGalNAcThr:Ala polymers 28, 57, or 170mers exhibited IRI activity that increases with chain length (Fig. 4A,B). Compared to PBS, the relative MGS reduction was 89% for sAFGP, 94% for 57mers, and 97% for 170mers. These data align with those of native AFGPs where AFGP1-5 (33.7–10.5kDa) show higher IRI than AFGP8 (2.65 kDa).^5,14^ Ice-shaping strength also increased with sAFGP chain length; 28mers resulted in amorphous, rounded crystals, while 57mers and 170mers produced angular crystals with hexagonal or rectangular morphologies (Fig. 4C).

**Fig. 4.**
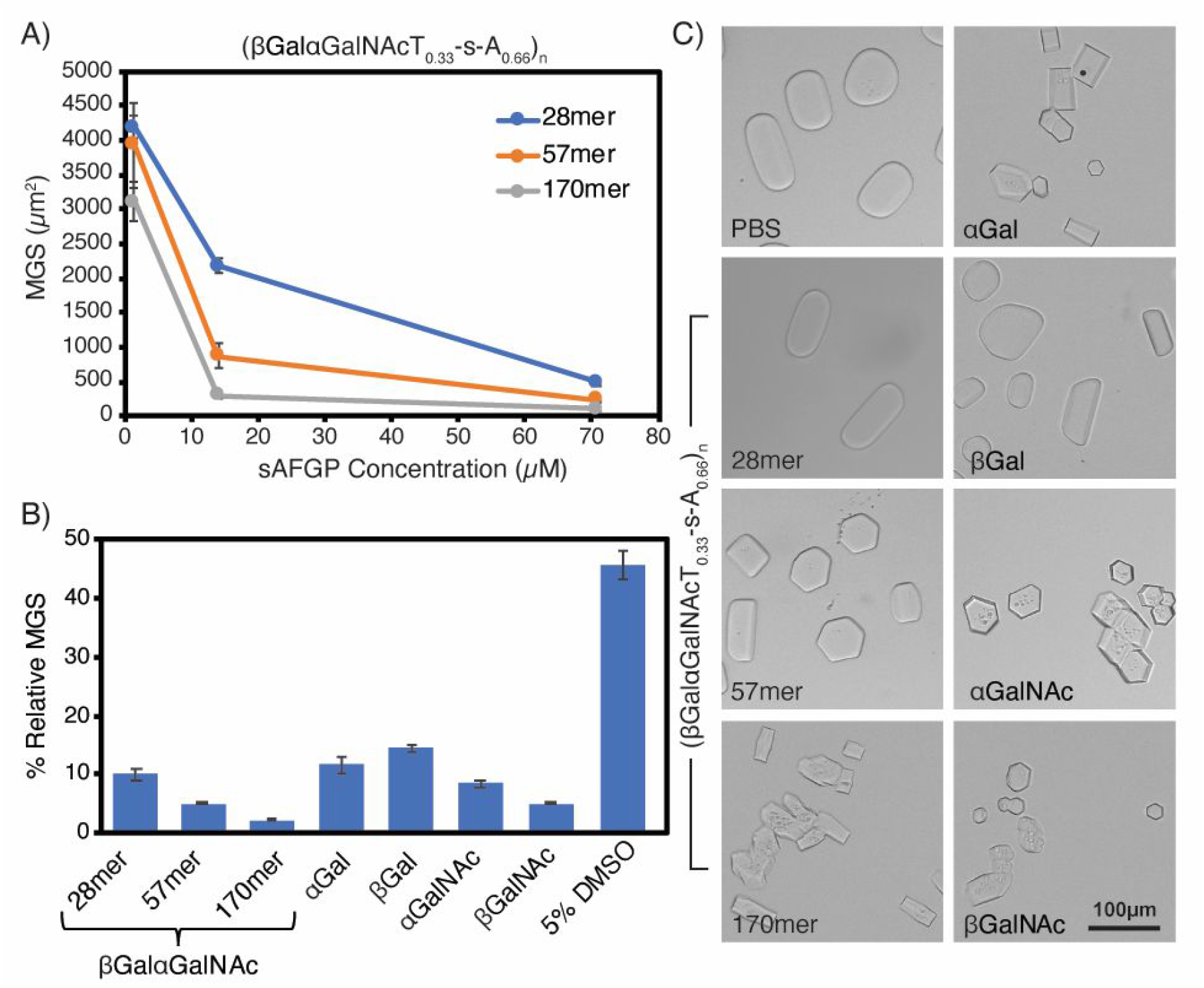
Ice binding data for sAFGPs composed of the native 1:2 glycoT:A ratio and with varied chain lengths and varied glycan structures. A) Observed absolute MGS at varied concentrations for sAFGPs with the native βGalαGalNAc disaccharide and with chain lengths of 28, 57, 170 residues. B) Quantified IRI data as % MGS relative to PBS for native disaccharide sAFGPs of varied chain lengths as compared to sAFGP 93mers bearing monosaccharides of varied structure and anomeric linkages, or 5% DMSO; sAFGP concentration is 71 µM in PBS; ice crystal MGS was determined from cooling splat assays; mean and standard deviation; table of statistical significance is in the SI. C) Ice shaping experiments for native disaccharide sAFGPs of varied chain lengths as compared to sAFGP 93mers bearing glycans of varied structure and anomeric linkages; sAFGP concentration is 71 µM in PBS.

Finally, we examined ice-shaping and IRI activity of sAFGPs with the native Thr:Ala ratio, moderate chain lengths (93mers), and varied glycans. All sAFGPs, regardless of glycan identity, exhibited strong IRI activity (Fig. 4B). Structures lacking NAc groups (αGal and βGal) had slightly lower activity than those with NAc (αGalNAc and βGalNAc) but remained potent antifreeze agents. Relative ice crystal MGSs from (αGalT_0.33_-*s*-A_0.66_)_93_ and (βGalT_0.33_-*s*-A_0.66_)_93_ solutions were reduced by 88% and 85%, respectively. (αGalNAcT_0.33_-*s*-A_0.66_)_93_ and (βGalNAcT_0.33_-*s*-A_0.66_)_93_ displayed higher activity, resulting in a relative MGS reduction of 92% and 97%. Comparing IRI activity on a mass basis yielded a similar trend (see SI). By contrast, 5% DMSO only achieved a 54% relative reduction. In a side-by-side trial, higher PVA concentrations were required to achieve a similar MGS reduction as our sAFGPs (see SI). Overall, our highest MW sAFGP of native composition, (βGalαGalNAcT_0.33_-s-A_0.66_)_170_, demonstrated the strongest IRI properties.

Ice-shaping patterns of glycan-variable sAFGPs aligned with IRI trends (Fig. 4C). αGal- or βGal-sAFGPs resulted in a mix of rounded and rectangular crystals, while αGalNAc- and βGalNAc-sAFGPs produced predominantly ordered, angular-faced structures. Intriguingly, (αGalNAcT_0.33_-*s*-A_0.66_)_93_ induced hexagonal crystal growth similar to native AFGPs^5,14^ and our disaccharide-sAFGP. These data suggest the αGalNAcThr plays a role in sAFGP ice-binding, either directly or indirectly by favorably orienting Ala or Thr methyls.

Our findings contradict the widely-cited study by Tachibana et al., which claimed the α-linkage, C2 NAc, and disaccharide were essential for antifreeze activity.^34^ By contrast, our structures with mono- and di-saccharides, α- and β-linkages, and with or without C2 NAc exhibit potent IRI activity. In the prior work, conclusions were drawn from structures of only ca. 6–9 amino acids that cannot adopt long-range conformations and have differing entropic considerations than macromolecular proteins. By contrast, our sAFGPs are comparable in size to native AFGPs and adopt ordered, stable PPII structures. Interestingly, our data also indicate the tripeptide repeat is unnecessary since our structures have statistically distributed residues.

### Cytocompatibility, cell internalization, and degradation of sAFGPs

In polar fish, AFGPs are present in the interstitial fluid of all body tissues except brain tissue, but intracellular accumulation has not been observed.^70^ Surprisingly, little is known regarding intracellular accumulation of AFGPs in the context of cryopreservation. A single report describes internalization of AFGP8 in human embryonic liver cells and trout gill cells.^71^ Additionally, limited evidence suggests cell membrane interactions.^72–74^ Therefore, we investigated the interaction of our sAFGPs with live human cells.

We labeled (βGalαGalNAcT_0.33_-*s*-A_0.66_)_57_ with AF594 using *N*-hydroxysuccinimide ester chemistry. We investigated internalization in human red blood cells (hRBCs), white blood cells (Raji), and embryonic kidney cells (HEK293), representing suspension and adherent models either of therapeutic value or to benchmark against the previous AFGP8 study. Following a 1-hour incubation with sAFGP-594 at 37°C, we observed substantial internalization and distribution within Raji and HEK cells, but not within hRBCs (Fig. 5A and SI). As hRBCs do not normally endocytose^75^, this suggests an endocytic uptake mechanism rather than passive translocation. Additionally, we examined uptake at endocytosis-inhibiting temperatures (4 °C, 23 °C), and minimal sAFGP was internalized (Fig. 5A). Work is underway to probe this phenomenon.

**Figure 5.**
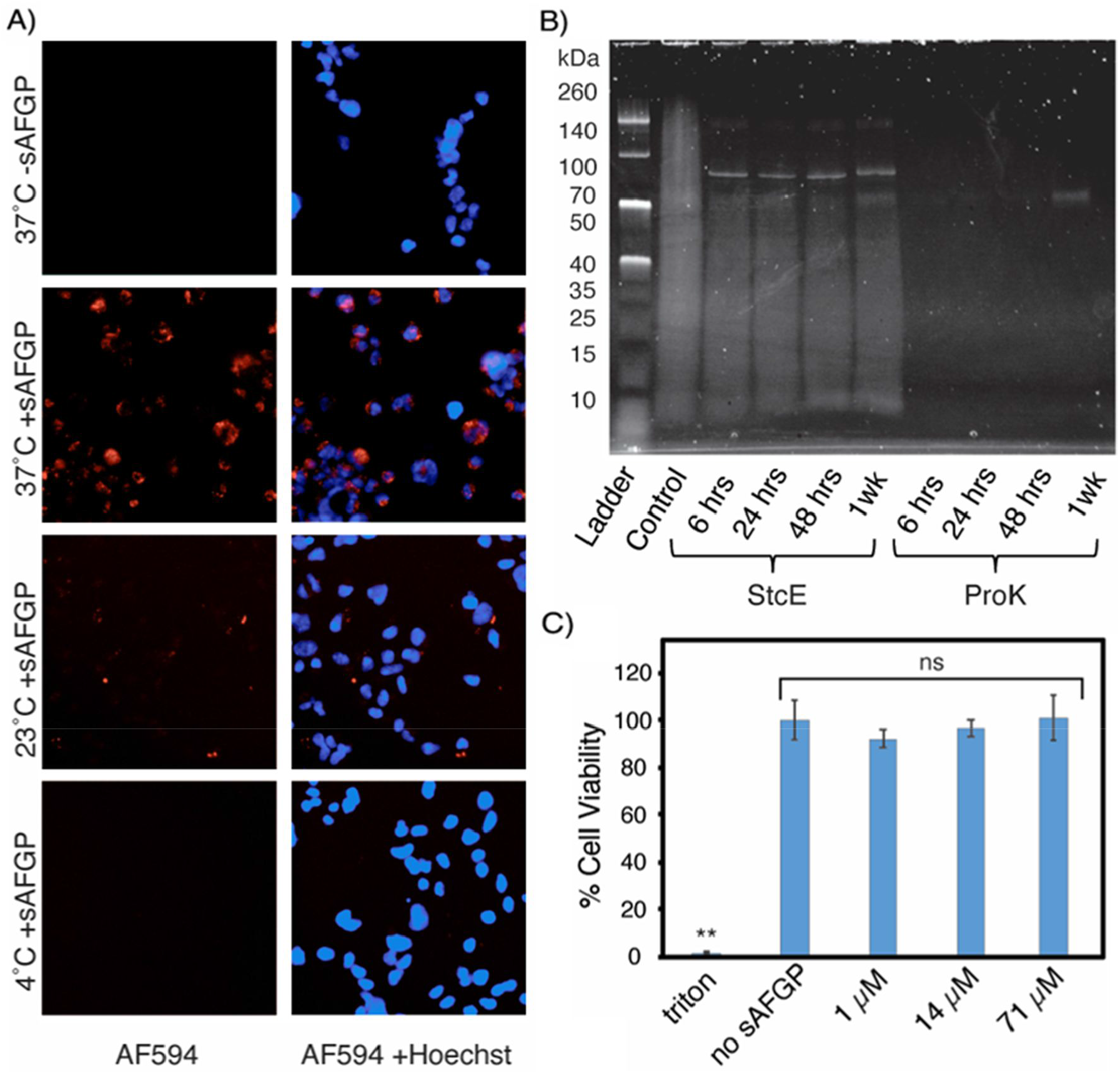
Cellular internalization, biodegradation, and cytocompatibility of sAFGPs with the native 1:2 glycoT:A composition and bearing the native disaccharide. A) Internalization of sAFGP 10 µM AF594-(βGalαGalNAcT_0.33_-s-A_0.66_)_57_ in HEK293 cells at 4, 23, or 37 °C. B) SDS-PAGE of protease-treated (βGalαGalNAcT_0.33_-s-A_0.66_)_98_ at varied timepoints, stained with glycoprotein-specific Pro-Q Emerald 300. C) HEK 293 cell viability as determined by CCK8 assay following 24-hour incubation with sAFGP (βGalαGalNAcT_0.33_-s-A_0.66_)_57_ at the indicated concentrations; no sAFGP is media alone as a negative control and Triton X-100 was a positive control; standard deviation; ** indicates p < 0.01.

Considering (s)AFGPs can be endocytosed, we investigated their potential for protease degradation. We treated moderately-sized sAFGP (βGalαGalNAcT_0.33_-s-A_0.66_)_98_ with non-specific proteinase K (ProK), which cleaves preferentially after hydrophobic sites^76^, and glycoprotease secreted protease of C1 esterase (StcE), which cleaves before αGalNAc-Ser/Thr^77^. ProK efficiently degraded the sAFGP within six hours while StcE only partially degraded the sample over one week (Fig. 5B); likely StcE prefers a monosaccharide substrate. Overall, sAFGPs are unlikely to bioaccumulate.

Toxic effects were unexpected since native AFGPs are present at up to 25 mg/mL in fish blood^1^ and similar synthetic glycopolypeptides are non-toxic^78^. Considering their validated internalization behavior, we assessed (βGalαGalNAcT_0.33_-s-A_0.66_)_57_ with HEK293 cells. CCK8 assays were conducted after 24 hours at 37°C with sAFGP concentrations relevant to IRI. At the highest concentration tested, there were no statistically significant effects on cellular viability (Fig. 5C), indicating the material’s suitability for cryopreservation applications.

### sAFGPs as Cellular Cryoprotectants

Cryopreservation is crucial for preserving cells and tissues for regenerative medicine and research, but requires use of cryoprotective agents (CPAs) to prevent ice crystal formation and cell damage.^12,13^ Little has changed in over 70 years^79^ and DMSO and glycerol are standard CPAs despite well-known toxic effects.^69,80^ Hydroxyethyl starch (HES) is a newer better-tolerated CPA, favorable due to membrane-impermanence, but still has associated toxic effects.^81^ Plus, excessive solution viscosity presents processing challenges. AFGPs have potential to revolutionize biomedical cryopreservation by replacing or reducing toxic CPAs. However, conflicting results and a lack of consensus on optimal cryopreservation conditions hinder their widespread use.^82,83^

For an initial study, we benchmarked our sAFGPs against a report utilizing native AFGPs combined with HES to freeze hRBCs.^72^ In their study, flash-freezing in liquid nitrogen with 130 mg/mL HES and slow thawing at ∼20°C resulted in 12% cell survival. Addition of 1– 800 µg/mL AFGP1–5 increased cell recovery up to 24%, but did not scale linearly and was optimized at 200 µg/mL. In our hands, hemolysis assays after flash-freezing of hRBCs with 130 mg/mL HES and 0–400 µg/mL (βGalαGalNAcT_0.33_-*s*-A_0.66_)_170_ followed by slow thaw at 23°C resulted in 3% cell recovery for PBS and 80% for HES alone, which hindered observation of any sAFGP effects (Fig. 6A, B). Therefore, we assessed hRBC hemolysis at varied HES concentrations and identified 40 mg/mL as having 20% post-thaw viability (see SI). Freeze-thaw experiments with 40 mg/mL HES supplemented with 0–400 µg/mL sAFGP showed no statistically significant effect of the sAFGPs on hRBC survival. Similar results were obtained for HEK293 cells (see SI).

**Figure 6.**
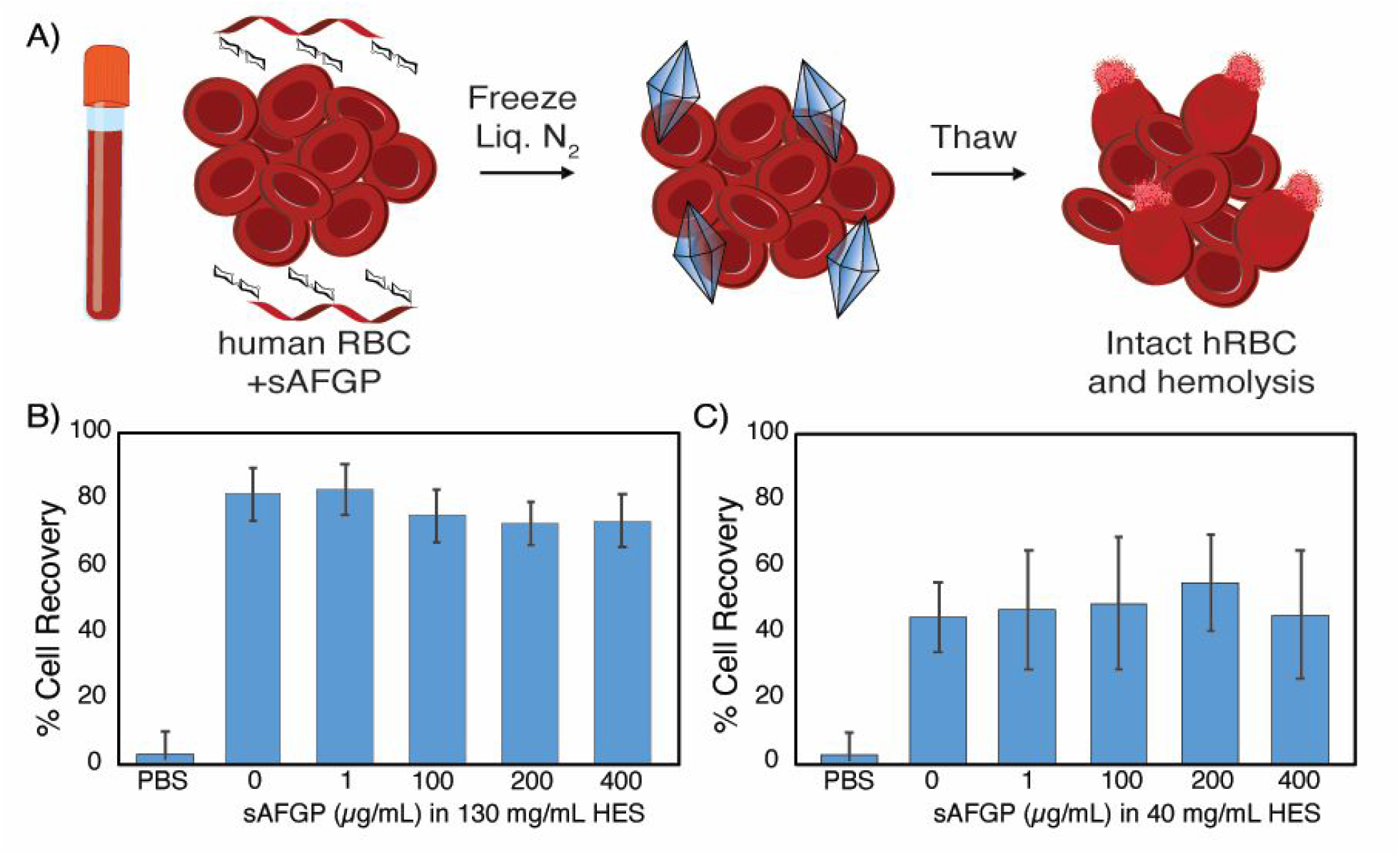
Cryopreservation of hRBCs in sAFGP supplemented HES solutions or PBS alone. A) Schematic representation of hRBC cryopreservation experimental workflow and cell hemolysis as the assessment metric of cell survival. B, C) Post-thaw intact hRBC cell recovery by hemolysis assays after freezing with sAFGP (βGalαGalNAcT_0.33_-s-A_0.66_)_170_ and B) 130mg/mL HES or C) 40mg/mL HES. Data are the result of two separate experiments performed in triplicate. Within each experiment, the cell recovery is not statistically different across the varied treatments.

This result was not surprising considering the modest effects of the native AFGP-HES combination in the prior study. In fact, in their work, freeze-thaw with AFGPs alone actually decreased cell survival as compared to PBS alone. Differences in our studies could be due to HES molecular weight or functionalization degree, which were not reported, varying thaw methodology, or the heterogeneity of native AFGPs and their challenging purification. Given the potent IRI and ice shaping activity of the sAFGPs, which are comparable to native AFGPs, further research on freezing methodology and CPA combinations are warranted and are underway.

## Conclusions

AFGPs have diverse applications in agriculture, food, surface coatings, and biomedical tissue cryopreservation. However, their isolation from polar organisms is cumbersome and impractical, hindering research on these unique molecules and their mechanisms of action. Here, we present a rapid and scalable method to synthetic AFGPs, allowing facile customization of molar mass and amino acid and glycan composition. We investigated a range of structures and found that hydrophobic Ala is essential for antifreeze activity, and potency increasing with molecular weight. While the native disaccharide displayed the highest potency, all glycan structures examined exhibited IRI properties. Our synthetic AFGPs endocytosed by human cell lines, are non-toxic and biodegradable, but did not alter red blood cell cryopreservation outcomes when used in combination with HES. In ice-binding studies, the synthetic AFGPs performed essentially identically to native AFGPs, indicating their promise as surrogates for these elusive natural structures.

## Methods

### General method for polymerization of NCAs

All polymerizations were prepared in a N_2_ filled glovebox. NCAs were dissolved in anhydrous THF at 50mg/mL in a glass vial or bomb tube. To the NCA solution, a 30mg/mL solution of (PMe_3_)_4_Co in THF was added and the tube was sealed. The NCA:(PMe3)4Co ratio ranged from 10:1 to 80:1, yielding different length polypeptides. The vials were left in the glove box at RT and the bomb tubes were removed from the gloved box and heated at 50°C for 5-72hrs. The reaction progress was monitored by attenuated total reflectance-Fourier transformed infrared spectroscopy (ATR-FTIR). Upon completion, the polypeptides were analyzed with SEC/MALS/RI.

### General method for polymerization of statistical copolymers

Copolymers were prepared in a N_2_ filled glovebox in a manner similar to homopolymers. The NCAs were dissolved in THF at 50mg/mL and mixed at a variety of NCA molar ratios. (PMe3)4Co catalyst in THF (30mg/mL) was added to the combined NCA solutions and the reaction progressed at RT and was monitored by ATR-FTIR. Polypeptides that remained soluble were analyzed using SEC/MALS/RI.

### General method for observing dynamic ice shaping

10uL of solution containing sAFGP in 1X PBS was placed on a microscope slide and sandwiched between a cover slip. The stage was rapidly cooled at a rate of 10°C/min to-30°C to freeze the solution. The stage was then slowly warmed to −2.5°C at a rate of 8°C/min. Then the stage was warmed to −1.8°C at a rate of 0.5°C/min. The stage temperature was then increased at a rate of 0.05°C/min to −1.5 to −1°C depending on the polypeptide solution to isolate individual crystals. The stage was then cooled at 0.02°C/min to −2 to −1.5°C to observe dynamic ice shaping. The stage was then toggled between melting and freezing rates to observe the ice crystal change as the temperature was increased and then decreased. Images of the single crystals were taken as the temperature was decreased to observe ice crystal growth.

### General method for cooling splat assays

Using a micropipette, 10µL of sAFGP in PBS was dropped from 2 meters through a PVC pipe onto a precooled glass slide (aluminum block resting in a bed of dry ice) at −78.5°C to form a thin wafer. The slide containing the ice splat was quickly moved to a temperature-controlled microscope stage (Linkam LTS120, WCP, and T96 controller) precooled to −6.4°C. See SI notes 1–3 for details. Typically, ice wafers are annealed at a temperature ranging from −6–8°C to ensure that a eutectic phase is present at the crystal boundary and the ice is able to undergo recrystallization.^84,85^ Use of PBS or a saline solution is essential due to the overestimation of IRI activity if observed in pure water.^84,85^, The ice wafers were annealed for 40 minutes at −6.4°C, and images of the ice crystals were recorded at 0, 20, and 40 minutes using cross polarizers (MOTICAM S3, MOTIC BA310E LED Trinocular) to observe ice recrystallization inhibition. The stage chamber was purged with N_2_ to prevent condensation from growing on the ice splat (See SI note 4). From the images, ice MGS was determined using image processing software or manual measurements. To ensure statistical significance, three images were obtained for each sample, and grain sizes within a minimum of three 150^2^ µm^2^ regions per sample were measured. Regions toward the center of the wafer, rather than near the edges, were selected.

### General method for glycopolypeptide cellular internalization studies with HEK293 cells

HEK 293 cells were plated on three 24 well plates and incubated at 37°C in 5% CO_2_ overnight to allow the cells to adhere. After 24 hours, the cell media was removed. A 100µM solution of A594-(βGalαGalNAcT_0.33_-s-A_0.66_)_57_ in MilliQ was diluted in complete media to make a 10uM solution of polypeptide. The solution was sterile-filtered before 300µL of the solution was applied to the cells. Additionally, 300µL of complete media was applied to additional wells of cells to serve as the untreated control. The 24 well plates were then placed at 37°C, RT, and 4°C to incubate for 1 hour. After one hour, the media was removed and the cells were rinsed 3x with DPBS. 500µL of Hoescht stain was then added to each well and cells were incubated for 10 minutes at RT. The cells were then fluorescently imaged to observe the localization of the fluorescent polymer.

### General method for freezing and thawing hRBCs for cryoprotection studies

The cryopreservation of hRBC was conducted following published procedures.^72^ In short, 50µL hRBCs (prepared as discussed in SI) were mixed with 50µL of DPBS containing cryoprotectants or control solution in cryovials. The vials were then placed in a liquid nitrogen bath for 20 minutes. The samples were then thawed at room temperature for 20 minutes. Control samples were also prepared according to this publication. hRBCs were added to water and frozen for 100% hemolysis and for 0% hemolysis, hRBCs were incubated with DPBS at room temperature for 1 hr. For all hRBC cryopreservation experiments, 2-hydroxyethyl starch (Spectrum Chemical, H3012) was used.

## TOC Graphic

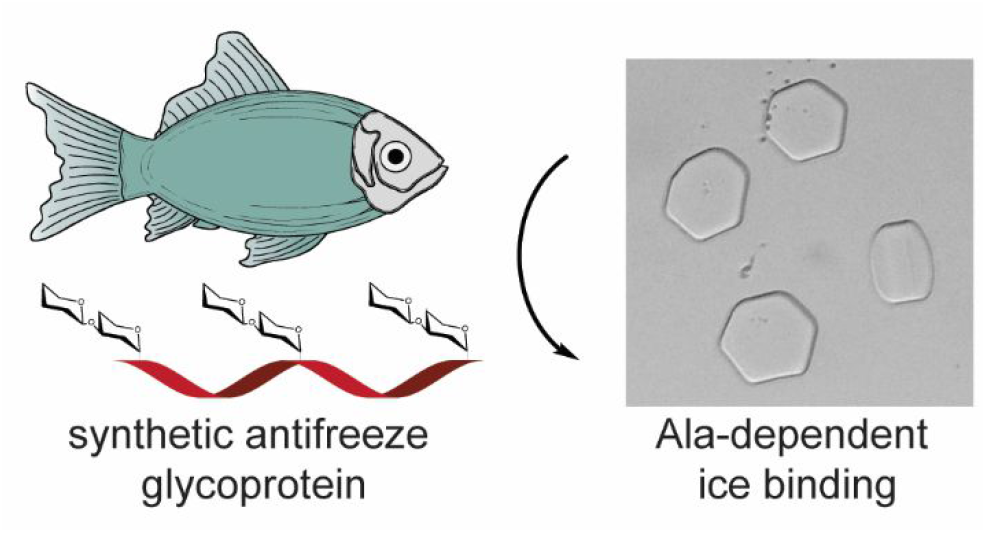

## Synopsis

Scalable, tunable polymer mimics of antifreeze glycoproteins found in polar fish bind and shape ice crystals in an Ala-dependent manner, offering new opportunities for cryoprotection.

## Notes

### Competing Interest Statement

The authors have declared no competing interest.

