## Supplemental info for "Synthetic antifreeze glycoproteins with potent ice-binding activity"

#### I. Instrumentation and General Methods

Reactions were conducted under an inert atmosphere of $N_2$, and anhydrous reactions were conducted using oven-dried glassware unless otherwise stated. All glassware was oven dried at 120°C. Hexanes and dichloromethane were purified by first purging with dry nitrogen, followed by passage through columns of activated 3Å molecular sieves. THF was purified by first purging with dry nitrogen, followed by passage through columns of activated alumina. Infrared spectra were recorded on a Bruker Alpha ATR-FTIR Spectrophotometer. All polymerizations were monitored for completion via ATR-FTIR. Deionized water (18 MΩ-cm) was obtained by passing in-house deionized water through a Thermo Scientific MicroPure UV/UF purification unit (MilliQ) resulting in AST grade 1 water (ultrapure water). Tandem size exclusion chromatography/refractive index (SEC/MALS/RI) was performed on an Agilent 1260 Infinity liquid chromatograph pump equipped with a Wyatt DAWN HELEOS-II light scattering (LS) and Wyatt Optilab T-rEX refractive index (RI) detectors. Separations were achieved using $10^5$, $10^4$, and $10^3$Å Phenomenex Phenogel 5 µm columns using 0.10 M LiBr in DMF as the eluent at 60 °C. All GPC/LS samples were prepared at concentrations of 3 mg/mL. Dn/dc values were calculated by batch injection of a series of polymer concentrations or by determining the degree of polymerization by $^1$H NMR of polyethyleneglycol (PEG) end-capped polymers and fitting to the observed RI data. The applied dn/dc for α/βGalOAc$_4$Thr, α/βGalNAcOAc$_3$Thr, and βGalαGalNAcThr copolymers were 0.0456, 0.0462, and 0.0723, respectively. CD measurements of the polypeptide solutions were recorded in quartz cells with a path length of 0.1 cm, on a JASCO J-1500 CD spectrophotometer. $^1$H and $^{13}$C NMR spectra were recorded on a Varian Mercury spectrometer (400 MHz) or a Bruker Avance NEO 500 (500 MHz) and are reported relative to deuterated solvent. Data for $^1$H NMR are reported as follows: chemical shift (δ ppm), multiplicity, coupling constant (Hz) and integration. Data for $^{13}$C NMR spectra are reported in chemical shift. Common solvent impurities found in spectra are labelled.[1]

#### II. Synthetic Procedures

##### II.a Glycosylated Amino Acid Preparation

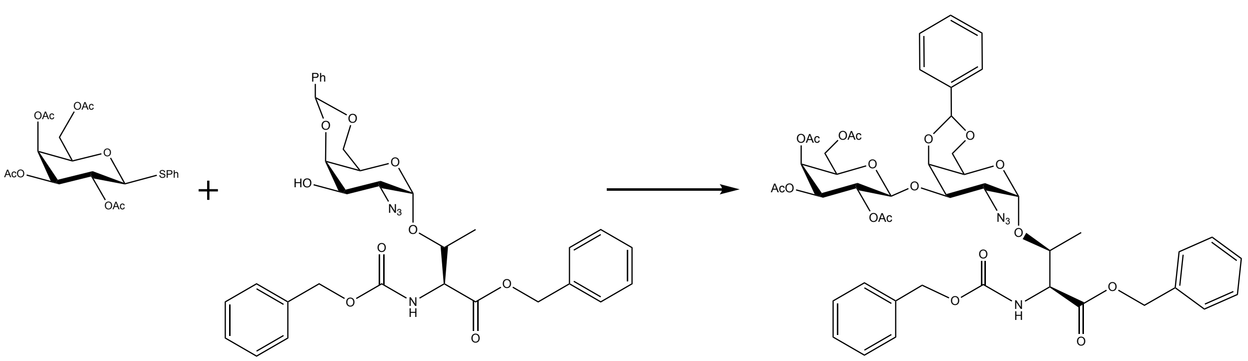

Synthetic scheme for the coupling of the second sugar to αGalN$_3$Thr.

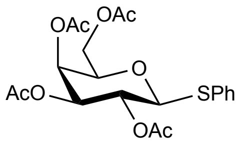

**1-Thiophenyl-2,3,4,6-acetyl-β-d-galactopyranoside, GalOAc$_4$-SPh** was synthesized using published methods.[2]

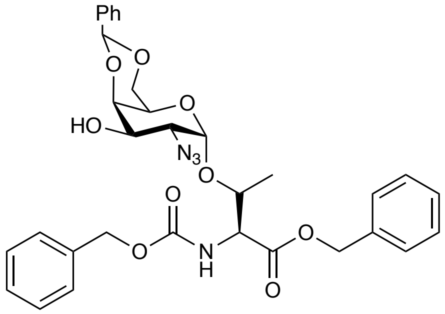

**N-α-(Carbobenzyloxy)-*O*-(2-azido-2-deoxy-4,6-benzylidene-α-D-galactopyranosyl)-L-threonine benzyl ester, Z-Thr(benzylidene-αGalN$_3$)-OBn** was synthesized using published methods.[3]

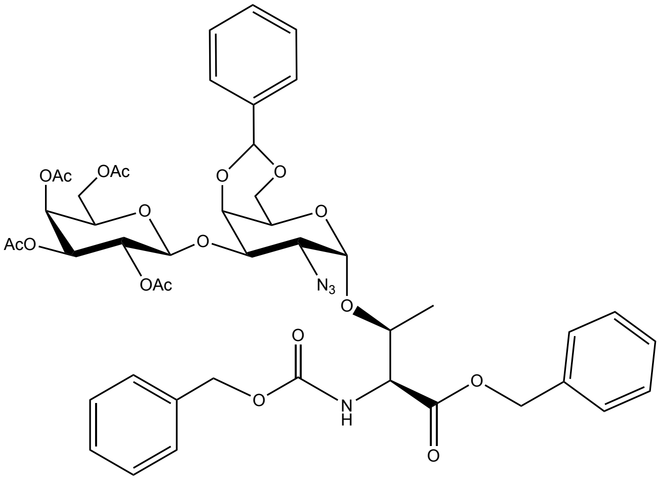

**N-α-(Carbobenzyloxy)-*O*-(2,3,4,6-tetra-*O*-acetyl-β-D-galactopyranosyl)-(1,3)-(2-azido-2-deoxy-4,6-benzylidene-α-D-galactopyranosyl)-L-threonine benzyl ester, Z-Thr(βGalOAc$_4$-(1,3)-benzylidene-αGalN$_3$)-OBn**

GalOAc$_4$-SPh and Z-Thr(benzylidene-αGalN$_3$)-OBn coupling was performed by modifying a published procedure.[4] Anhydrous DCM (0.53mL) and 3Å molecular sieves (0.17g) were added to a flask containing GalOAc$_4$-SPh (0.093g, 1.33 equiv) and Z-Thr-OBn (0.0982g, 1 equiv) and the mixture was cooled to -45˚C. Next, a solution of N-iodosuccinimide (0.143g, 4 equiv) in acetonitrile (0.63mL) was added followed by triflic acid (0.0056mL, 0.4 equiv). The reaction was stirred for 2 hours and reaction progress was monitored by TLC (2:1 hexanes:EtOAc, phosphomolybdic acid). Upon completion, the reaction was diluted with EtOAc and filtered through celite and cotton. The filtrate was washed 2x with aqueous 10% $Na_2S_2O_3$, then 2x with aqueous saturated $NaHCO_3$, and last 2x with aqueous saturated NaCl. The organic layer was dried over $Mg_2SO_4$, filtered, and concentrated. The crude product was purified using column chromatography (2:1 hexanes:EtOAc). The fractions were analyzed with TLC (2:1 hexanes:EtOAc, phosphomolybdic acid). Like fractions were combined and concentrated resulting in 0.122g (80% yield). $^1$H NMR (400 MHz, cdcl$_3$) δ 7.55 – 7.49 (m, 2H), 7.41 – 7.33 (m, 13H), 5.67 (d, *J* = 9.5 Hz, 1H), 5.52 (s, 1H), 5.41 (d, *J* = 3.5 Hz, 1H), 5.29 (dd, *J* = 10.4, 7.8 Hz, 1H), 5.21 (s, 2H), 5.16 (d, *J* = 2.7 Hz, 2H), 5.02 (dd, *J* = 10.4, 3.5 Hz, 1H), 4.92 (d, *J* = 3.6 Hz, 1H), 4.75 (d, *J* = 7.9 Hz, 1H), 4.50 – 4.41 (m, 2H), 4.34 (d, *J* = 3.2 Hz, 1H), 4.26 – 4.10 (m, 3H), 4.00 (d, *J* = 12.7 Hz, 1H), 3.97 – 3.89 (m, 2H), 3.75 (dd, *J* = 10.8, 3.6 Hz, 1H), 3.62 (s, 1H), 2.15 (s, 3H), 2.03 (s, 6H), 1.99 (s, 3H), 1.31 (d, *J* = 6.2 Hz, 3H).

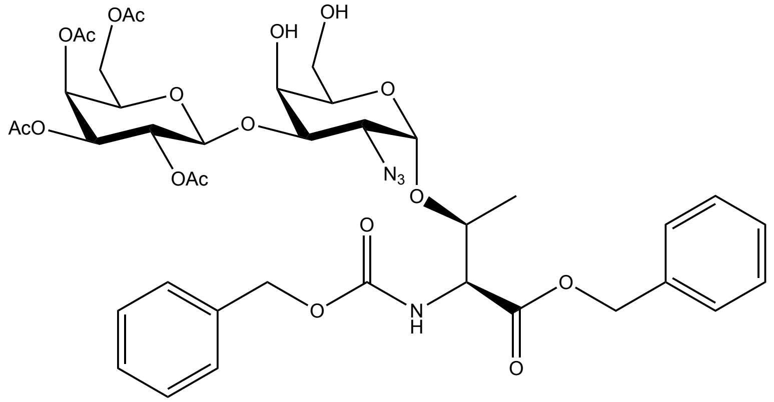

**N-α-(Carbobenzyloxy)-*O*-(2,3,4,6-tetra-*O*-acetyl-β-D-galactopyranosyl)-(1,3)-(2-azido-2-deoxy-α-D-galactopyranosyl)-L-threonine benzyl ester, Z-Thr(βGal(OAc)$_4$-(1,3)-αGalN$_3$)-OBn**

Acetic acid (28.9mL) and DI water (7.5mL) (4:1 acid:water) were added to a flask containing Z-Thr(βGal(OAc)$_4$-(1,3)-phenylisopropylidene-αGalN$_3$)-OBn (2.0g). The reaction was heated to 80˚C for 3 hours. The reaction was monitored by TLC (2:1 EtOAc:hexanes, phosphomolybdic acid). Upon completion, the reaction was concentrated with toluene to produce a fluffy white solid (1.81g, 91%). $^1$H NMR analysis showed complete removal of the benzylidene, and the product was used directly in the next reaction.

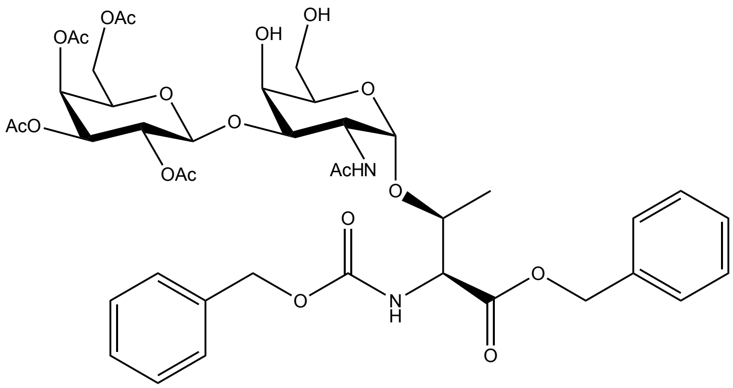

**N-α-(Carbobenzyloxy)-*O*-(2,3,4,6-tetra-*O*-acetyl-β-D-galactopyranosyl)-(1,3)-(2-acetamido-2-deoxy-α-D-galactopyranosyl)-L-threonine benzyl ester, Z-Thr(βGal(OAc)$_4$-(1,3)-αGalNAc)-OBn**

Z-Thr(βGal(OAc)$_4$-(1,3)-αGalN$_3$)-OBn (1.81g) was dissolved in THF (40mL), acetic acid (13.3mL), and acetic anhydride (26.6mL). Zinc powder (2.1g) was added to the solution and then saturated aqueous $CuSO_4$ solution (4mL) was added. The reaction was stirred at room temperature for 1 hour. The reaction was monitored by TLC (1.5:1 EtOAc:hexanes, phosphomolybdic acid). Upon completion, the reaction was filtered through celite and cotton, and the filtrate was concentrated with toluene to produce a chalky white solid which was used directly in the next reaction.

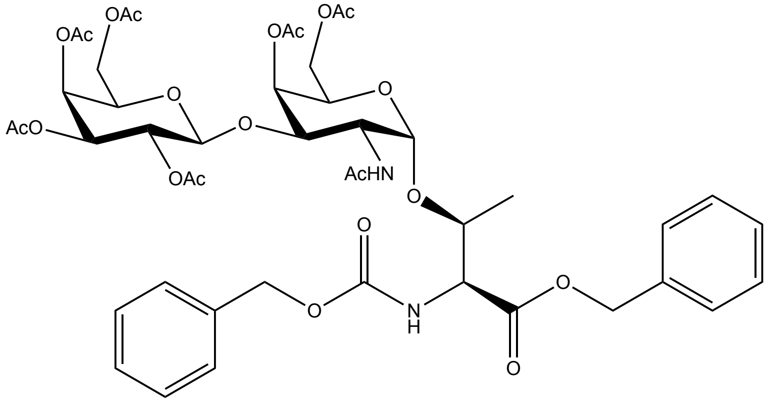

**N-α-(Carbobenzyloxy)-*O*-(2,3,4,6-tetra-*O*-acetyl-β-D-galactopyranosyl)-(1,3)-(2-acetamido-2-deoxy-4,6-*O*-acetyl-α-D-galactopyranosyl)-L-threonine benzyl ester, Z-Thr(βGal(OAc)$_4$-(1,3)-αGalNAc(OAc)$_3$)-OBn**

To acetylate the free hydroxyls, Z-Thr(βGal(OAc)$_4$-(1,3)-αGalNAc)-OBn (0.085g) was cooled to 0˚C and dissolved in acetic anhydride (0.67mL) and pyridine (1.33mL). The reaction was stirred overnight and allowed to warm to room temperature. The reaction was monitored by TLC (3:1

EtOAc:hexanes, phosphomolybdic acid). Upon completion, the reaction was cooled to 0˚C and quenched with 5mL DI water. The aqueous layer was extracted 3x with EtOAc. The organic layer was then washed 2x with 1M HCl, then 2x with aqueous saturated $NaHCO_3$, and lastly 2x with aqueous saturated NaCl, and dried over $Na_2SO_4$. The crude product was purified using column chromatography (1:3 hexanes:EtOAc). The fractions were analyzed with TLC (1:3 hexanes:EtOAc, phosphomolybdic acid). Like fractions were combined and concentrated resulting in 0.059g (63% yield). $^1$H NMR (500 MHz, $CDCl_3$) δ 7.46 – 7.34 (m, 10H), 7.33 (s, 1H), 5.71 (d, *J* = 8.9 Hz, 1H), 5.48 (d, *J* = 9.4 Hz, 1H), 5.37 (dd, *J* = 7.4, 3.3 Hz, 2H), 5.23 (d, *J* = 11.8 Hz, 1H), 5.16 – 5.07 (m, 2H), 4.97 (dd, *J* = 10.6, 3.4 Hz, 1H), 4.83 (d, *J* = 3.8 Hz, 1H), 4.59 (d, *J* = 7.8 Hz, 1H), 4.46 (s, 2H), 4.24 (d, *J* = 6.4 Hz, 1H), 4.17 (d, *J* = 5.1 Hz, 3H), 4.13 – 4.08 (m, 2H), 3.98 (dd, *J* = 11.3, 7.4 Hz, 1H), 3.91 (s, 1H), 3.80 (dd, *J* = 11.0, 3.2 Hz, 1H), 2.15 (d, *J* = 4.5 Hz, 6H), 2.09 (s, 3H), 2.05 (d, *J* = 1.7 Hz, 9H), 1.99 (s, 3H), 1.35 (d, *J* = 6.3 Hz, 3H).

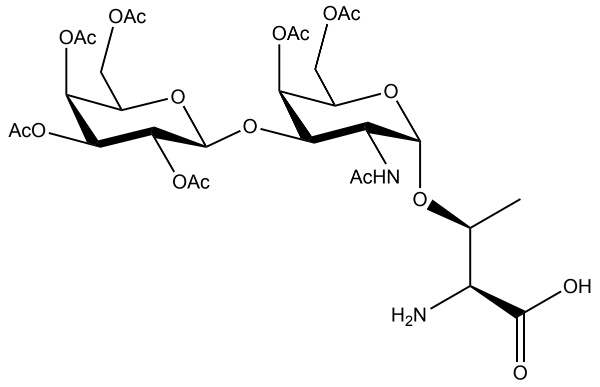

***O*-(2,3,4,6-tetra-*O*-acetyl-β-D-galactopyranosyl)-(1,3)-(2-acetamido-2-deoxy-4,6-*O*-acetyl-α-D-galactopyranosyl)-L-threonine, H-Thr(βGal(OAc)$_4$-(1,3)-αGalNAc(OAc)$_3$)-OH**

Z-Thr(βGal(OAc)$_4$-(1,3)-αGalNAc(OAc)$_3$)-OBn 0.6852g, 1equiv) was dissolved in MeOH (28,5mL, 0.025M) and added to a flask containing 10% Pd/C (0.137g, 20% starting material mass) under $H_2$. The reaction was stirred vigorously for 3 hours. The reaction progress was monitor by TLC (1:3 hexanes:EtOAc + 1% acetic acid, phosphomolybdic acid). After 3 hours, the reaction was filtered through cotton and a 0.45μm filter to remove the Pd/C. The reaction was then concentrated, resulting in a white solid. $^1$H NMR analysis confirmed the disappearance of aromatic protons belonging to the CBz and Bn protecting groups. The product was used directly in the next reaction without further purification.

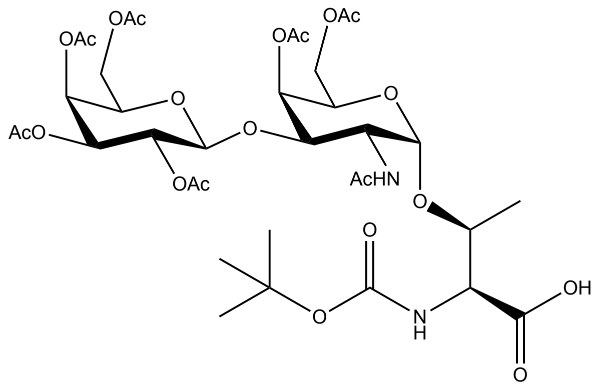

**N-α-(butoxycarbonyl)-*O*-(2,3,4,6-tetra-*O*-acetyl-β-D-galactopyranosyl)-(1,3)-(2-acetamido-2-deoxy-4,6-*O*-acetyl-α-D-galactopyranosyl)-L-threonine, Boc-Thr(βGal(OAc)$_4$-(1,3)-αGalNAc(OAc)$_3$)-OH**

H-Thr(βGal(OAc)$_4$-(1,3)-αGalNAc(OAc)$_3$)-OH (0.525g, 1 equiv) was added to a 1:1 mixture of THF/water (7mL) and cooled to 0˚C. In a modification to publish procedures[5], $NaHCO_3$ (0.749g, 12.5 equiv) and $BocO_2$ (0.778g, 5 equiv) were added consecutively at increased molar equivalents. The reaction was allowed to warm to RT and stirred overnight. The reaction progress was monitored by TLC (1:3 hexanes:EtOAc + 1% acetic acid, phosphomolybdic acid). Upon completion, the reaction was washed 3x with $Et_2O$ to remove excess $BocO_2$. The aqueous layer was cooled to 0˚C and acidified to pH 4-5 using acetic acid. The aqueous layer was extracted 3x with DCM and the combined organic layers dried over sodium sulfate. The organic layers were concentrated yielding a white solid. The crude product was purified using column chromatography (eluent 1:3 hexanes:EtOAc +1% acetic acid). Fractions were analyzed by TLC (1:3 hexanes:EtOAc + 1% acetic acid, phosphomolybdic acid) and combined to produce a white, fluffy solid (0.4935g, 83% yield over two steps). $^1$H NMR (400 MHz, $cd_3od$) δ 5.41 (d, *J* = 3.3 Hz, 1H), 5.35 (d, *J* = 3.3 Hz, 1H), 5.05 (dd, *J* = 10.5, 3.3 Hz, 1H), 5.02 – 4.96 (m, 1H), 4.89 (d, *J* = 3.8 Hz, 1H), 4.75 (d, *J* = 7.6 Hz, 1H), 4.34 (dd, *J* = 11.3, 4.0 Hz, 2H), 4.25 (dd, *J* = 8.0, 4.5 Hz, 1H), 4.19 – 4.09 (m, 4H), 4.06 (q, *J* = 6.8 Hz, 1H), 3.98 (td, *J* = 9.8, 5.4 Hz, 2H), 2.11 (d, *J* = 10.5 Hz, 7H), 2.03 (t, *J* = 4.1 Hz, 12H), 1.93 (s, 3H), 1.48 (s, 9H), 1.28 (s, 4H). 13C NMR (126 MHz, CDCl3) δ 173.31, 172.82, 170.52, 170.49, 170.39, 170.33, 170.15, 170.01, 169.71, 169.37, 156.17, 155.99, 129.05, 128.24, 100.41, 99.74, 80.28, 77.58, 70.90, 70.71, 68.80, 67.75, 66.82, 63.21, 62.85, 61.13, 58.20, 53.22, 49.13, 28.46, 28.38, 22.82, 20.71, 20.56, 18.33.

**II.b NCA Preparation**

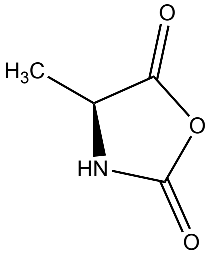

**Ala NCA** was synthesized using published methods.[6] $^{1}H$ NMR (500 MHz, $CDCl_3$) δ 5.72 (s, 1H), 4.41 (q, *J* = 7.0 Hz, 1H), 1.58 (d, *J* = 7.0 Hz, 3H). $^{13}C$ NMR (126 MHz, $CDCl_3$) δ 169.98, 151.65, 53.40, 17.95

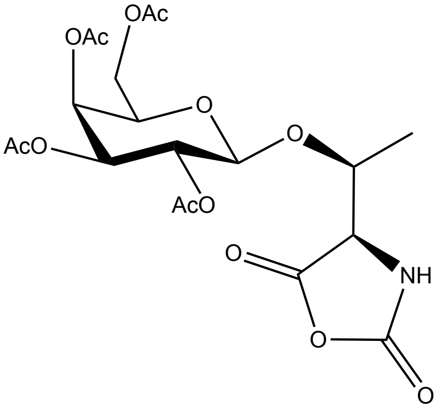

***O*-((2,3,4,6-tetra-O-acetyl)-β-D-galactopyranose))-L-threonine-N-carboxyanhydride, βGal(OAc)$_4$Thr NCA** was synthesized according to a published method.[3]

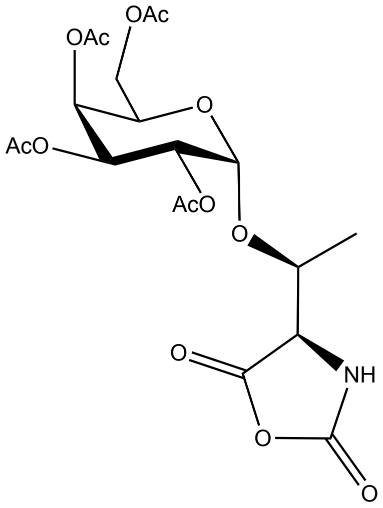

***O*-((2,3,4,6-tetra-O-acetyl)-α-D-galactopyranose)-L-threonine-N-carboxyanhydride, αGal(OAc)$_4$Thr NCA** was synthesized according to a published method.[3]

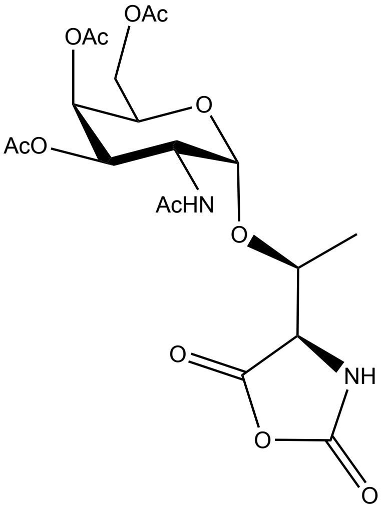

***O*-(2-acetamido-2-deoxy-3,4,6-tetra-O-acetyl-α-D-galactopyranosyl)-L-threonine-N-carboxyanhydride, αGalNAc(OAc)$_3$Thr NCA** was synthesized according to a published method.[3]

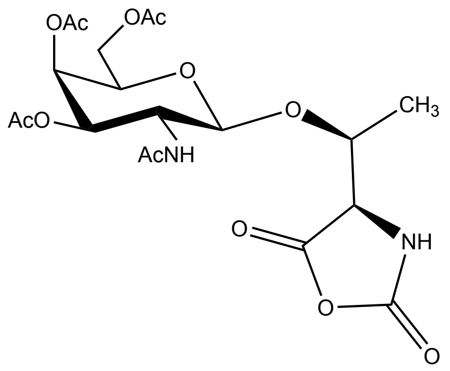

**O-((2-acetamido-2-deoxy-3,4,6-tri-O-acetyl)-β-D-galactopyranose)-L-threonine-N-carboxyanhydride, βGalNAc(OAc)$_4$Thr NCA** The cyclization of Boc-βGalNAc(OAc)$_3$Thr was completed by modifying publish methods.[3] The Boc-βGalNAc(OAc)$_3$Thr (0.1959g, 1equiv) was dissolved in anhydrous THF (3.57mL, 0.1M). Triphosgene (0.042g, 0.4 equiv) was added and the solution was cooled to 0˚C. Distilled TEA (0.0546mL, 1.1equiv) was added dropwise. The reaction was stirred for 3 hours and reaction progress was monitored by ATR-FTIR. After 3 hours the reaction was filtered through cotton to remove TEA-HCl salts. The reaction solution was evaporated under reduced pressure. The evaporate was sequestered in a tandem solvent trap system cooled by liquid $N_2$. The traps were immediately quenched with ammonium hydroxide. The crude product was purified using anhydrous silica chromatography[7] with 10% to 20% THF in DCM. The collected fractions were analyzed by ATR-FTIR. NCA-containing fractions were combined resulting in 0.089g of white solid (52% yield). $^1$H NMR (500 MHz, $CDCl_3$) δ 7.20 (s, 1H), 6.18 (d, $J$ = 8.6 Hz, 1H), 5.33 (d, $J$ = 3.5 Hz, 1H), 5.24 (dd, $J$ = 11.2, 3.6 Hz, 1H), 4.79 (d, $J$ = 8.3 Hz, 1H), 4.20 (d, $J$ = 5.9 Hz, 1H), 4.19 – 4.06 (m, 3H), 4.00 – 3.87 (m, 2H), 2.13 (d, $J$ = 4.9 Hz, 3H), 2.06 (d, $J$ = 4.3 Hz, 3H), 1.96 (dd, $J$ = 15.9, 5.0 Hz, 6H), 1.37 (d, $J$ = 6.3 Hz, 3H).$^{13}$C NMR (126 MHz, $CDCl_3$) δ 171.07, 171.01, 170.69, 170.33, 167.50, 100.61, 76.23, 71.24, 69.63, 68.06, 66.85, 62.59, 61.96, 51.58, 25.71, 23.38, 20.77, 17.39.

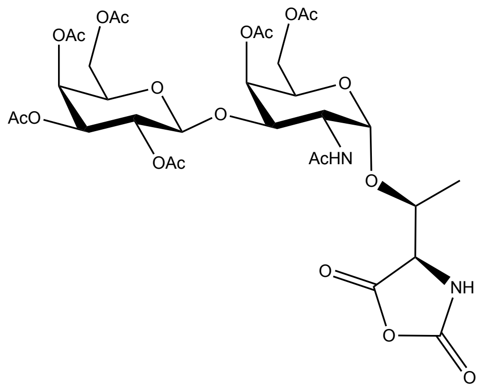

**N-α-(butoxycarbonyl)-*O*-(2,3,4,6-tetra-*O*-acetyl-β-D-galactopyranosyl)-(1,3)-(2-acetamido-2-deoxy-4,6-*O*-acetyl-α-D-galactopyranosyl)-L-threonine-*N*-carboxyanhydride, Thr(βGal(OAc)$_4$-(1,3)-αGalNAc(OAc)$_3$) NCA**

The cyclization of Boc-Thr(βGal(OAc)$_4$-(1,3)-αGalNAc(OAc)$_3$)-OH was completed by modifying published methods[3]. The Boc-Thr(βGal(OAc)$_4$-(1,3)-αGalNAc(OAc)$_3$)-OH (0.3373g, 1equiv) was dissolved in anhydrous THF (4.03mL, 0.1M). Triphosgene (0.048g, 0.4 equiv) was added and the solution was cooled to 0˚C. Distilled TEA (0.0448mL, 1.1equiv) was added dropwise. The reaction was stirred for 3 hours and reaction progress was monitored by ATR-FTIR. After 3 hours the reaction was filtered through cotton to remove TEA-HCl salts. The reaction solution was evaporated under reduced pressure. The evaporate was sequestered in a tandem solvent trap system cooled by liquid $N_2$. The traps were immediately quenched with ammonium hydroxide. The crude product was purified using anhydrous silica chromatography[7] with 10% to 30% to 50% THF in DCM. The collected fractions were analyzed by ATR-FTIR. NCA containing fractions were combined resulting in 0.2232g of white solid (72% yield). $^1$H NMR (500 MHz, CDCl3) δ 7.34 (s, 1H), 5.97 (d, J = 7.4 Hz, 1H), 5.35 (d, J = 3.1 Hz, 2H), 5.19 (d, J = 3.6 Hz, 1H), 5.12 (dd, J = 10.4, 7.9 Hz, 1H), 4.97 (dd, J = 10.4, 3.5 Hz, 1H), 4.65 (d, J = 8.0 Hz, 1H), 4.39 – 4.31 (m, 1H), 4.31 (s, 1H), 4.23 – 4.10 (m, 5H), 4.00 – 3.88 (m, 3H), 2.17 (s, 3H), 2.09 (d, J = 10.4 Hz, 6H), 2.05 (d, J = 2.6 Hz, 9H), 1.96 (s, 3H), 1.41 (d, J = 6.5 Hz, 3H).$^{13}$C NMR (126 MHz, CDCl3) δ 171.30, 170.86, 170.69, 170.50, 170.22, 170.20, 168.25, 152.45, 100.18, 99.44, 75.31, 72.23, 71.26, 70.98, 68.59, 68.53, 66.87, 62.86, 62.77, 61.25, 49.59, 23.18, 20.94, 20.88, 20.84, 20.82, 20.80, 20.64, 17.72.

**II.c sAFGP Preparation**

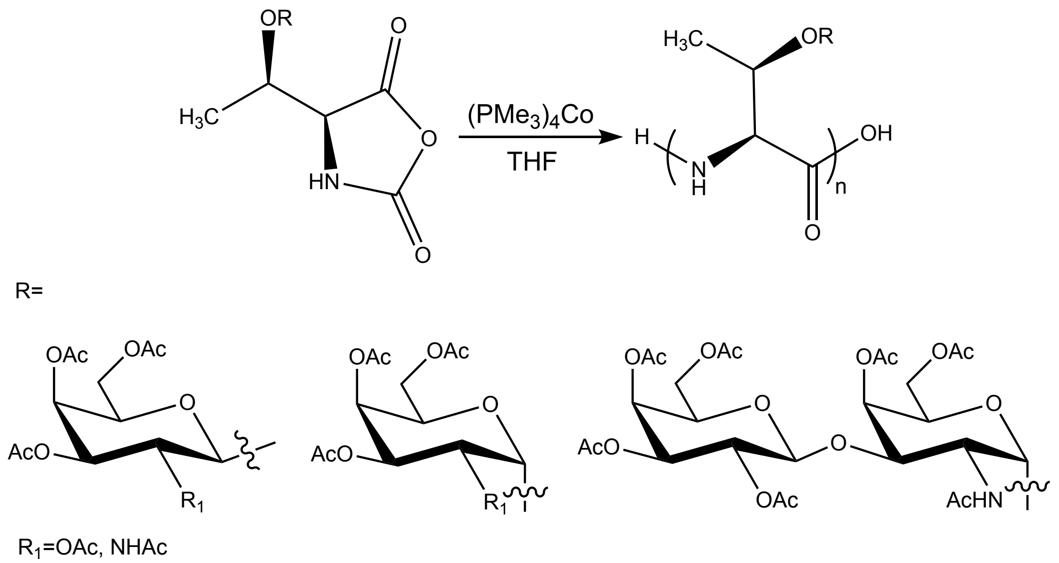

#### General method for polymerization of NCAs

All polymerizations were prepared in a $N_2$ filled glovebox. NCAs were dissolved in anhydrous THF at 50mg/mL in a glass vial or bomb tube. To the NCA solution, a 30mg/mL solution of $(PMe_3)_4Co$ in THF was added and the tube was sealed. The NCA:$(PMe_3)_4Co$ ratio ranged from 10:1 to 80:1, yielding different length polypeptides. The vials were left in the glove box at RT and the bomb tubes were removed from the gloved box and heated at 50˚C for 5-72hrs. The reaction progress was monitored by ATR-FTIR. Upon completion, the polypeptides were analyzed with SEC/MALS/RI.

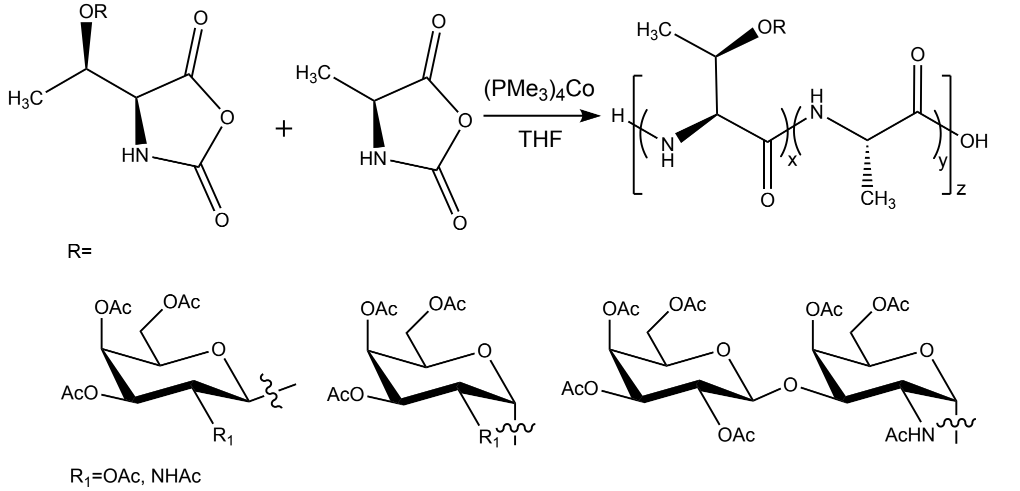

#### General method for polymerization of statistical copolymers

Copolymers were prepared in a $N_2$ filled glovebox in a manner similar to homopolymers. The NCAs were dissolved in THF at 50mg/mL and mixed at a variety of NCA molar ratios. $(PMe_3)_4Co$

catalyst in THF (30mg/mL) was added to the combined NCA solutions and the reaction progressed at RT and was monitored by ATR-FTIR. Polypeptides that remained soluble were analyzed using SEC/MALS/RI.

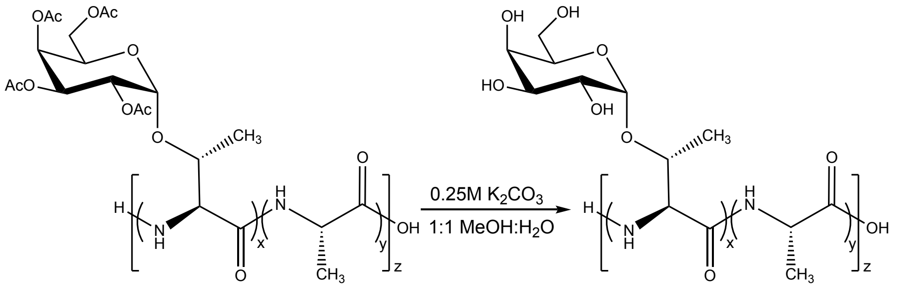

**General method for the deacetylation of AcO-protected polymers**

The polypeptide was suspended in 0.25M $K_2CO_3$ in 1:1 MeOH:$H_2O$ and stirred overnight. The deacetylated polymer in solution was transferred to a 1kDa spin filter and concentrated at 4000 x g for 20 minutes. The concentrate was diluted three times with ultrapure water and spin filtered at 4000 x g for 20 minutes each time. The concentrate was recovered, frozen, and lyophilized. Samples can also be dialyzed against ultrapure water in 2000 MWCO dialysis tubing.

**General method for labeling glycopolypeptides with AF594 fluorophore**

The deacetylated polypeptide was added to a 100µM solution of sodium bicarbonate in ultrapure water for a final polymer concentration of 5 mg/mL. AF594 was dissolved in DMSO at 10 mg/mL. Then 8 molar equivalents of AF594 were added to the polymer sodium bicarbonate solution. The tube containing the solution was wrapped in foil and placed on a shaker plate to react overnight. The solution was transferred to a 1kDa spin filter and concentrated at 4000 x g for 20 minutes. The concentrate was diluted three times with ultrapure water and spin filtered at 4000 x g for 20 minutes each time. The concentrate was recovered, frozen, and lyophilized. Samples can also be dialyzed against ultrapure water in 2000 MWCO dialysis tubing. The labeled glycopolypeptide was stored at -20˚C and shielded from light.

**III. General methods for sAFGP biological and ice binding assays**

**General method for protease digestion of glycopolypeptides**

Polypeptide is added to ultrapure water to make a 10 µg/µL stock solution. Polypeptide stock solution, 2X PBS, and protease (StcE or Proteinase K) were added to a reaction tube such that the protease to substrate ratio is 0.1. StcE was a gift from the lab of Carolyn Bertozzi and was expressed and purified according to literature.[8] Protease K was obtained from ThermoFisher (#AM2542). The solution is balanced with ultrapure water so that the final concentration of polymer is 1 μg/μL in 1X PBS. The solution was placed in a 37˚C water bath and 10uL timepoints were removed at 6, 24, 28 hours and after 7 days. Upon removal from the reaction vessel, timepoints are treated at 95˚C for 10 minutes to stop the protease activity. Samples are stored in -80˚C until analysis with electrophoresis.

**General method for electrophoresis and SDS-Page**

Polypeptide (20 µg) (with or without denatured protease) in PBS was combined with 4X loading buffer from BioRad to make a 1X solution of dye and polypeptide. Thermo Fisher Spectra Multicolor Broad Range Protein Ladder was applied to a separate well. The entire volume was applied to a well in a Bis-tris 4-12% gel from BioRad. The gel ran for 40 minutes at 175 V. Pro-Q ® Emerald 300 Lipopolysaccharide Gel Stain Kit from ThermoFisher was used to stain the glycopolypeptide. Modifications to the established protocol include staining gel for 100 minutes and creating fresh oxidizer from periodic acid for each gel. After completing the staining process gels were imaged using a standard gel imager and exposed for 4.2 seconds. When establishing a protocol for staining SDS-Page gels containing our polypeptides, we also tested Coomassie Blue R-250 and G-250, Gel Code Blue, and silver stain. None of these stains worked with our glycopolypeptides.

**General method for observing dynamic ice shaping**

To observe dynamic ice shaping 10uL of solution containing polypeptide in 1X PBS was placed on a microscope slide and sandwiched between a cover slip. The stage was rapidly cooled at a rate of 10˚C/min to -30˚C to freeze the polymer solution. The stage was then slowly warmed to -2.5˚C at a rate of 8˚C/min. Then the stage was warmed to -1.8˚C at a rate of 0.5˚C/min. The stage temperature was then increased at a rate of 0.05˚C/min to -1.5 to -1˚C depending on the polypeptide solution to isolate individual crystals. The stage was then cooled at 0.02˚C/min to -2 to -1.5˚C to observe dynamic ice shaping. The stage was then toggled between melting and freezing rates to observe the ice crystal change as the temperature was increased and then decreased. Images of the single crystals were taken as the temperature was decreased to observe ice crystal growth.

**General method for cooling splat assays**

To make the ice wafer, 10µL of solution containing polypeptide in PBS was dropped from 2 meters through a PVC pipe onto a precooled slide using a micropipette (see picture below) (Note 1 and

2). The slide was cooled on an aluminum block resting in a bed of dry ice (see picture below) (Note 3) . The slide containing the ice splat was quickly moved to the temperature controlled stage (Linkam LTS120, WCP, and T96 controller) precooled to -6.4˚C. The stage chamber was purged with $N_2$ to prevent condensation from growing on the ice splat (Note 4). The ice splat was annealed for 40 minutes and images of the ice crystals were recorded at 0, 20, and 40 minutes using cross polarizers (MOTICAM S3, MOTIC BA310E LED Trinocular) to observe ice recrystallization inhibition.

Notes:

1. To remove the static from the PVC pipe post purchase, unscented dryer sheets were pushed through the pipe and brushed alone the pipe openings.
2. With some polymer solutions, the surface tension made releasing the drop from the pipette difficult. To overcome this problem the outside of the pipette tip was wiped with immersion oil. Excess oil was removed with a clean wipe before dispensing solution.
3. It is important to have the slide positioned close to the opening of the PVC pipe so air flow does not alter the drop’s trajectory.
4. The flowrate of $N_2$ into the cryostage was reduced to < 0.1L/min after the stage was purged with $N_2$ to remove the ambient air introduced when placing the slide on the temperature controlled stage. Reducing the flowrate is important to prevent sublimation.

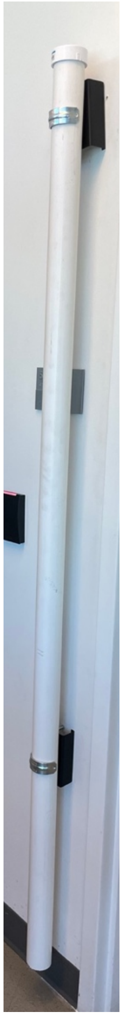

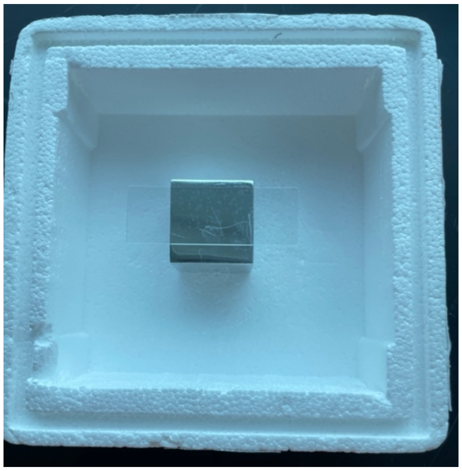

**General method for quantifying ice recrystallization inhibition**

For all polypeptide solutions, Image J (Fiji) was used to analyze the mean grain size (MGS) of the ice crystals. For each cooling splat, three images were taken of different areas of the crystal. Briefly, ice crystals were traced by hand to determine the MGS. Statistical analysis showed that 150μm x 150μm region of the image resulted in a MGS that is reprehensive of the entire population. We randomly selected the region of the image to analyze. For samples that resulted in little to no IRI activity, we circled 75 crystals moving radially outward from a randomly selected point. In all cases, three images were collected for each splat assay, and the ice crystal areas for each image were averaged. The average and standard deviation of the three images are presented for each splat.

**General method for HEK293 thawing and cell culture**

HEK293 cells (used in experiments from passage number 19-30, gifted from Dr. Tara Deans lab) were thawed rapidly at 37˚C. Cells were diluted 10-fold in prewarmed DMEM 1X (Corning) with 10% FBS (VWR) supplemented with 1% penicillin-streptomycin (Gibco) and 1% L-glutamine (Gibco) and centrifuged. The suspension was then centrifuged and the media aspirated. The cell pellet was resuspended in DMEM with 10% FBS supplemented with 1% penicillin-streptomycin and 1% L-glutamine. The cell suspension was transferred to a culture flask and incubated at 37˚C in 5% $CO_2$ to allow the cells to adhere. Cells were grown to 70-80% confluency and then passaged. Cells were passed a minimum of three times post thaw before use in experiments.

**General method for Raji thawing and cell culture**

Raji cells (passage number6, gifted from Dr. Tara Deans lab) were thawed rapidly at 37˚C and diluted in RPMI-1640 medium (ATCC) supplemented with 10% FBS supplemented with 1% penicillin-streptomycin and 1% L-glutamine. The cell suspension was diluted 10-fold in prewarmed RPMI-1640 media with 10% FBS (VWR) supplemented with 1% penicillin-streptomycin (Gibco) and 1% L-glutamine (Gibco). The suspension is then centrifuged and the media aspirated. The cell pellet is resuspended in RPMI-1640 media with 10% FBS supplemented with 1% penicillin-streptomycin and 1% L-glutamine. The suspension was diluted 1:10 in the same media and incubated at 37˚C in 5% $CO_2$. Cells were grown to a concentration of 2 x $10^6$ cells/mL and then diluted 10-fold to allow for expansion. Cells were expanded a minimum of three times post-thaw before use in experiments.

**General method for determining HEK293 cell viability with a CCK8 assay**

HEK 293 cells were plated at a density of 10,000 cells/well in a 96 well plate using DMEM with 10% FBS supplemented with 1% penicillin-streptomycin and 1% L-glutamine. The cells were incubated at 37˚C in 5% $CO_2$ for 24 hours to allow the cells to adhere to the plate. The cells were then treated with polymer dissolved in complete media (DMEM with 10% FBS supplemented with 1% penicillin-streptomycin and 1% L-glutamine) for a final polymer concentration of 1.4, 14, or 70.7 μM. Additionally, other wells of cells are treated with 100-X Triton to kill cells for a positive control or complete media to serve a negative control. The treated cells were again incubated at 37˚C in 5% $CO_2$ for 24 hours. The cells were then dosed with 10uL of CCK-8 solution (Dojindo) and incubated at 37˚C in 5% $CO_2$ for 3 hours. The absorbance at 450 nm of the treated cells was measured after the 3 hours of incubation using a BioTek Synergy HTK multi-mode reader. For the HEK 293 experiments described here and in future methods, cells we used at passage number 21-31.

**General method for human red blood cell (hRBC) preparation**

hRBCs from a single unidentified patient were received one day after the drawing of the patient blood. Prior to acquiring the cells, the hRBCs underwent one centrifugation step and the plasma and buffy coat were removed. Upon, receiving the cells the hRBCs were dissolved in DPBS and then centrifuged at 500 x g for 5 min. The PBS was removed from the pelleted cells. This process was repeated twice to remove platelets and blood proteins. The hRBC pellet was then resuspended in PBS such that the final volume of cells was ~40%. The prepared cells were kept at 4˚C when not in use and all studies were completed within 7 days.

**General method for glycopolypeptide cellular internalization studies with HEK293 cells**

HEK 293 cells were plated on three 24 well plates and incubated in DMEM with 10% FBS supplemented with 1% penicillin-streptomycin and 1% L-glutamine at 37˚C in 5% $CO_2$ overnight to allow the cells to adhere. After 24 hours, the cell media was removed. A 100μM solution of A594-(βGalαGalNAcT$_{0.33}$-*s*-A$_{0.66}$)$_{57}$ in ultrapure water was diluted in DMEM with 10% FBS supplemented with 1% penicillin-streptomycin and 1% L-glutamine to make a 10μM solution of polypeptide. The solution was sterile-filtered before 300μL of the solution was applied to the cells. Additionally, 300μL of DMEM with 10% FBS supplemented with 1% penicillin-streptomycin and 1% L-glutamine was applied to additional wells of cells to serve as the untreated control. The 24 well plates were then placed at 37˚C, RT, and 4˚C to incubate for 1 hour. After one hour, the media was removed and the cells were rinsed 3x with DPBS. Then, 500μL of Hoechst stain diluted in DMEM was then added to each well and cells were incubated for 10 minutes at RT. The cells were then fluorescently imaged to observe the localization of the fluorescent polypeptide.

**General method for glycopolypeptide cellular internalization studies with Raji cells**

Raji cells grown in RPMI-1640 media with 10% FBS supplemented with 1% penicillin-streptomycin and 1% L-glutamine were centrifuged and the media was aspirated. The cell pellet was resuspended in RPMI-1640 media with 10% FBS supplemented with 1% penicillin-streptomycin and 1% L-glutamine with a 10μM of A594-(βGalαGalNAcT$_{0.33}$-*s*-A$_{0.66}$)$_{57}$. To make the polypeptide supplemented media, 100μM solution of A594-(βGalαGalNAcT$_{0.33}$-*s*-A$_{0.66}$)$_{57}$ in ultrapure water was diluted in RPMI-1640 media with 10% FBS supplemented with 1% penicillin-streptomycin and 1% L-glutamine. The media/polypeptide solution was sterile-filtered before being applied to the cells. RPMI-1640 media with 10% FBS supplemented with 1% penicillin-streptomycin and 1% L-glutamine media was applied to additional vials of cells to serve as the untreated control. The polypeptide treated cell suspension was split into three 2mL vials and incubated at 37˚C, 23˚C, or 4˚C for 1 hour. After one hour, the cells were centrifuged and the media was removed. The cells were washed 3x with DPBS and centrifuged between each wash. Then, 500µL of Hoechst stain diluted in RPMI-1640 media with 10% FBS supplemented with 1% penicillin-streptomycin and 1% L-glutamine was added to each vial and the cells were incubated for 10 minutes at 23˚C. The Hoechst media was removed and the cells were resuspended in RPMI-1640 media with 10% FBS supplemented with 1% penicillin-streptomycin and 1% L-glutamine. The cells were then fixed with paraformaldehyde and fluorescently imaged to observe the localization of the fluorescent polypeptide.

**General method for glycopolypeptide cellular internalization studies with hRBCs**

To begin, 100µL of washed hRBCs (prepared as detailed above) were added to a 1.5 mL tube in triplicate. Then 11µL of a 100µM solution of A594-($\beta$Gal$\alpha$GalNAcT$_{0.33}$-*s*-A$_{0.66}$)$_{57}$ in ultrapure water was added to each vial for a final concentration of 10µM. The vials were then placed at 37˚C, RT, and 4˚C to incubate for 1 hour. After one hour, the cells were pelleted by centrifugation at 500 x g for 5 min and the PBS was removed. The cells were rinsed 3x with DPBS and then resuspend. From there 10uL of cell suspension was applied to a slide and then covered with a coverslip. The cells were then fluorescently imaged to observe the localization of the fluorescent polypeptide. No internalization was observed.

**General method for freezing HEK 293 cells for cryoprotection studies**

Cells are cultured until 70-80% confluent. Cells are treated with 2mL of 0.25% trypsin + EDTA for 1 min, neutralized with 4mL of DMEM with 10% FBS supplemented with 1% penicillin-streptomycin and 1% L-glutamine, and then centrifuged at 125 x g for 3 min to pellet the cells. The media and trypsin are aspirated away and the pellet is resuspended in DMEM with 10% FBS and cryoprotectant. The 1mL volumes of suspension were transferred into 2mL cryovials and placed in a CoolCell (Corning) or Mr. Frosty (Nalgene) controlled freezing unit inside a -80˚C freezer for 24hrs. After 24hrs, the vials are thawed and analyzed.

**General method for thawing HEK 293 cells for cryoprotection studies**

To thaw frozen cells used cryopreservation studies, cryovials containing 1mL of cells in cryoprotection media (DMEM with 10% FBS) are transferred from -80˚C to a bed of dry ice. The vials are then rapidly thawed in a 37˚C bath until the ice disappears. The cell suspension is then diluted 10-fold into pre-warmed complete media (DMEM with 10% FBS supplemented with 1%

penicillin-streptomycin and 1% L-glutamine) dropwise. The suspension is then centrifuged and the media is aspirated. The cell pellet is resuspended in complete media and a trypan blue assay is performed to determine cell membrane integrity.

**General method determining membrane integrity with trypan blue assay**

To begin, 10µL of cell suspension is added to 10µL of trypan blue and mixed. Then 10µL of the combined solution is applied to a hemocytometer and the total number of cells and the number of blue cells were counted. The percent of cells with intact membranes was determined using the following equation.

$$\frac{\#\ total\ cells - \#\ blue\ cells}{\#\ total\ cells} * 100 = \%\ cells\ with\ intact\ membranes$$

**General method for freezing and thawing hRBCs for cryoprotection studies**

The cryopreservation of hRBC was conducted following published procedures[9]. In short, 50µL hRBCs (prepared as discussed previously) were mixed with 50µL of DPBS containing cryoprotectants or control solution in cryovials. The vials were then placed in a liquid nitrogen bath for 20 mins. The samples were then thawed at room temperature for 20 mins. Control samples were also prepared according to this publication. hRBCs were added to water and frozen for 100% hemolysis and for 0% hemolysis, hRBCs were incubated with DPBS at room temperature for 1 hr. For all hRBC cryopreservation experiments, 2-hydroxyethyl starch (Spectrum Chemical, H3012) was used.

**General methods for measuring hRBC hemolysis and cell recovery**

Hemolysis and cell recovery were determined with modifications to published procedures [9]. After thawing 60µL of cell suspension was diluted into 540µL of DPBS and centrifuged at 500 x g for 5 min. Then 20 µL of the supernatant was added to 480 µL of DPBS. Then 100µL of the diluted supernatant solution was added to a 96-well plate in triplicate. The absorbance at 414 nm was measured. Hemolysis and cell recovery were determined following the equations in the published procedure.[9]

#### IV. Supplementary Figures and Tables

**Table S1.** Concentrations of glycopolypeptide solutions used for CD samples. All CD data in the manuscript (Fig. 2) and supplementary information are reported in mean residue molar ellipticity, which normalizes for samples of different concentrations and polypeptide lengths.

| Name[a] | mg/mL | mM |
|---|---|---|
| $(\beta GalT_{0.33}\text{-}s\text{-}A_{0.66})_{93}$ | | 1.14 |
| $(\alpha GalT_{0.33}\text{-}s\text{-}A_{0.66})_{93}$ | 0.5 | |
| $(\beta GalNAcT_{0.33}\text{-}s\text{-}A_{0.66})_{93}$ | 0.5 | |
| $(\alpha GalNAcT_{0.33}\text{-}s\text{-}A_{0.66})_{93}$ | 0.5 | |
| $(\beta Gal\alpha GalNAcT_{0.33}\text{-}s\text{-}A_{0.66})_{28}$ | 0.25 | |
| $(\beta Gal\alpha GalNAcT_{0.33}\text{-}s\text{-}A_{0.66})_{57}$ | 0.5 | |
| $(\beta Gal\alpha GalNAcT_{0.33}\text{-}s\text{-}A_{0.66})_{138}$ | 0.5 | |
| $(\beta Gal\alpha GalNAcT_{0.33}\text{-}s\text{-}A_{0.66})_{170}$ | 0.5 | |
| $(\beta Gal\alpha GalNAcT_{0.5}\text{-}s\text{-}A_{0.5})_{52}$ | | 1.85 |
| $(\beta Gal\alpha GalNAcT_{0.66}\text{-}s\text{-}A_{0.33})_{46}$ | | 1.14 |
| $(\beta GalNAcT_{0.4}\text{-}s\text{-}A_{0.6})_{93}$ | | 1.69 |
| $(\beta GalT_{0.5}\text{-}s\text{-}A_{0.5})$ | | 1.48 |
| $(\beta GalT_{0.66}\text{-}s\text{-}A_{0.33})$ | | 0.85 |
| $(\beta GalT_{0.75}\text{-}s\text{-}A_{0.25})$ | | 0.62 |
| $(\beta GalT)_n$ | | 1.07 |
| $(\beta Gal\alpha GalNAcT_{0.33}\text{-}s\text{-}A_{0.66})_{57}$ in PBS | | 1.23 |
| $(BGal\alpha GalNAcT_{0.33}\text{-}s\text{-}A_{0.66})_{98}$ from temp. melt | | 2.47 |

**Table S1.** Statistical significance of quantified IRI MGS from manuscript Fig. 4B, mean and standard deviation reported in Fig. 4B. Data processed with one-way ANOVA and post-hoc Tukey test. ** indicates p<0.01, * indicates p <0.045, and ns is not significant.

| | 28mer | 57mer | 170mer | αGal | βGal | αGalNAc | βGalNAc | PVA 50 μM | PVA 100μM | 5% DMSO |
|---|---|---|---|---|---|---|---|---|---|---|
| 28mer | - | ** | ** | ns | ** | ns | ** | ** | ** | ** |
| 57mer | | - | ** | ** | ** | ** | ns | ** | ns | ** |
| 170mer | | | - | ** | ** | ** | ** | ** | * | ** |
| αGal | | | | - | * | ** | ** | ** | ** | ** |
| βGal | | | | | - | ** | ** | ns | ** | ** |
| αGalNAc | | | | | | - | ** | ** | ** | ** |
| βGalNAc | | | | | | | - | ** | ns | ** |
| PVA 50 μM | | | | | | | | - | ** | ** |
| PVA 100 μM | | | | | | | | | - | ** |
| 5% DMSO | | | | | | | | | | - |

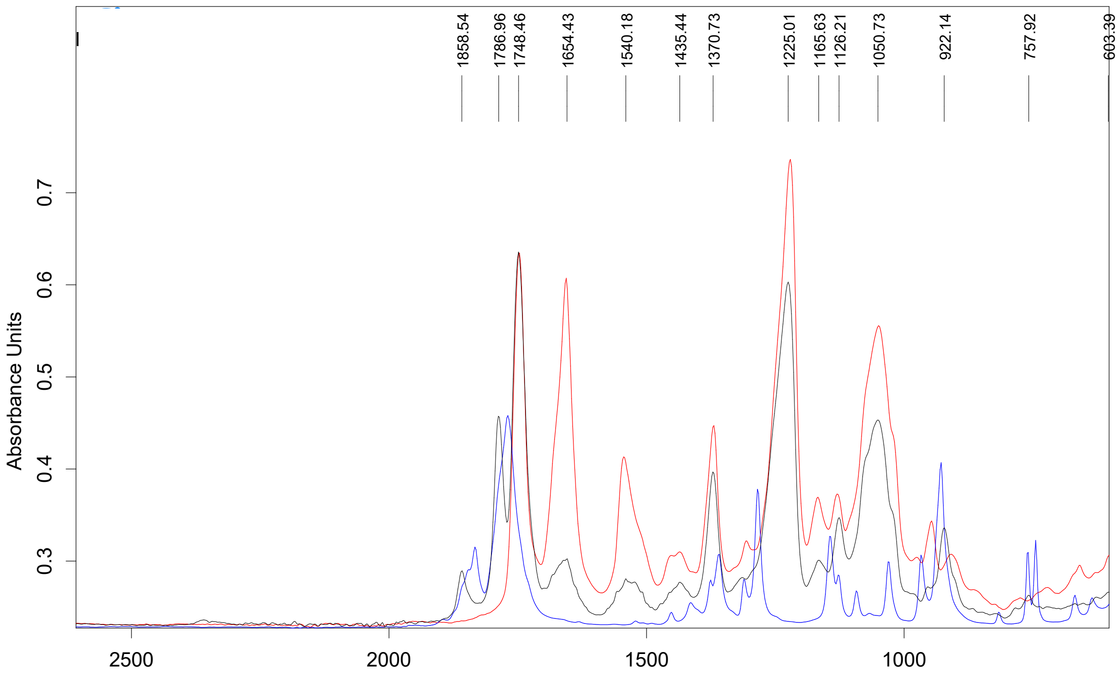

**Figure S1:** ATR-FTIR showing the disappearance of Ala NCA and βGalαGalNAcT NCA and formation of (βGalαGalNAcT$_{0.33}$-*s*-PA$_{0.66}$)$_n$.

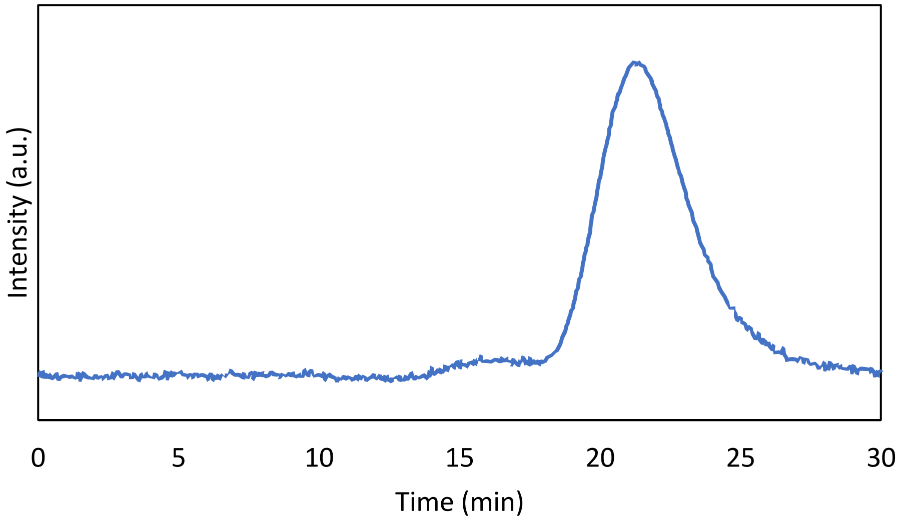

**Figure S2:** GPC/MALS trace for (βGalαGalNAcT$_{0.33\text{-}s\text{-}}$PA$_{0.66}$)$_{78}$

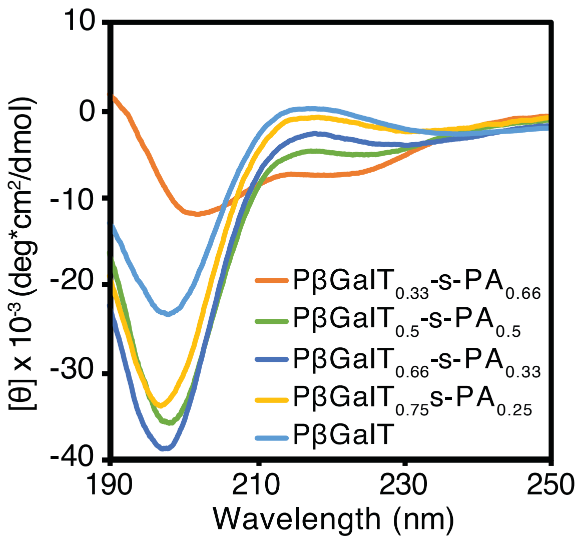

**Figure S3:** CD spectra of ($\beta$GalT$_x$-*s*-A$_y$)$_n$ at various amino acid concentrations. Spectra in ultrapure water at 25˚C.

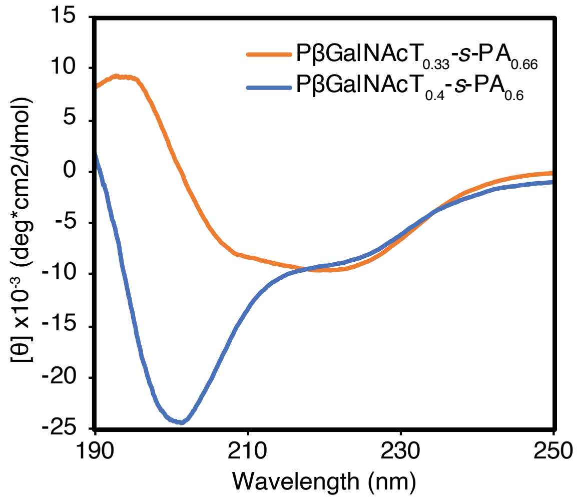

**Figure S4:** CD spectra of ($\beta$GalNAcT$_x$-*s*-A$_y$)$_n$ at various amino acid concentrations. An increase in 6.6% GalNAcThr content increases the solubility of the polymer. Spectra in ultrapure water at 25˚C.

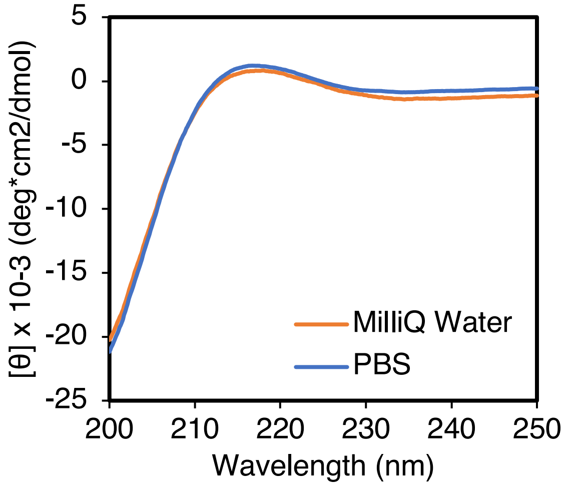

**Figure S5:** CD spectra of (βGalαGalNAcT$_{0.33}$-*s*-A$_{0.66}$)$_{57}$ in PBS versus ultrapure water water at 25˚C.

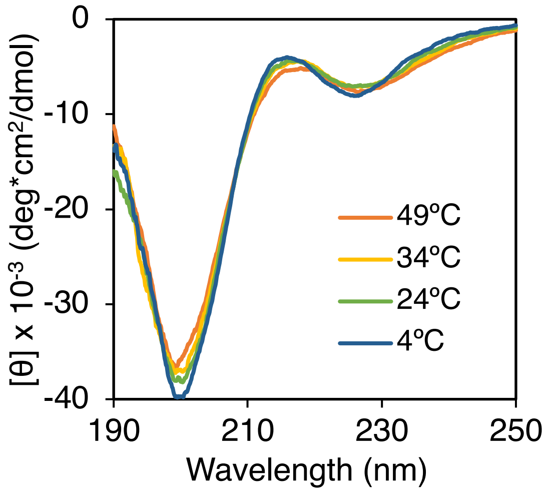

**Figure S6:** CD spectra of (βGalαGalNAcT$_{0.33}$-*s*-A$_{0.66}$)$_{98}$ at varied temperatures in ultrapure water.

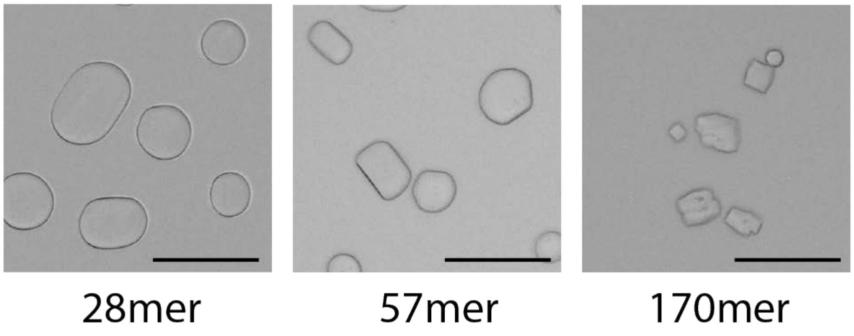

**Figure S7:** Ice shaping of (βGalαGalNAcT$_{0.33}$-*s*-A$_{0.66}$)$_n$ DP = 28, 57, 170 at 0.5 mg/mL. Scale bar = 100μm

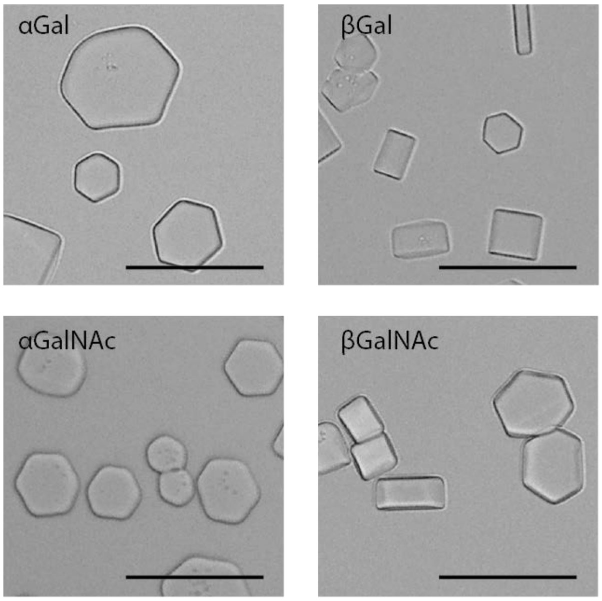

**Figure S8:** Ice shaping of (glyco$T_{0.33}$-*s*-$A_{0.66}$)$_{93}$ at 0.5 mg/mL. Scale bar = 100μm

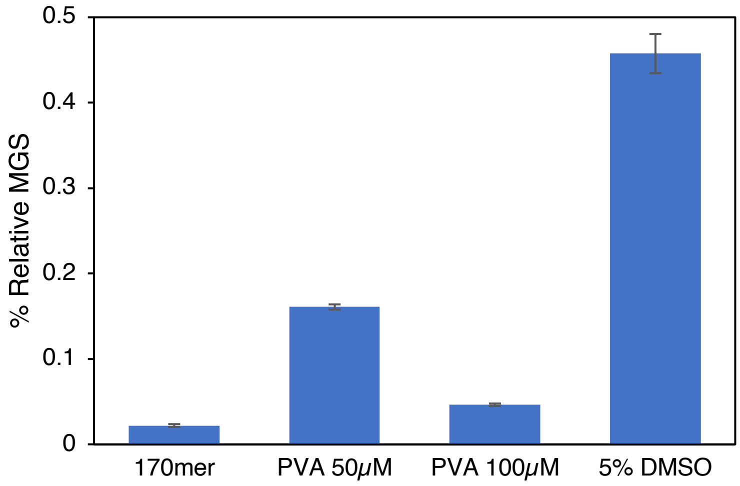

**Figure S9:** Quantified IRI data as % MGS relative to PBS for (βGalαGalNAc$T_{0.33}$-s-$A_{0.66}$)$_{170}$, PVA, and DMSO. sAFGP concentration is 71 μM in PBS. PVA concentrations are indicated on plot. PVA and DMSO in PBS. IRI experiments were conducted alongside the emerging cryoprotectant PVA ($MW_{avg}$ 13–23kDa, similar to the MW range of native AFGPs). 5% DMSO was selected because it is commonly used as a cryoprotective agent at this concentration. Ice crystal MGS was determined from cooling splat assays. Mean and standard deviation are plotted. Statistical significance presented in Table S1.

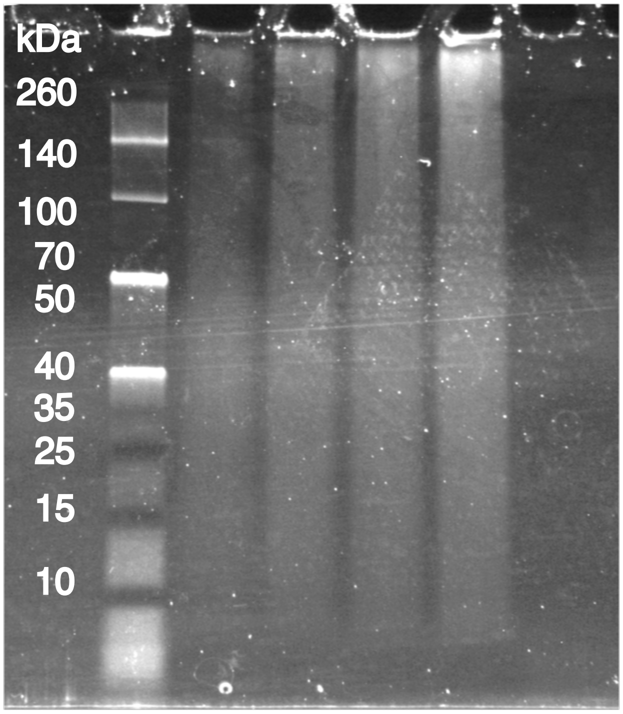

**Figure S10:** SDS-Page used to determine the concentration of sAFGP needed for protease digestion studies. ($\beta$Gal$\alpha$GalNAcT$_{0.33}$-*s*-A$_{0.66}$)$_{57}$ used for this experiment.

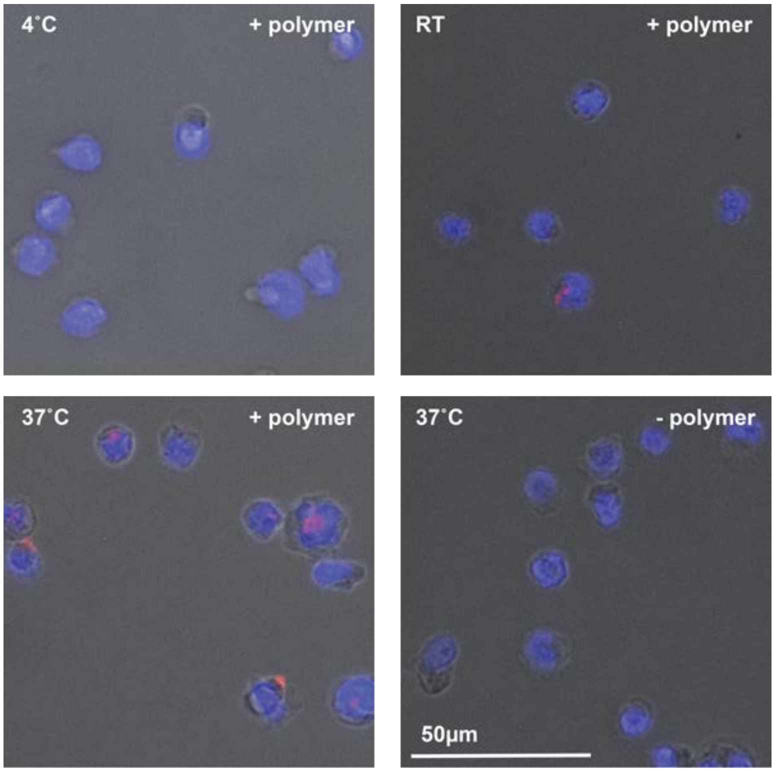

**Figure S11:** Internalization AF594-($\beta$Gal$\alpha$GalNAcT$_{0.33}$-*s*-A$_{0.66}$)$_{57}$ in Raji cells.

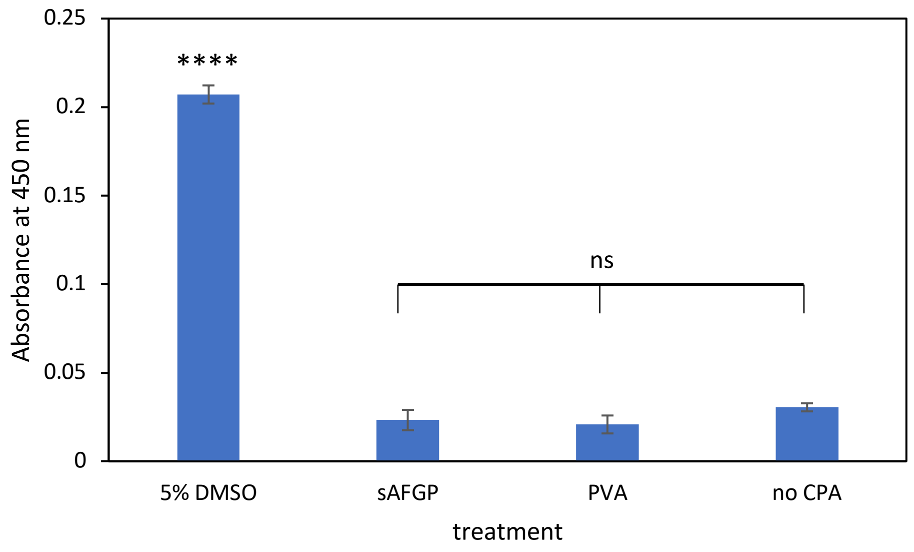

**Figure S12:** Cryopreservation of HEK293 cells with varying treatments. Viability related to absobance as determined by CCK8 assay. Cells treated with 20mg/mL sAFGP ($\beta$Gal$\alpha$GalNAcT$_{0.33}$-*s*-A$_{0.66}$)$_{57}$ and PVA(MW$_{avg}$ 13-23 kDa. Statistical significance was determined with a one-way ANOVA and post-hoc Tukey tests. ns indicates nonsignificant and **** indicates p<0.00001

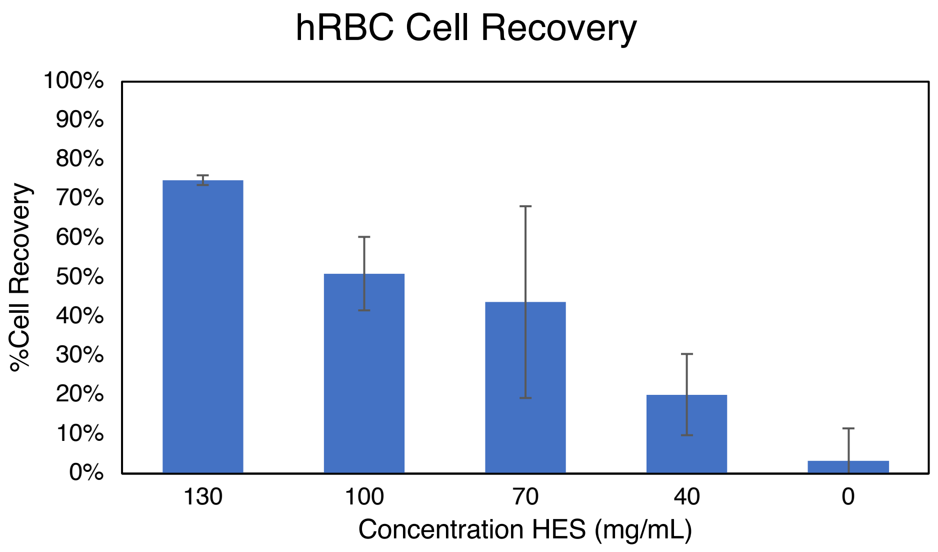

**Figure S13:** Cryopreservation of hRBC with varying HES concentrations.

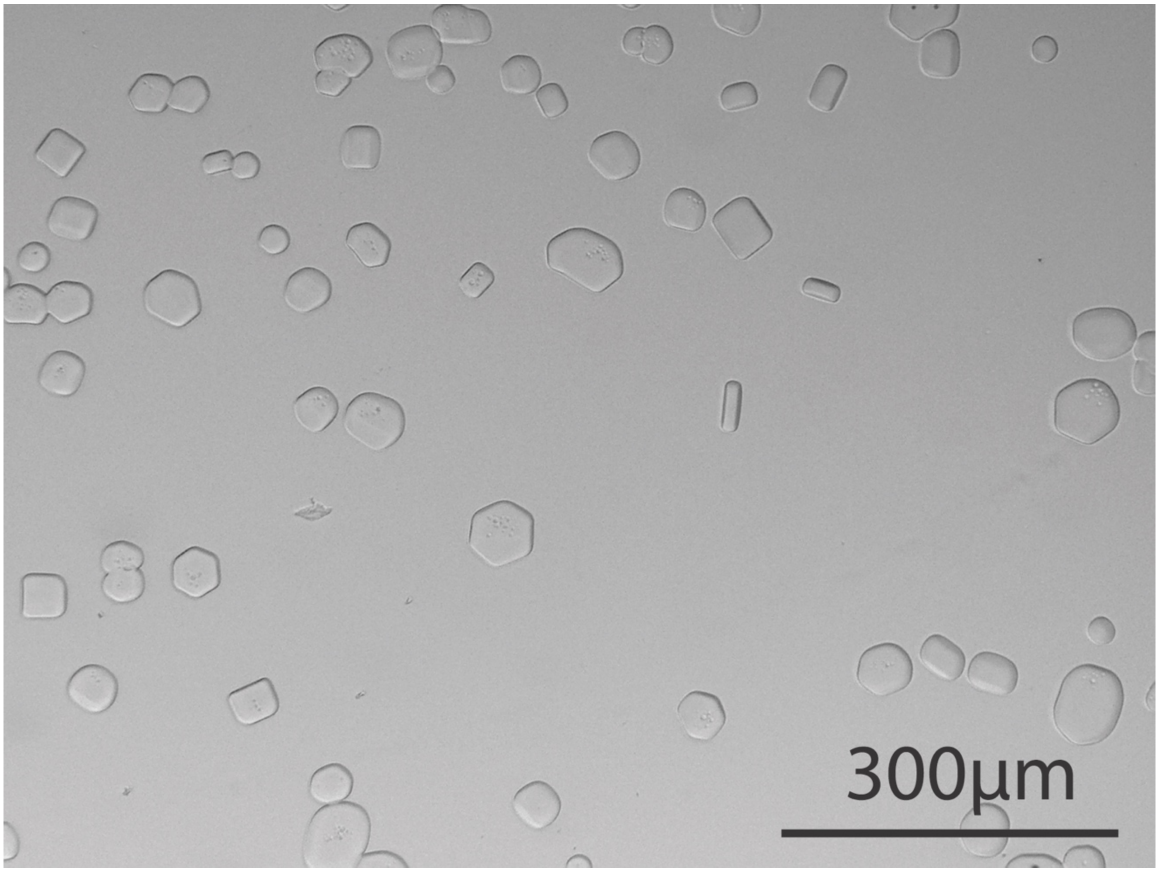

**Figure S14:** Field of view image of ice shaping with (βGalαGalNAcT0.33-s-PAla0.66)57

#### V. Cooling Splat – Ice Recrystallization Inhibition

Note: Statistical significance determined with a one-way ANOVA and post-hoc Tukey tests, ns indicates not significant, * indicates $p<0.05$ and ** $p<0.01$. Scale bar for all IRI images is 300µm.

$(\beta Gal\alpha GalNAcT_{0.33}\text{-}s\text{-}A_{0.66})_{28}$

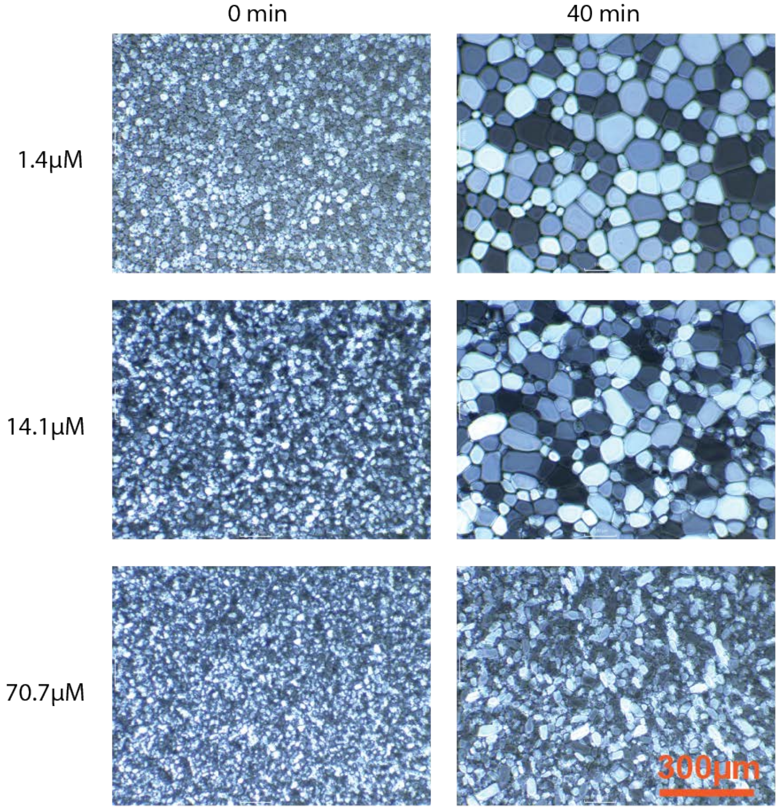

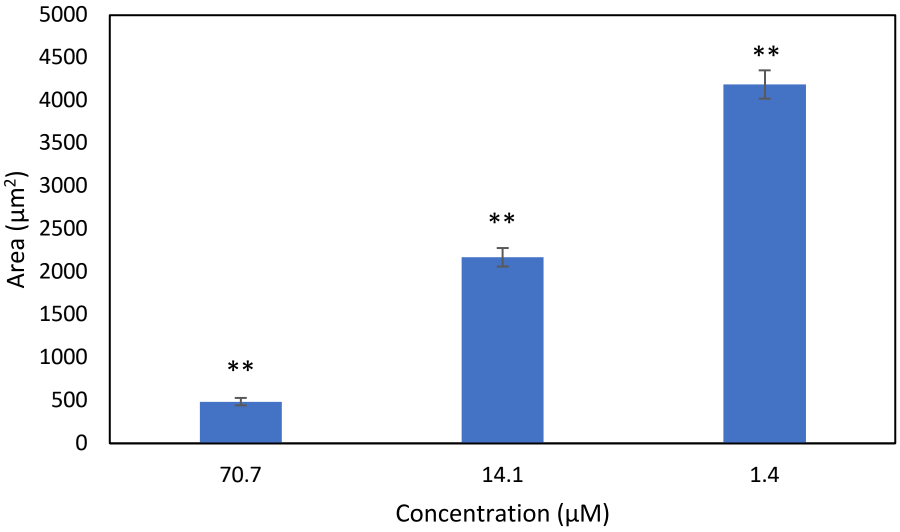

$(\beta Gal\alpha GalNAcT_{0.33}\text{-}s\text{-}A_{0.66})_{57}$

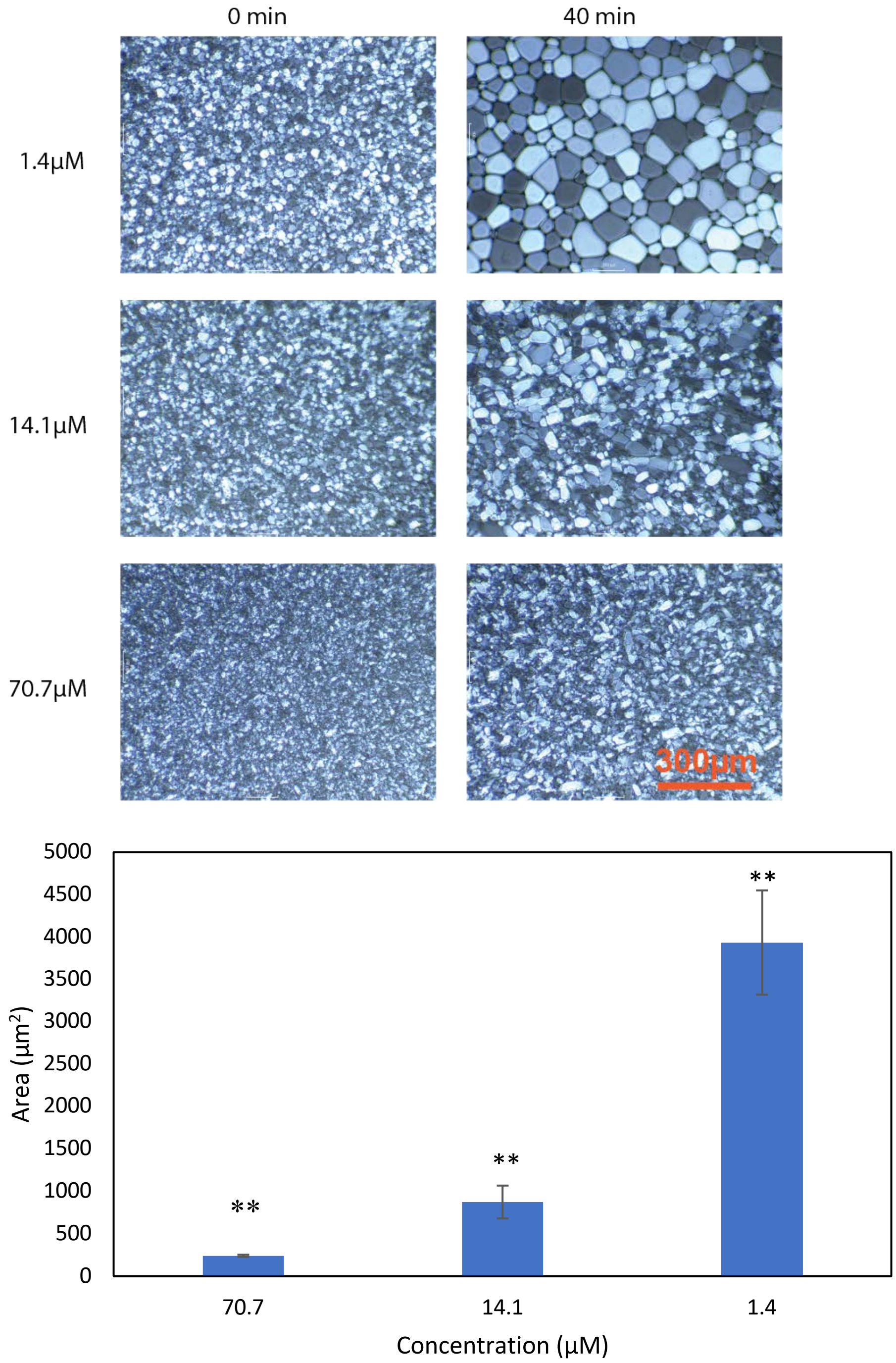

(βGalαGalNAcT$_{0.33}$-*s*-A$_{0.66}$)$_{170}$

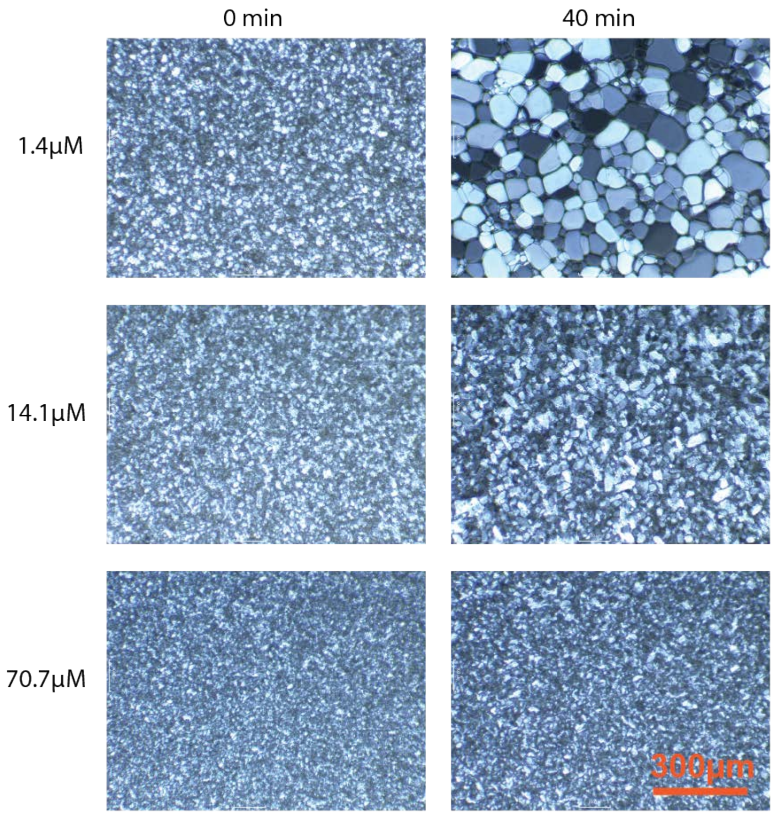

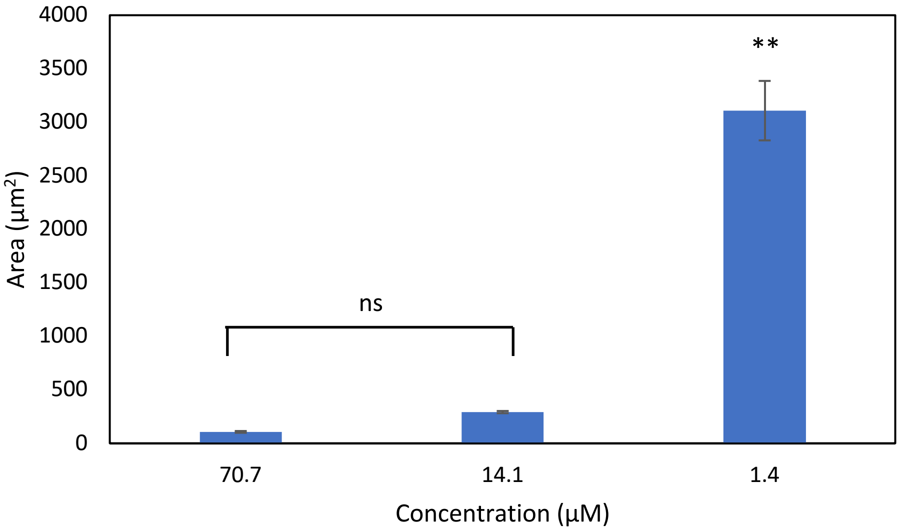

($\beta$Gal$\alpha$GalNAcT$_{0.5}$-*s*-A$_{0.5}$)$_{52}$

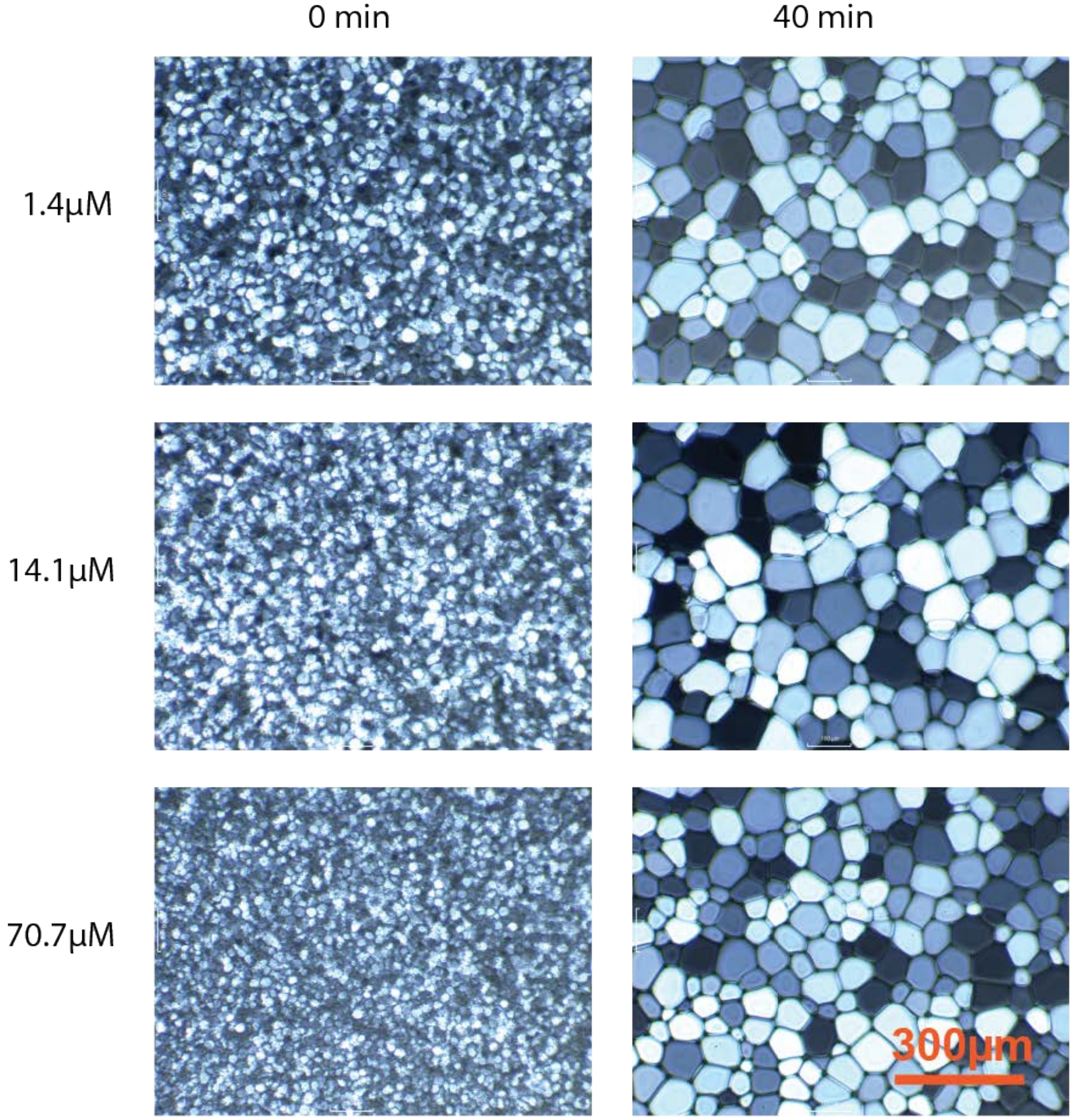

($\beta$Gal$\alpha$GalNAcT$_{0.66}$-*s*-A$_{0.33}$)$_{46}$

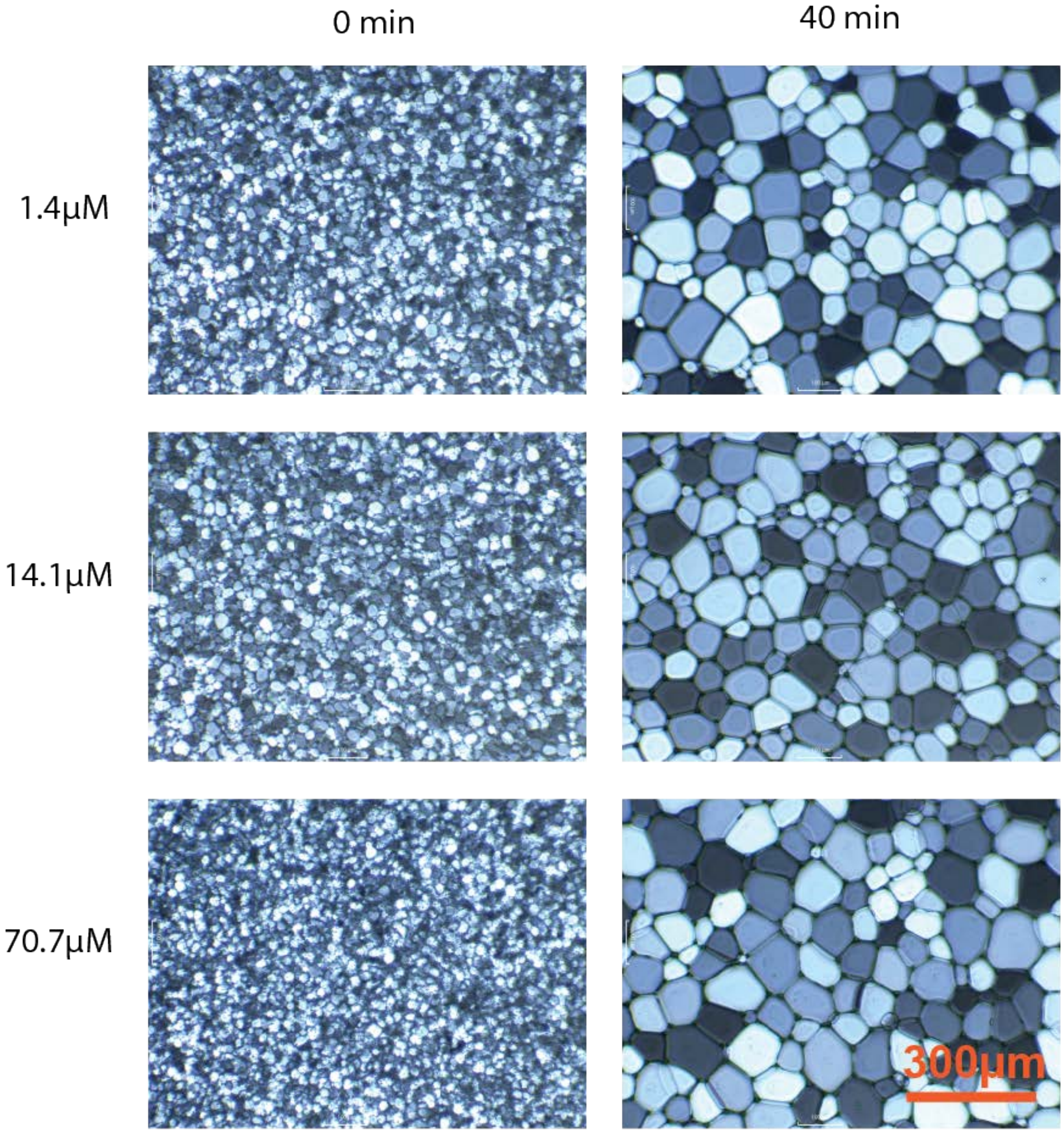

$(\alpha GalNAcT_{0.33}\text{-}\mathit{s}\text{-}A_{0.66})_{93}$

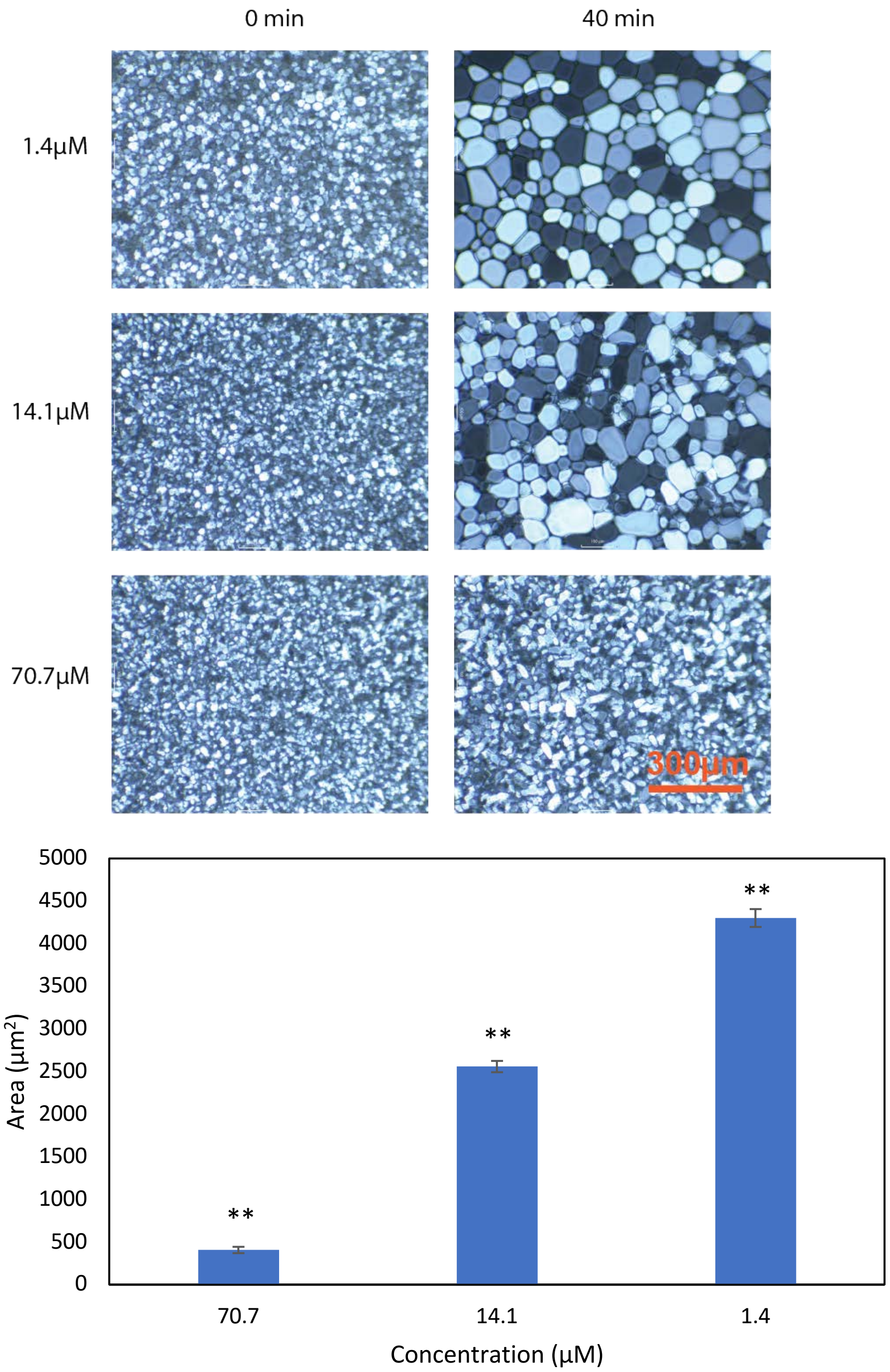

($\alpha$GalT$_{0.33}$-*s*-A$_{0.66}$)$_{93}$

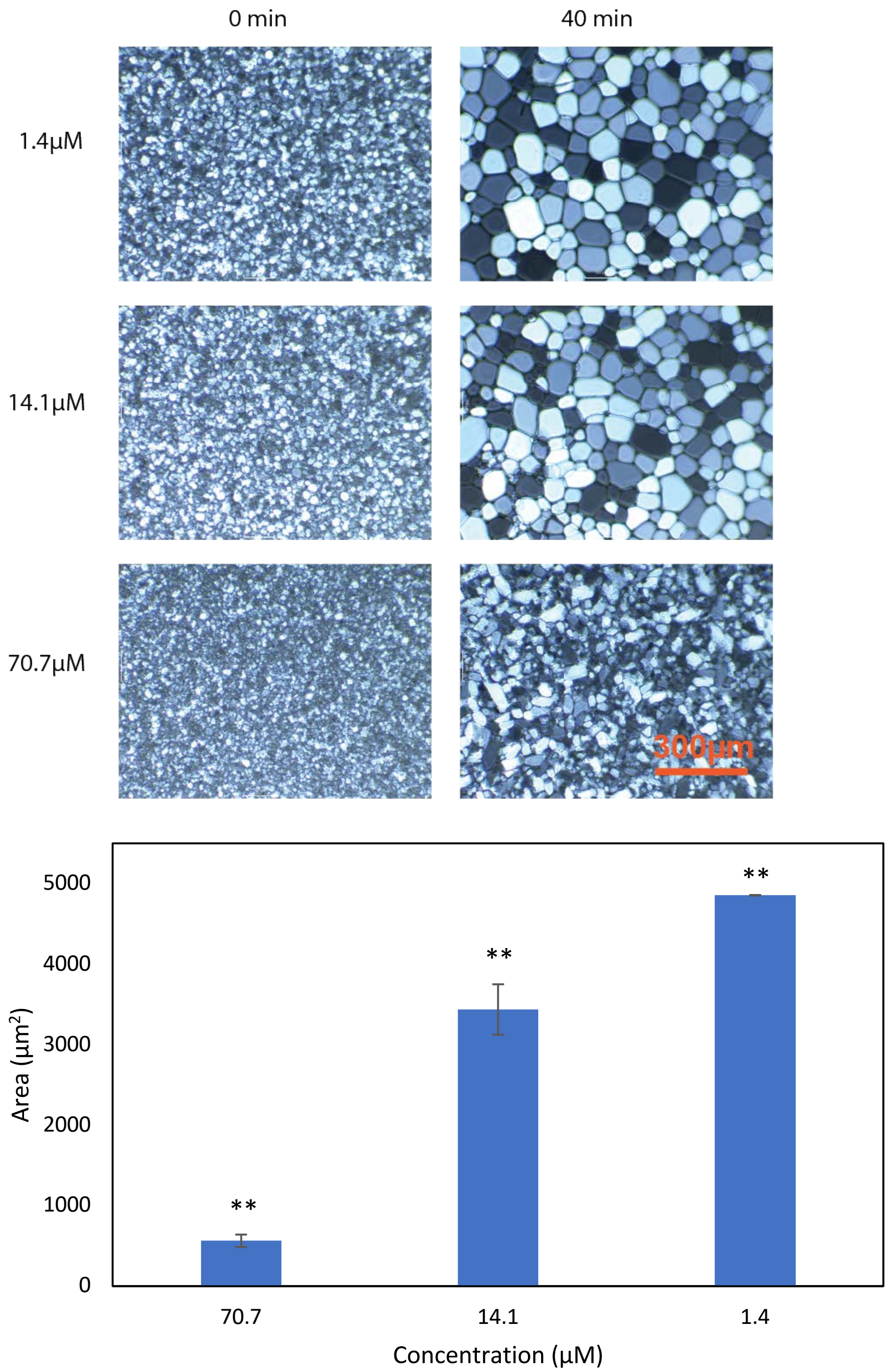

$(\beta GalNAcT_{0.33}\text{-}s\text{-}A_{0.66})_{93}$

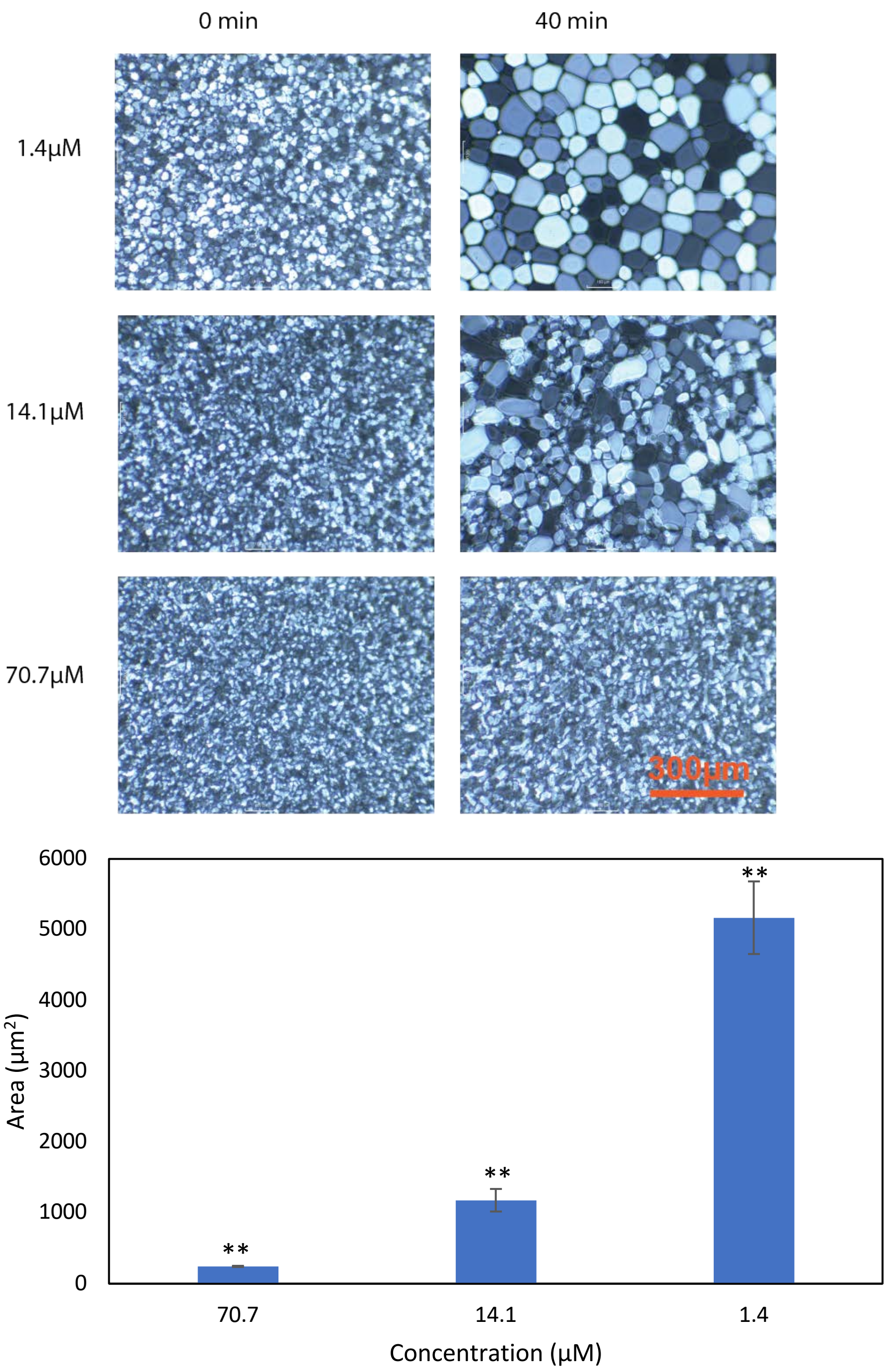

($\beta$GalT$_{0.33}$-*s*-A$_{0.66}$)$_{93}$

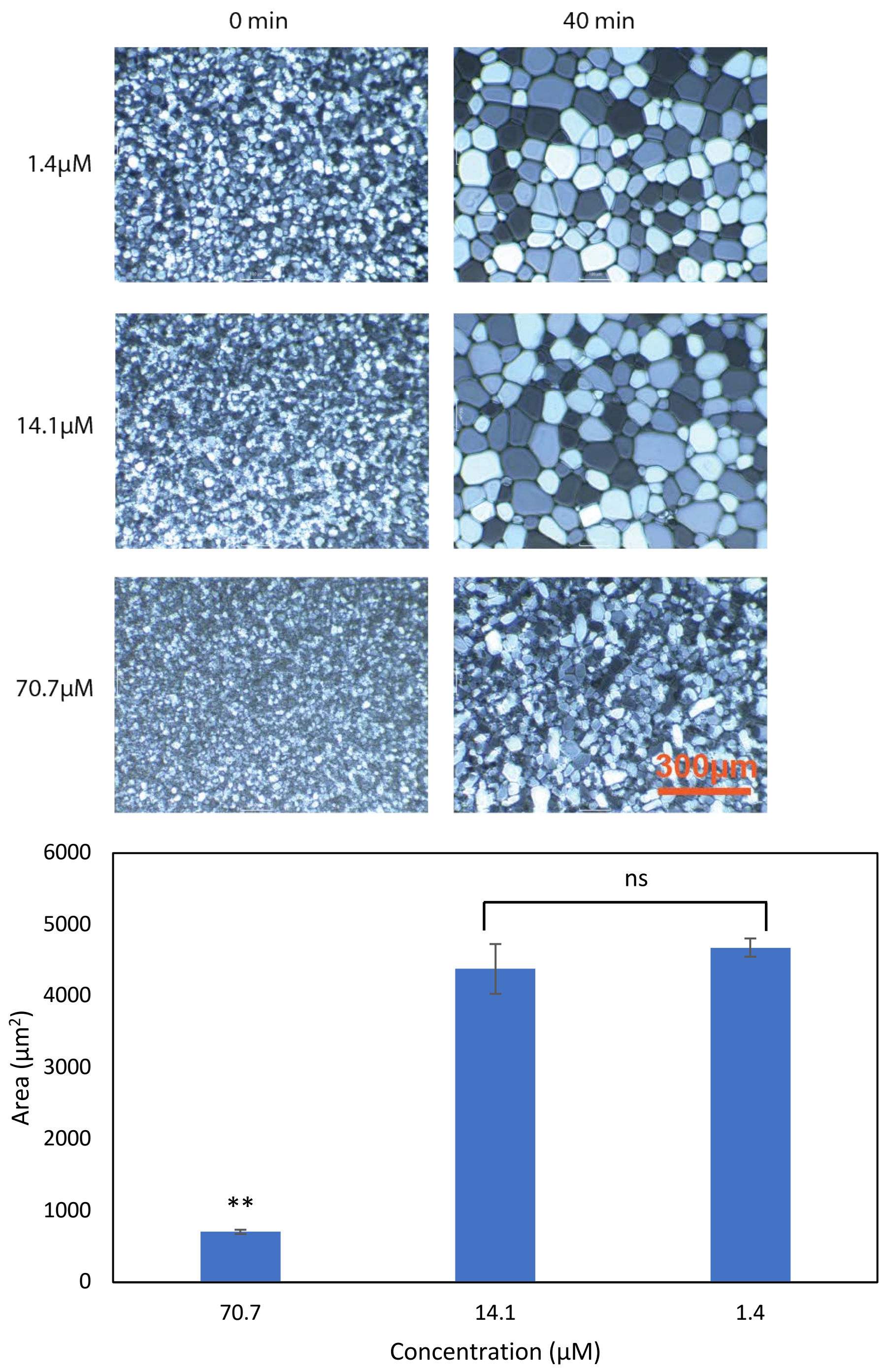

$(\beta Gal\alpha GalNAcT_{0.33}\text{-}s\text{-}A_{0.66})_n$ at 70.7μM

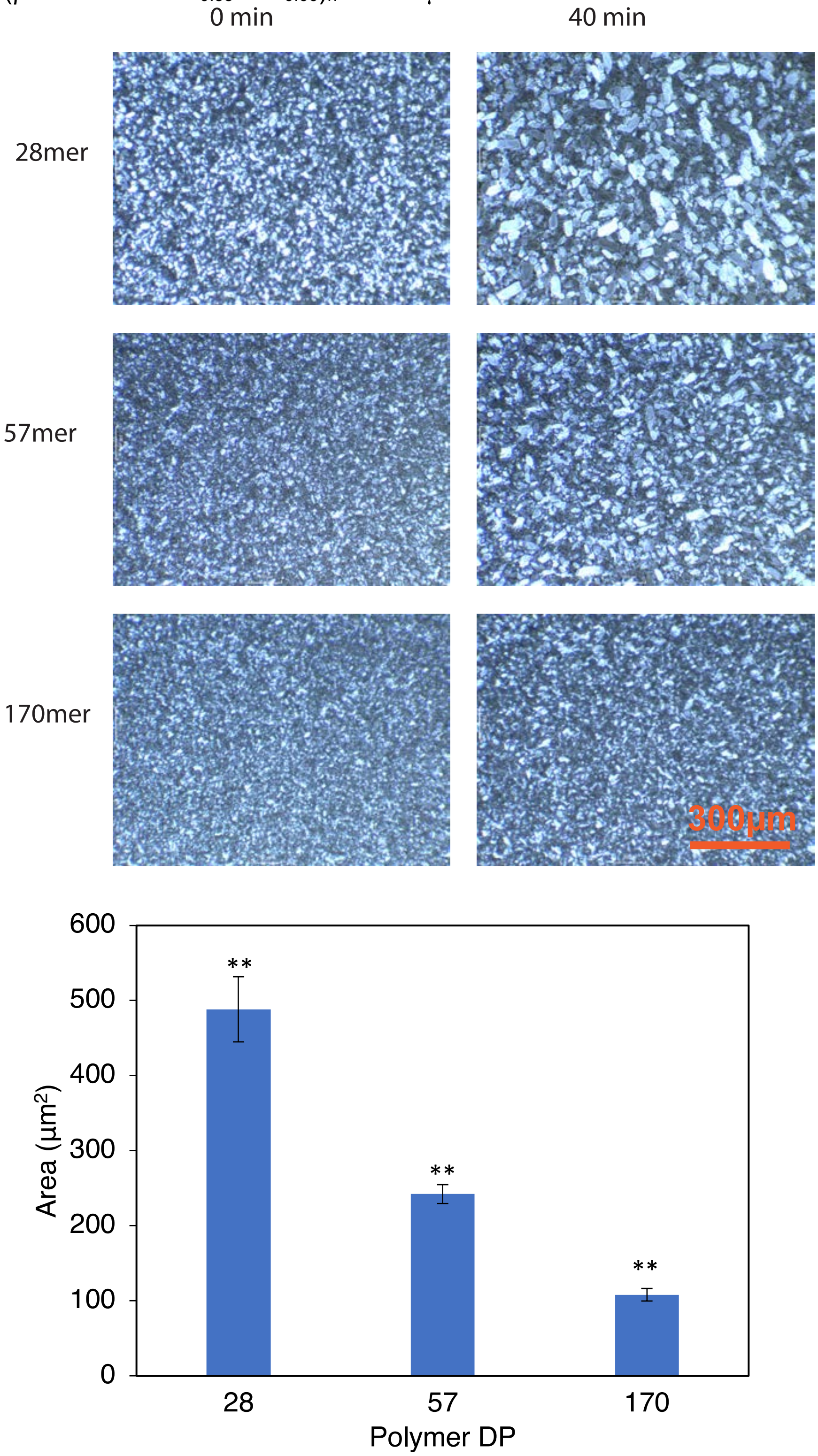

($\beta$Gal$\alpha$GalNAcT$_x$-*s*-A$_y$) at 70.7$\mu$M

x:y 0 min 40 min

1:2

1:1

2:1

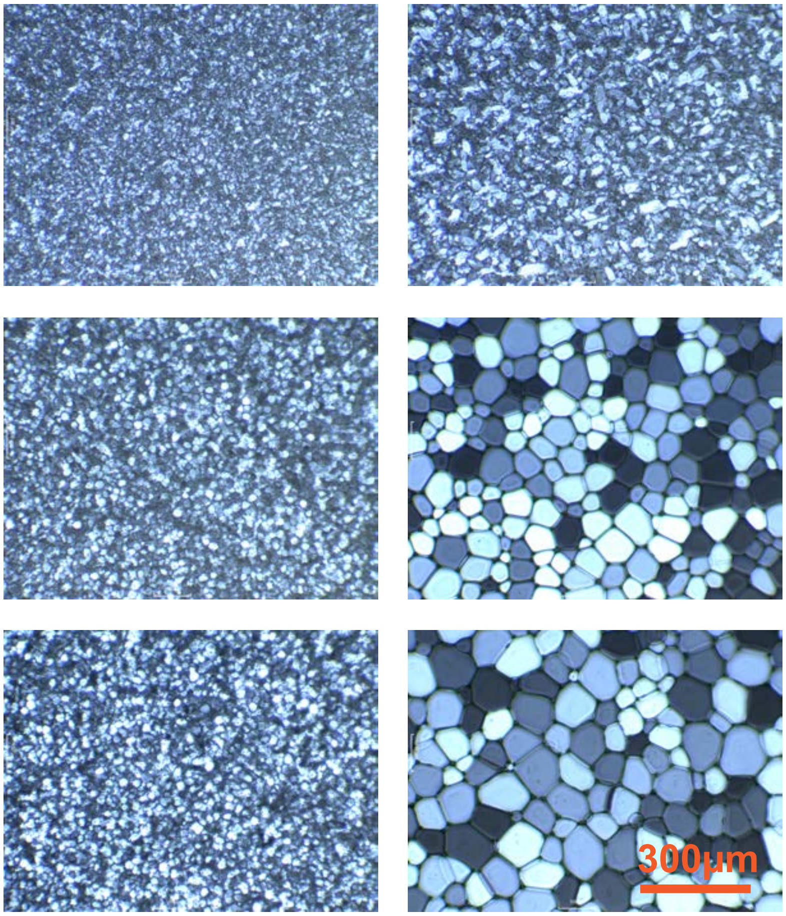

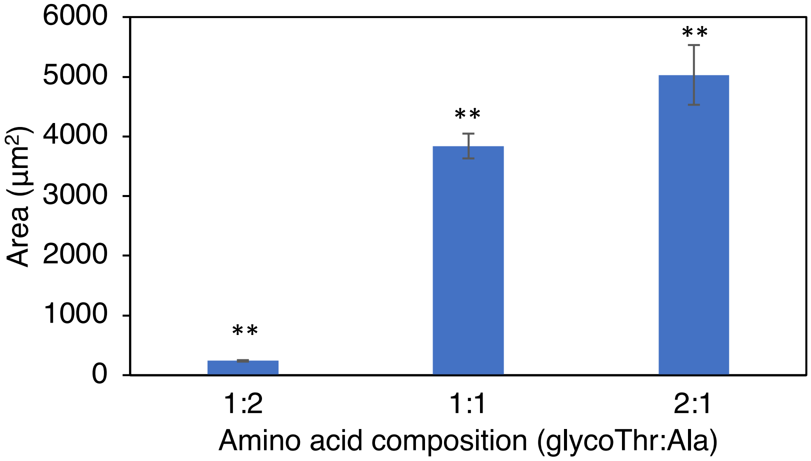

$(glycoT_{0.33}\text{-}s\text{-}A_{0.66})_{93}$ at 70.7μM

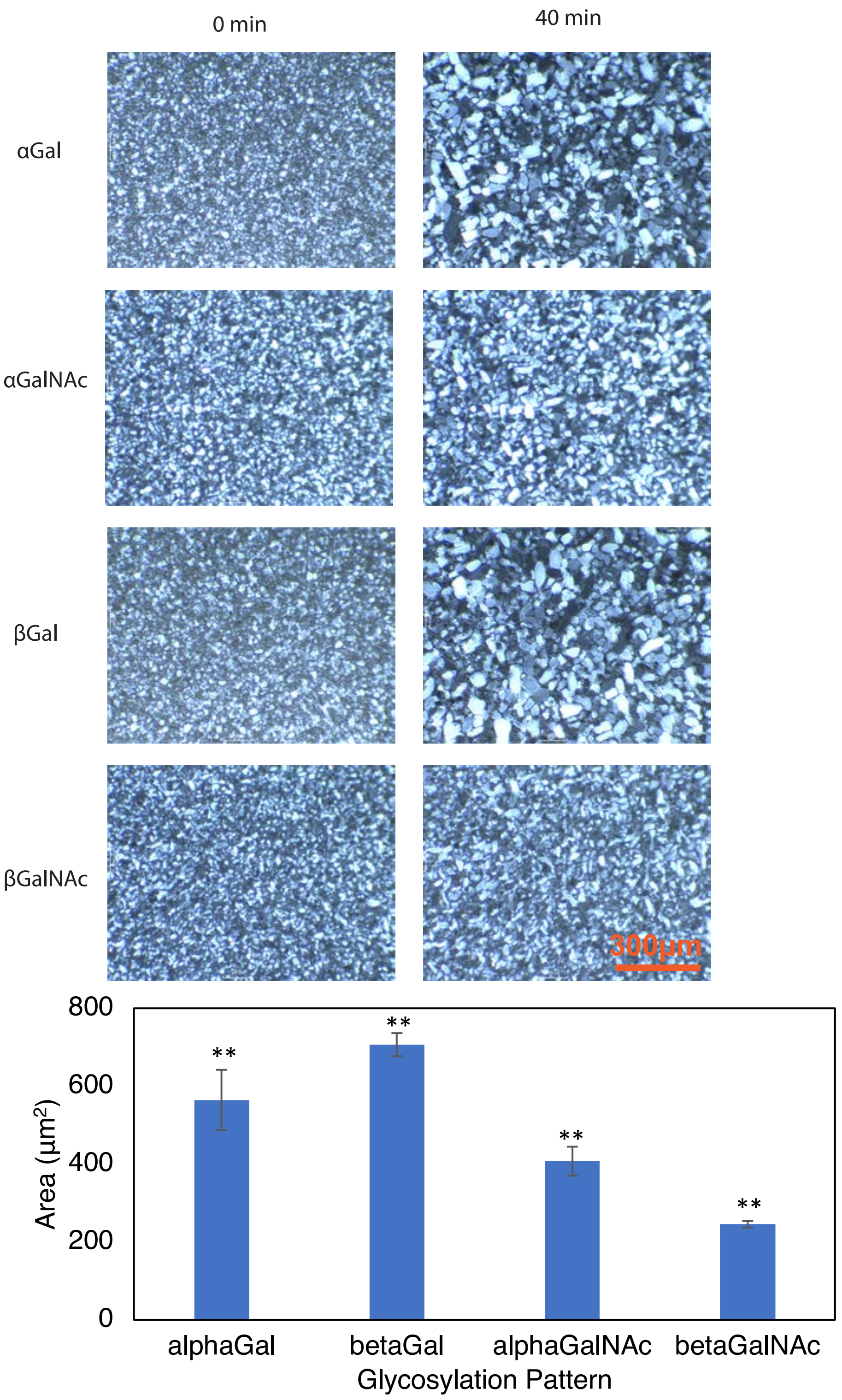

$(\beta Gal\alpha GalNAcT_{0.33}\text{-}s\text{-}A_{0.66})_n$ at 0.5 mg/mL

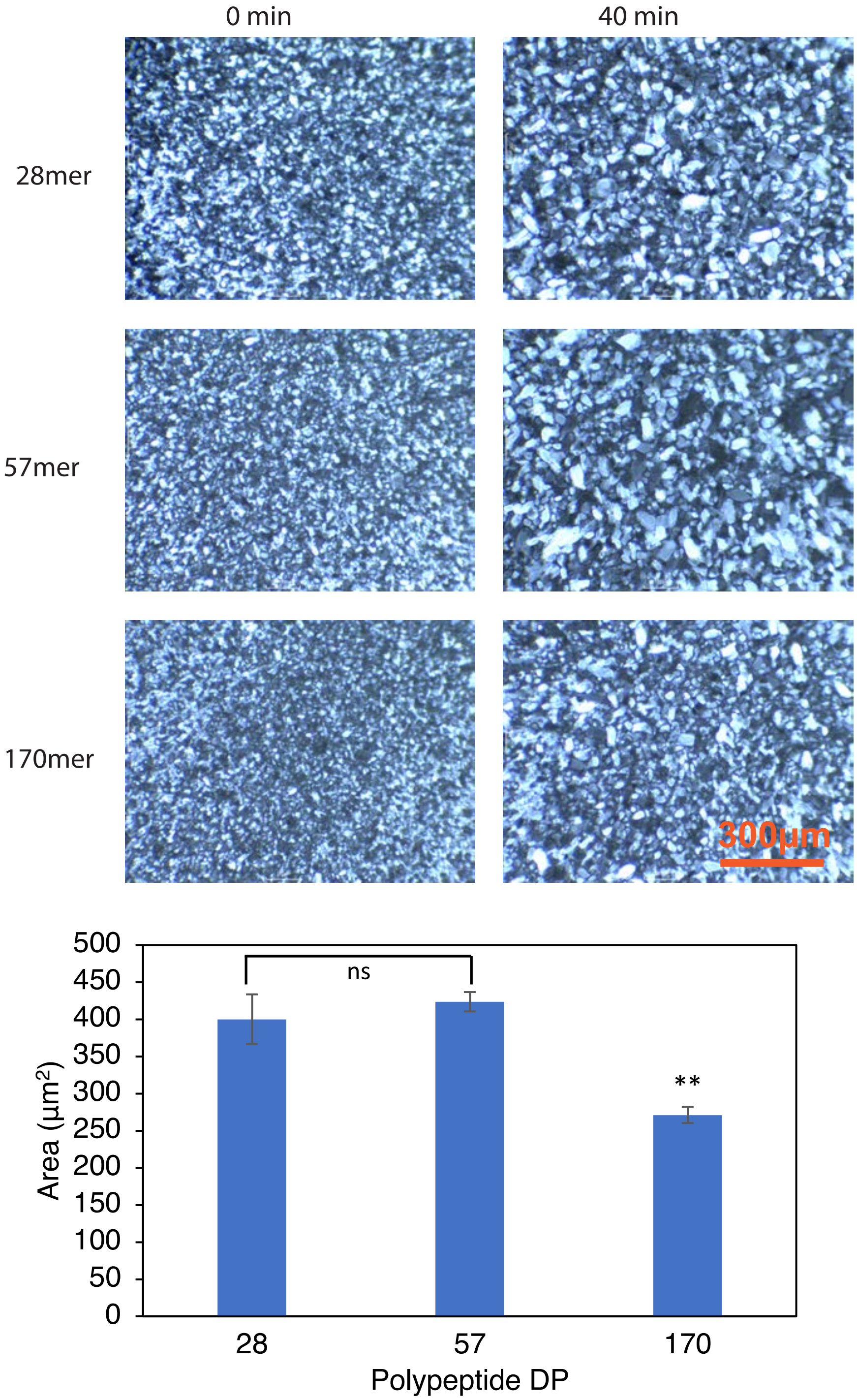

(glycoT$_{0.33}$-*s*-A$_{0.66}$)$_{93}$ at 0.5. mg/mL

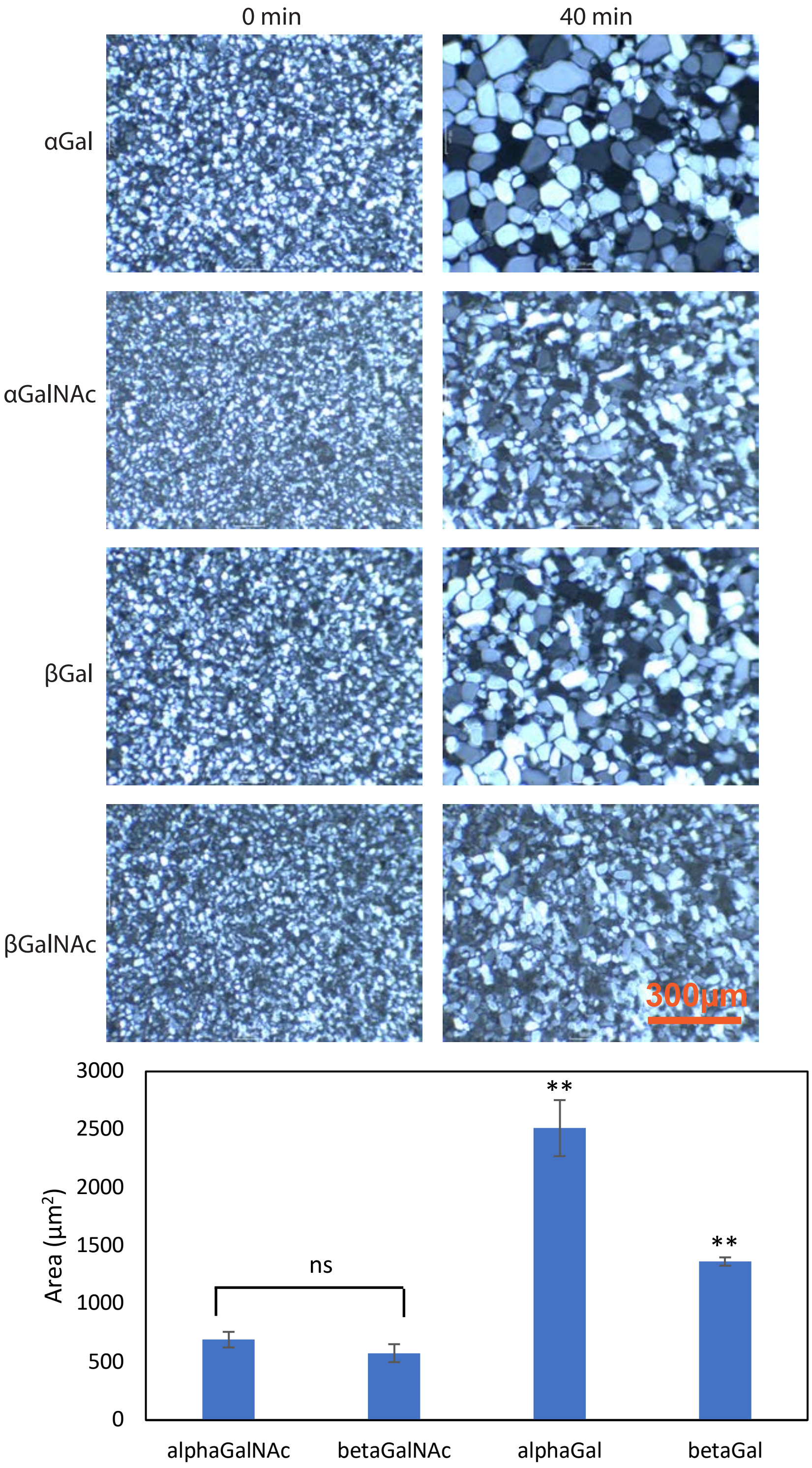

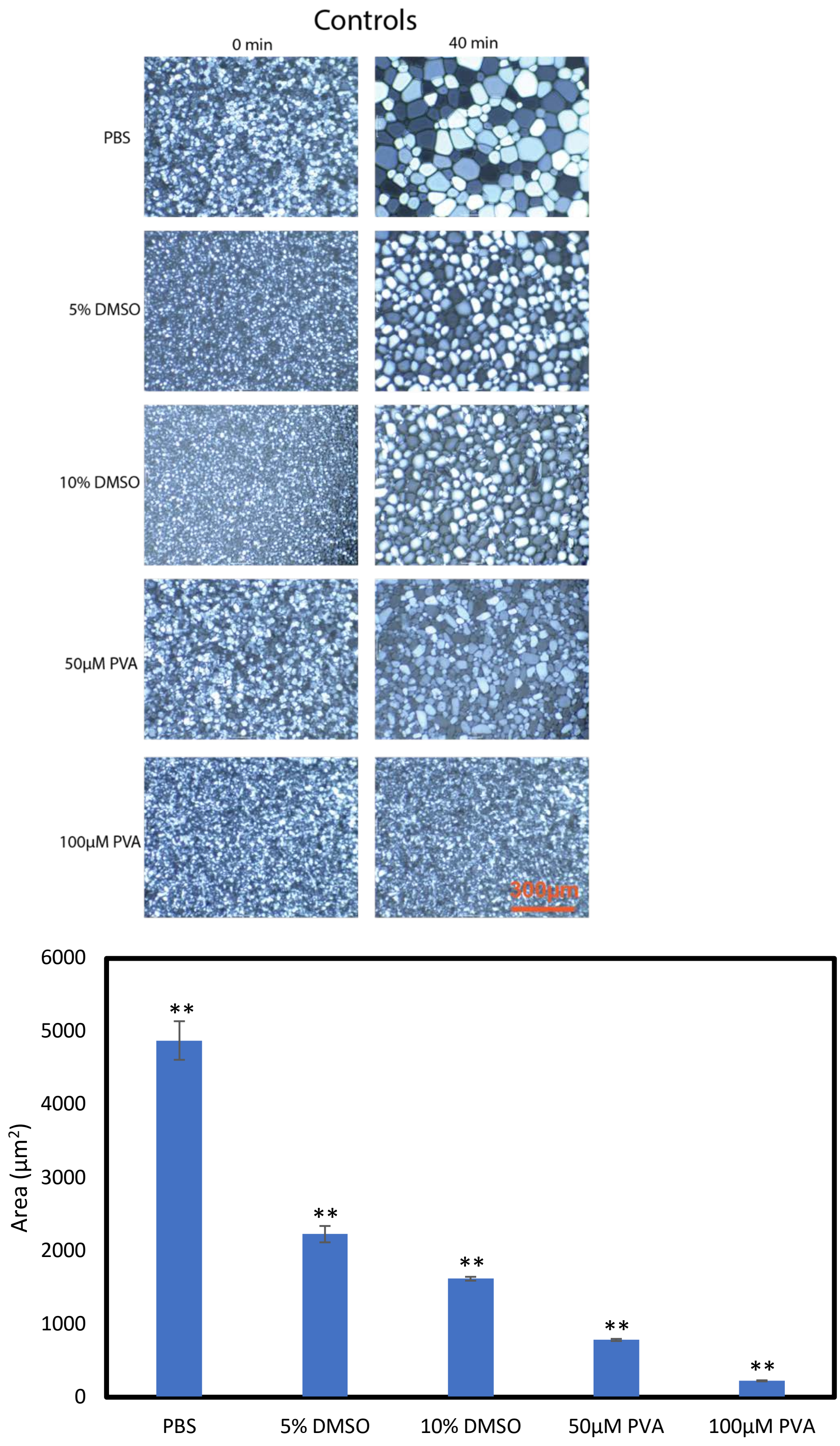
Controls
0 min
40 min
PBS
5% DMSO
10% DMSO
50µM PVA
100µM PVA
300µm
6000
5000
4000
3000
2000
1000
0
Area (µm2)
**
PBS
5% DMSO
10% DMSO
50µM PVA
100µM PVA

### VI. NMR Spectra

#### Ala NCA

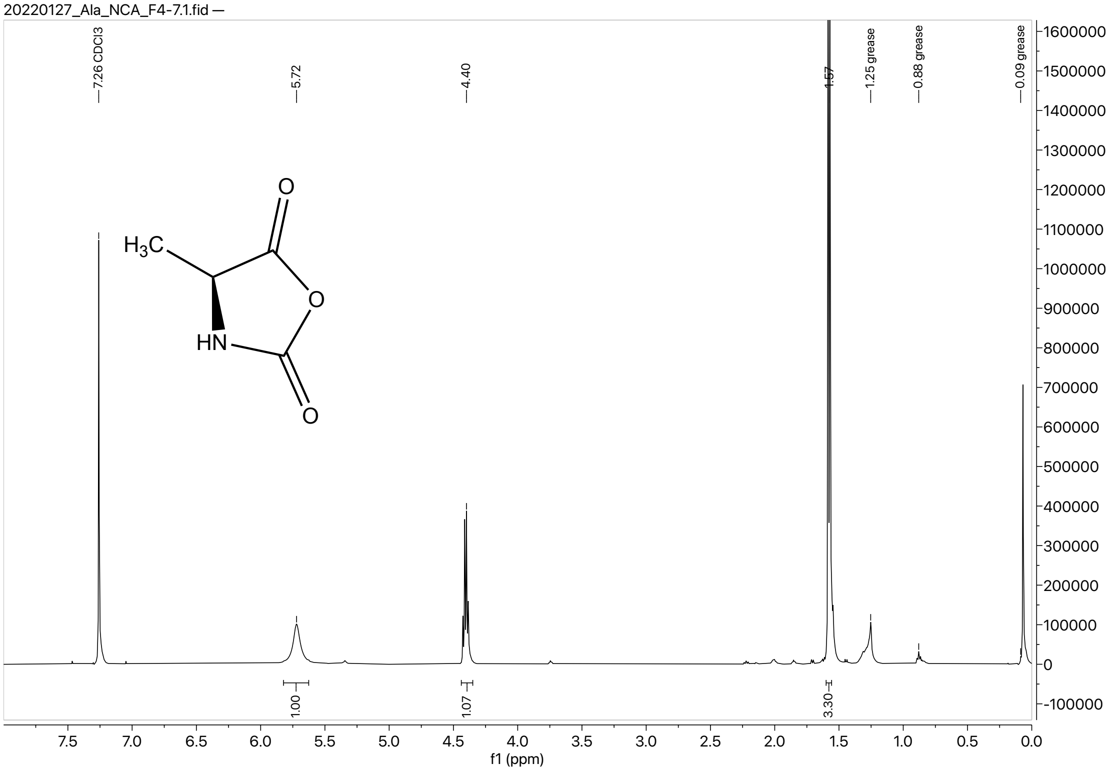

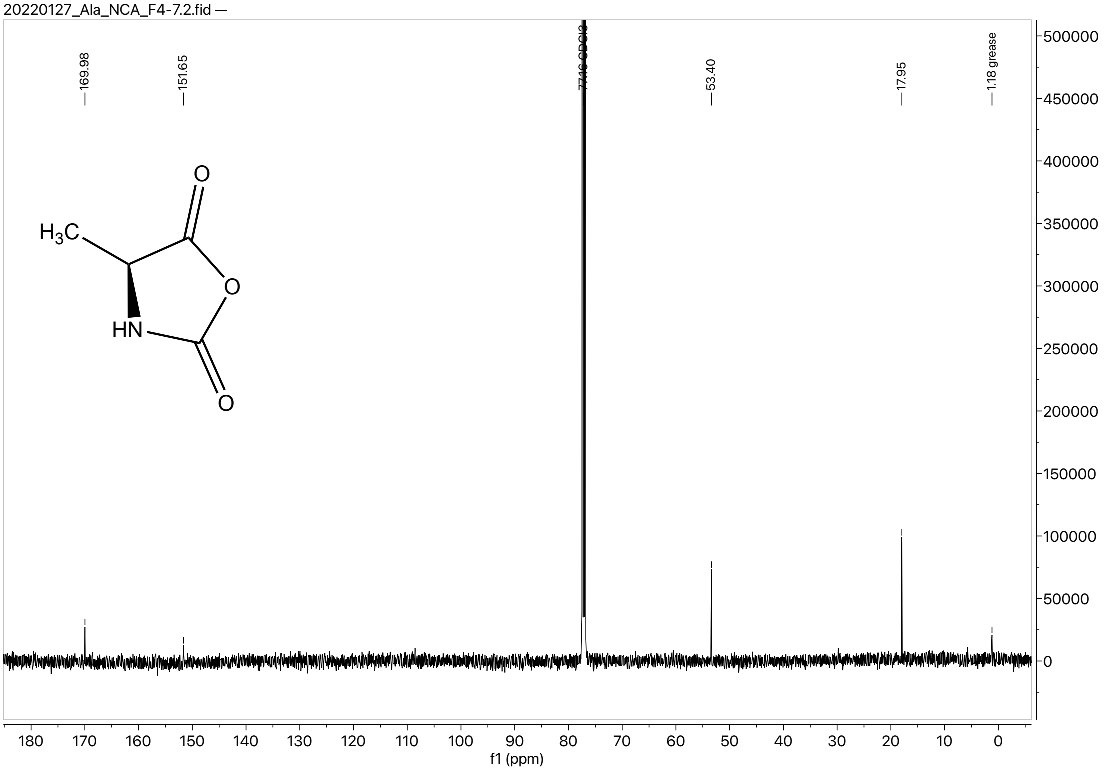

#### Z-Thr(βGalOAc$_4$-(1,3)-phenylisopropylidene-αGalN$_3$)-OBn

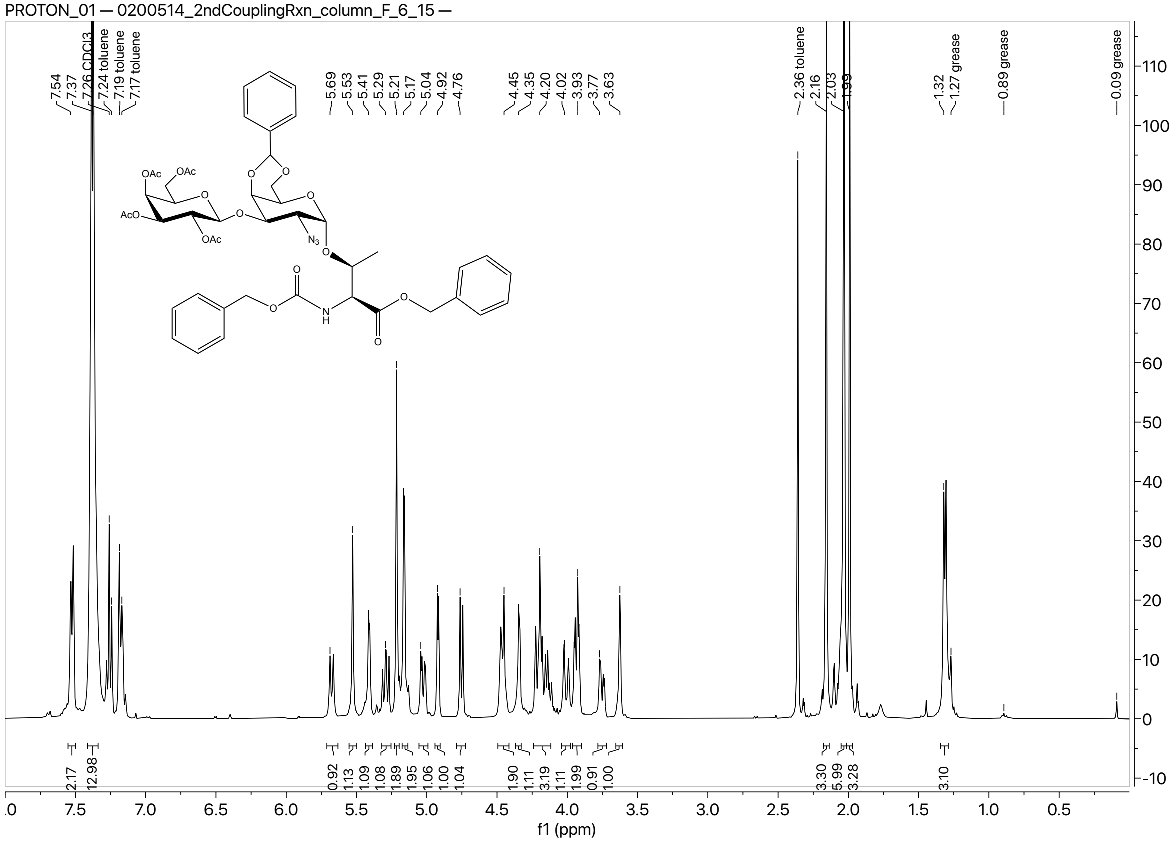

### Z-Thr(βGal(OAc)$_4$-(1,3)-αGalNAc(OAc)$_3$)-OBn

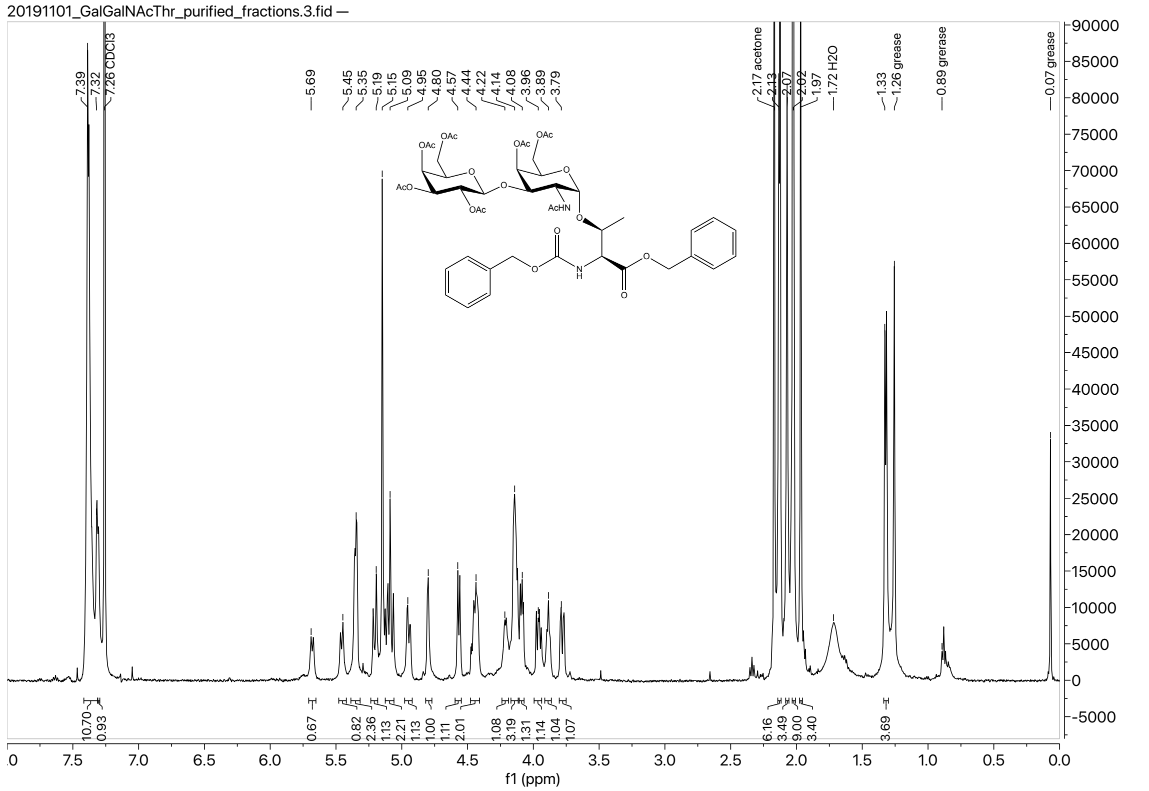

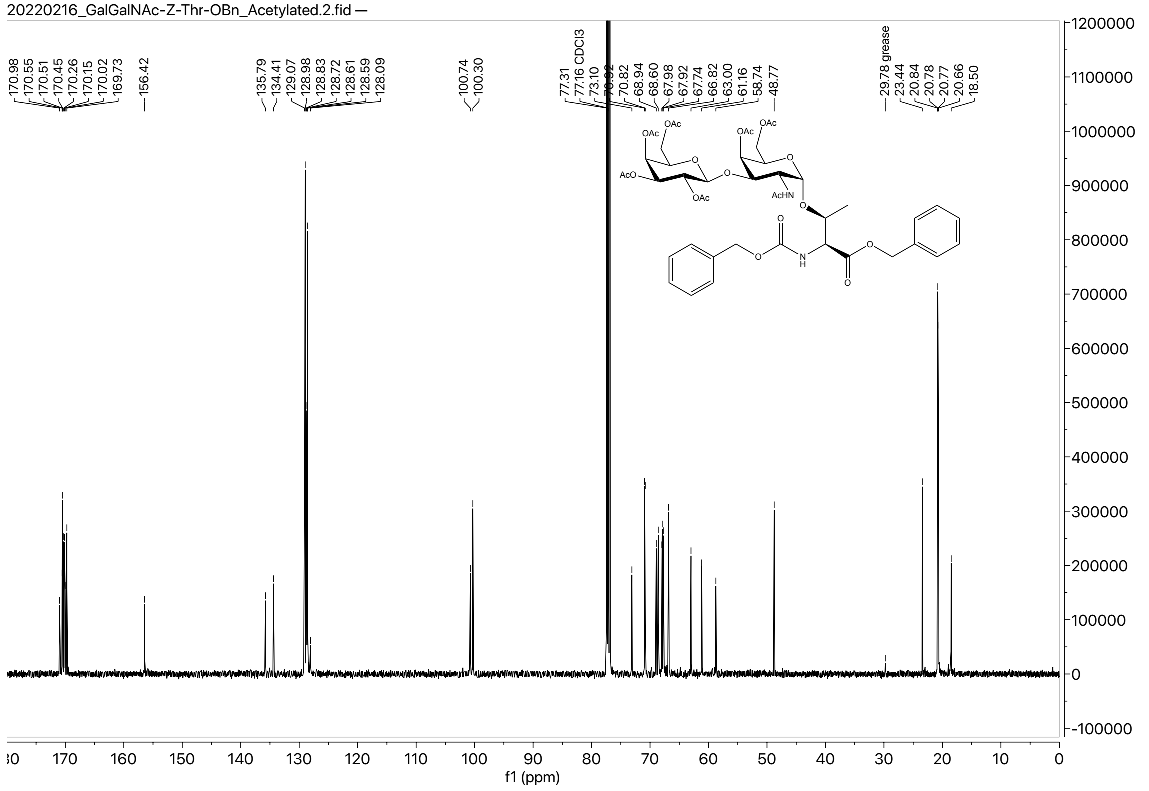

#### Boc-Thr(βGal(OAc)$_4$-(1,3)-αGalNAc(OAc)$_3$)-OH

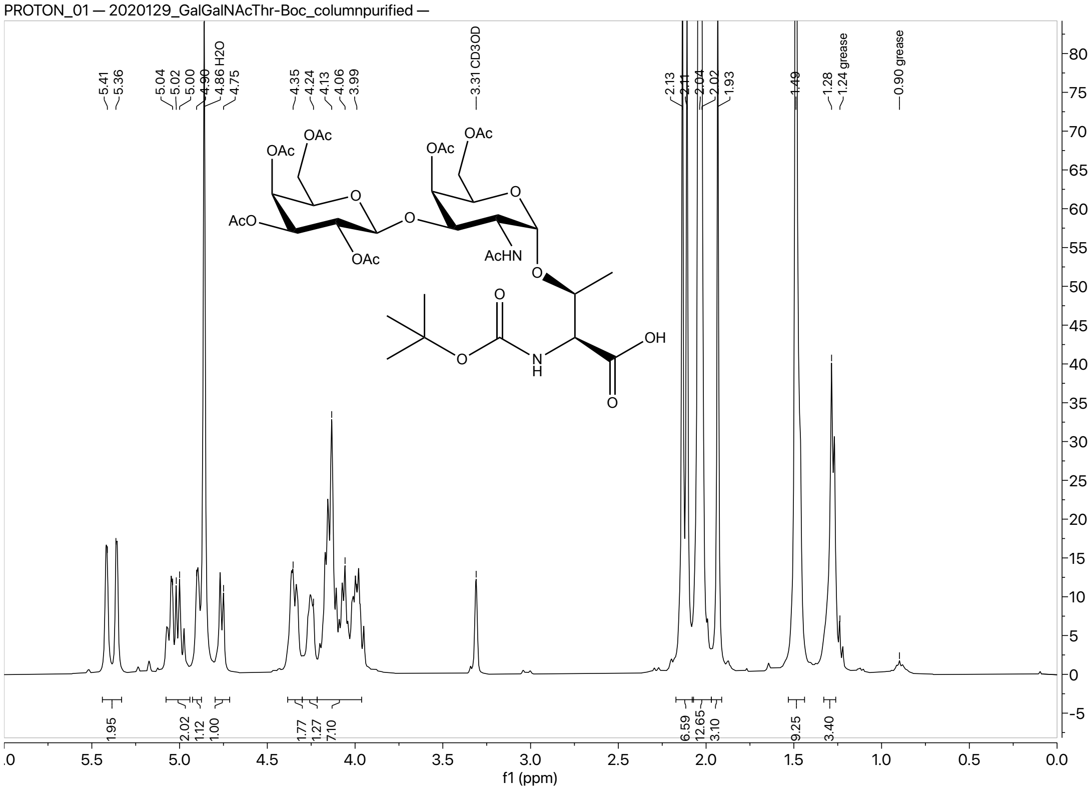

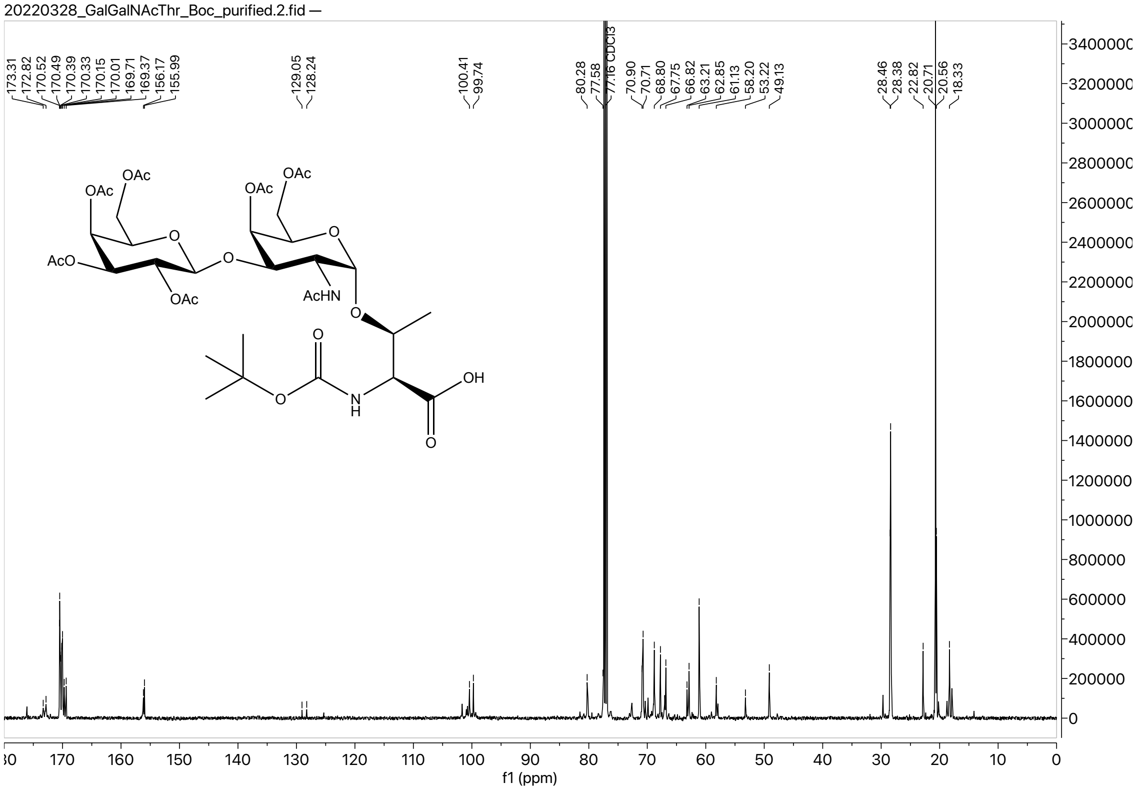

#### Thr(βGal(OAc)$_4$-(1,3)-αGalNAc(OAc)$_3$) NCA

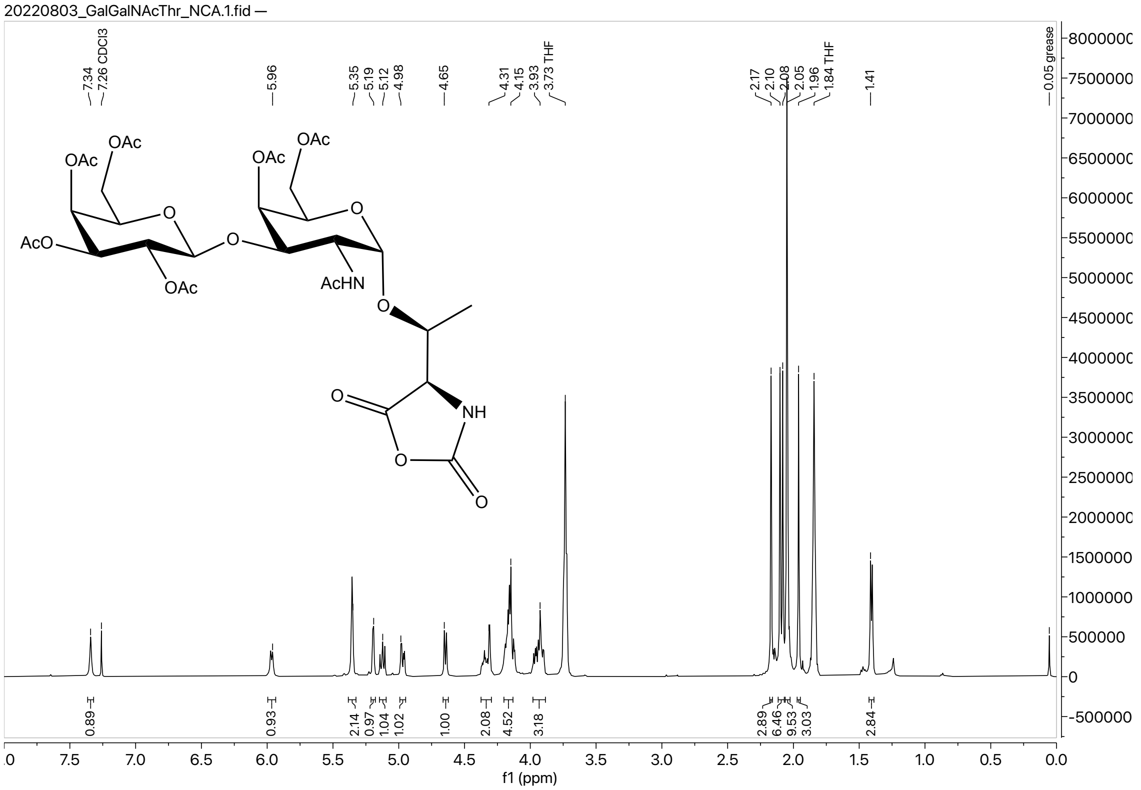

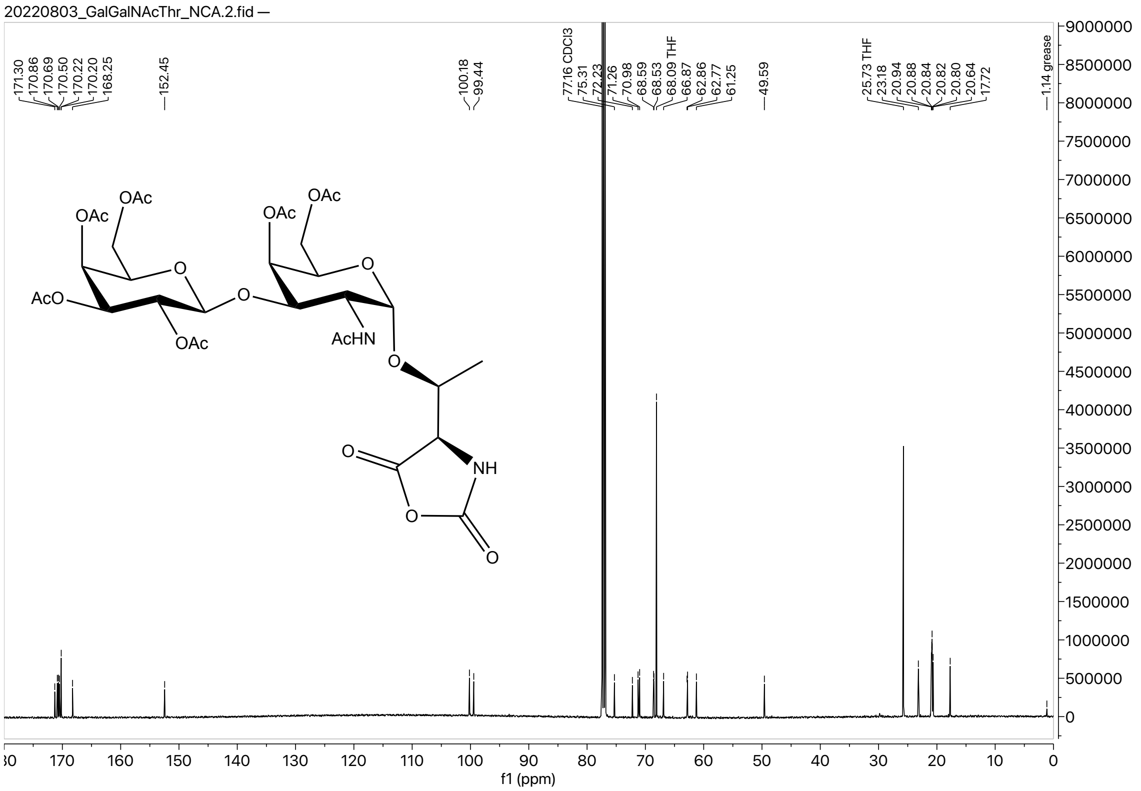

#### Z-Thr(βGalNAc(OAc)$_3$)-OBn

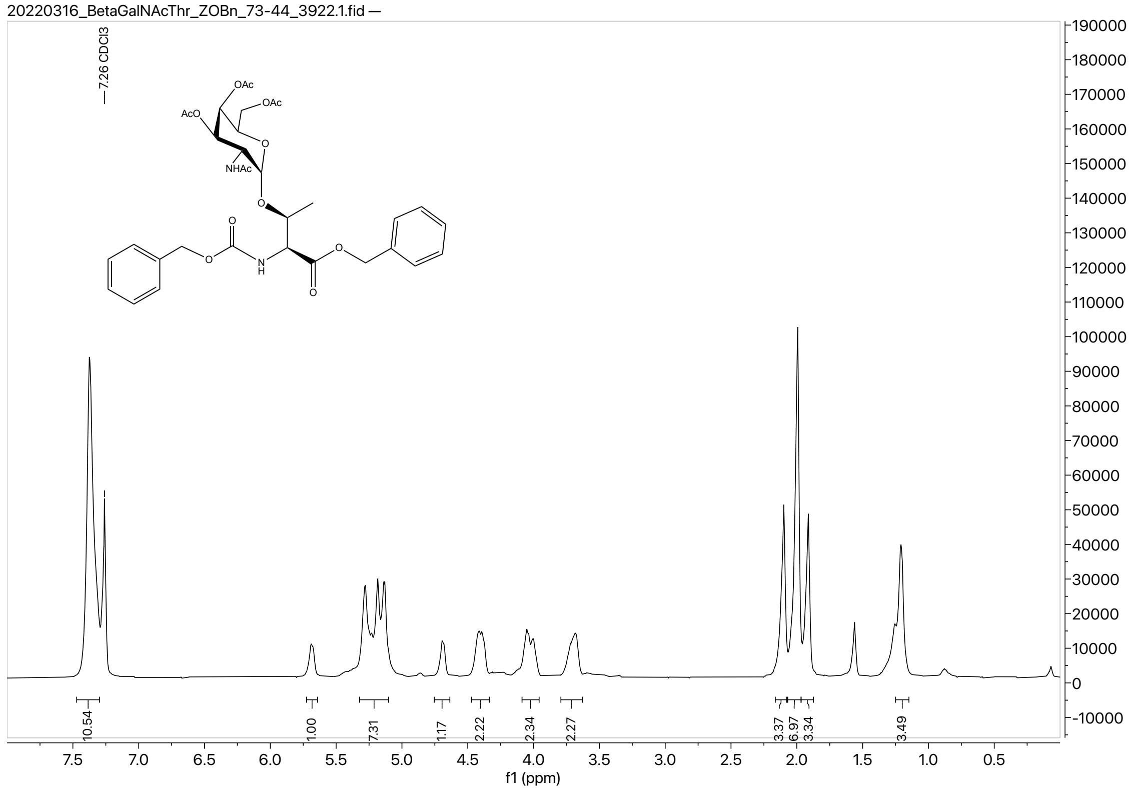

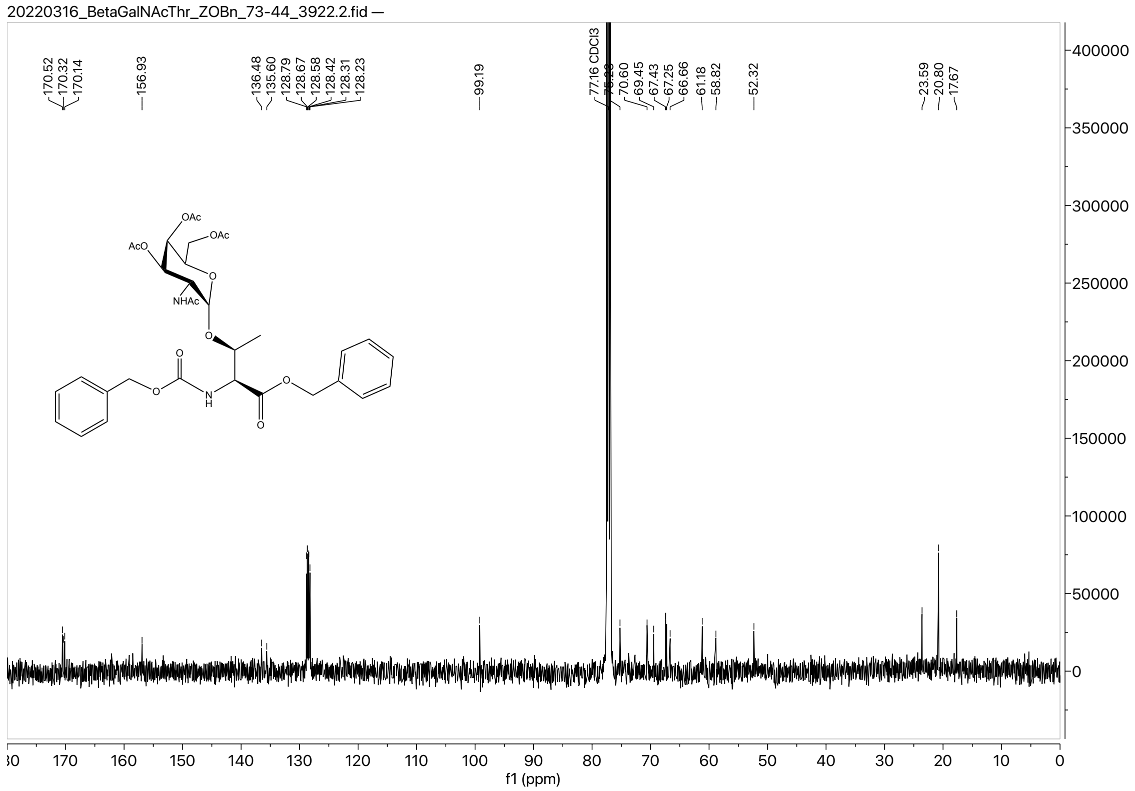

### Boc-Thr(βGalNAc(OAc)₃)-OH

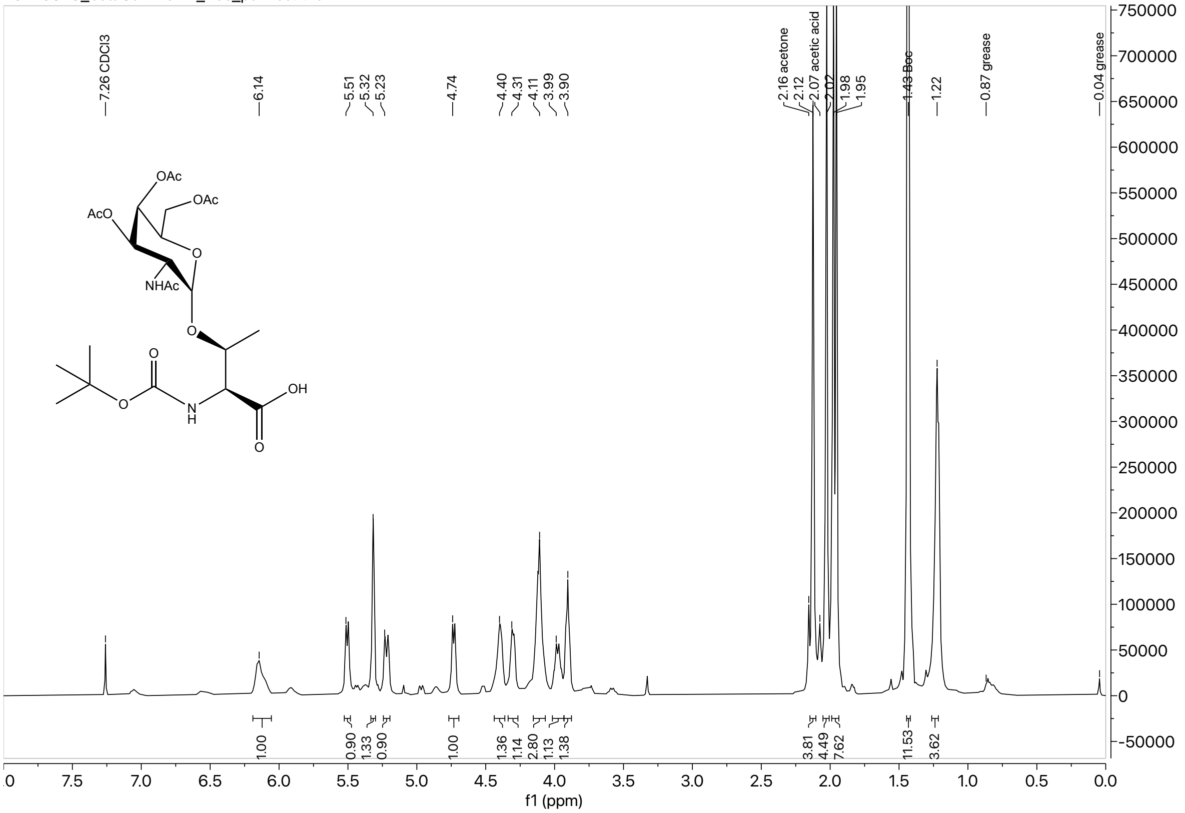

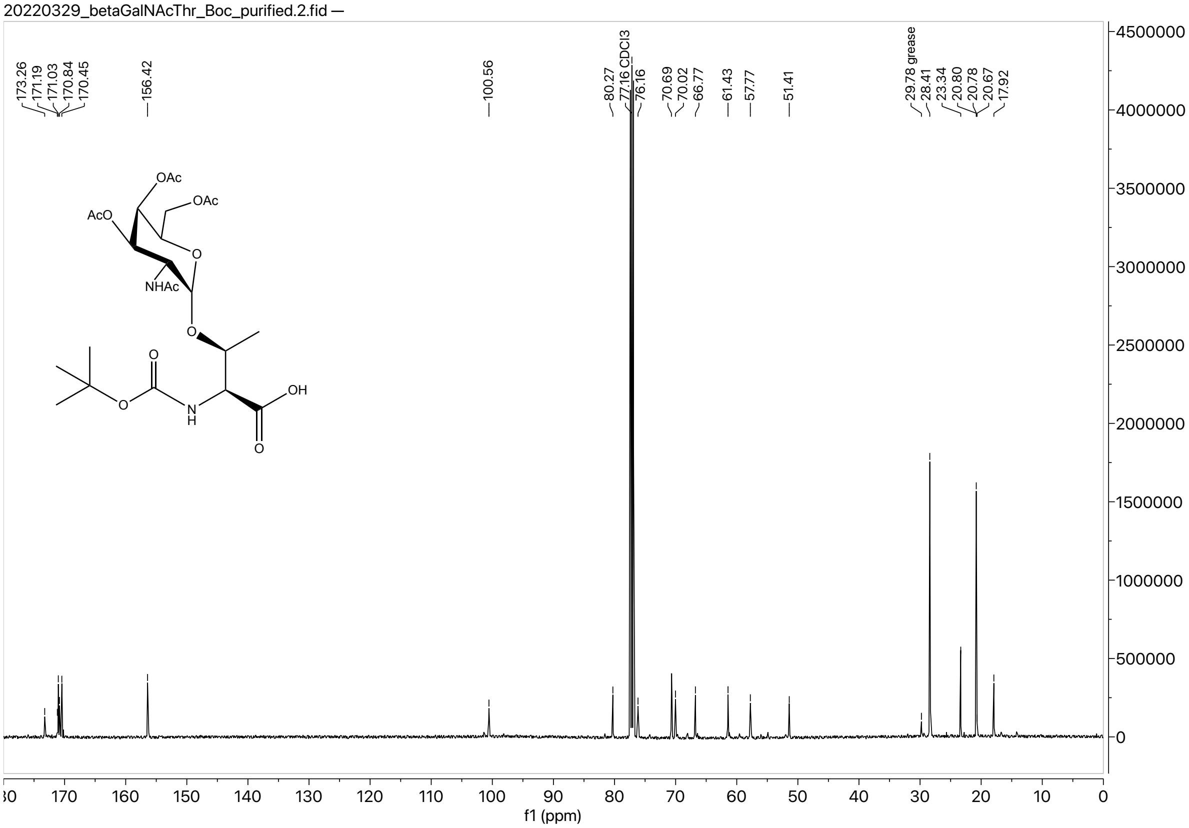

#### Thr(βGalNAc(OAc)$_3$) NCA

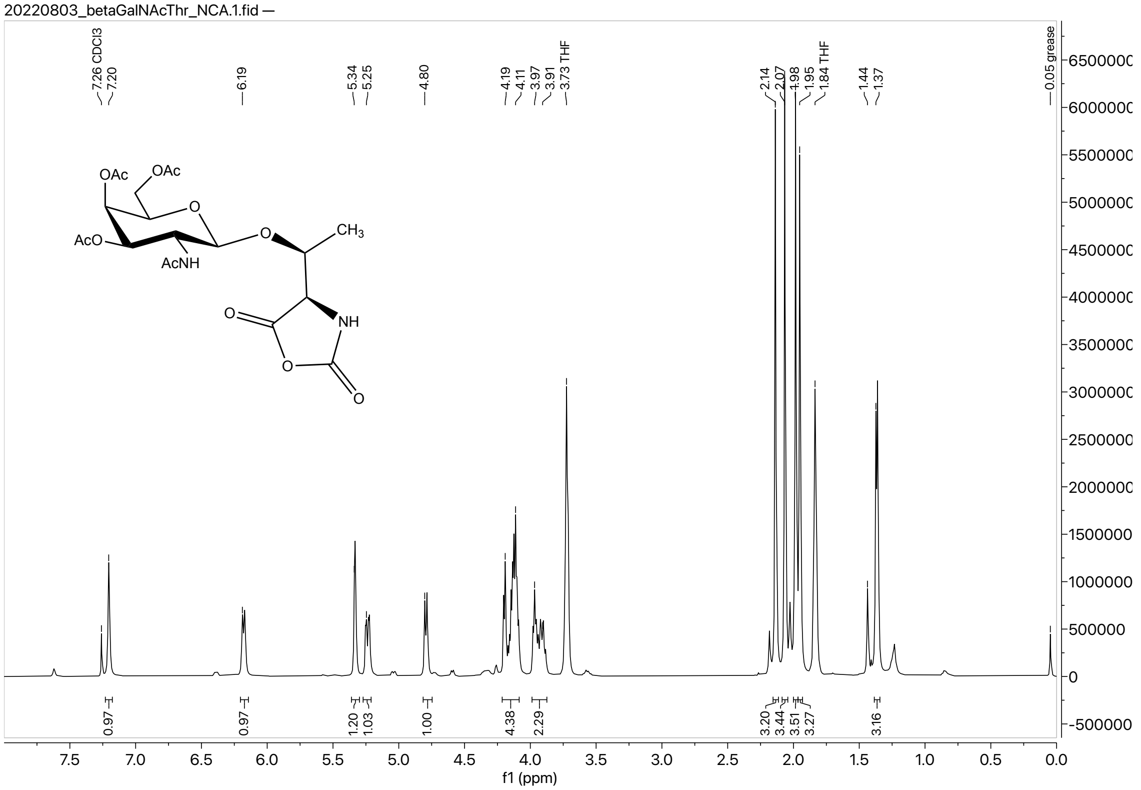

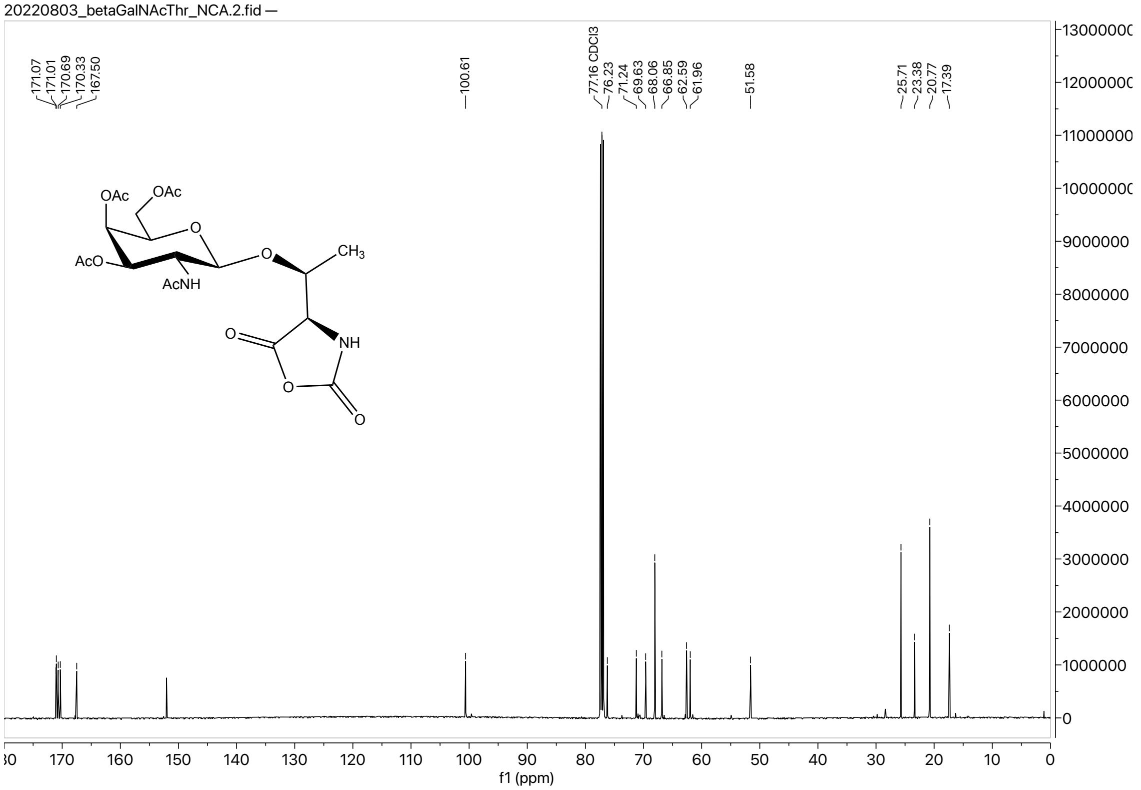

#### (βGalαGalNAcT$_{0.33}$-co-A$_{0.66}$)$_{93}$

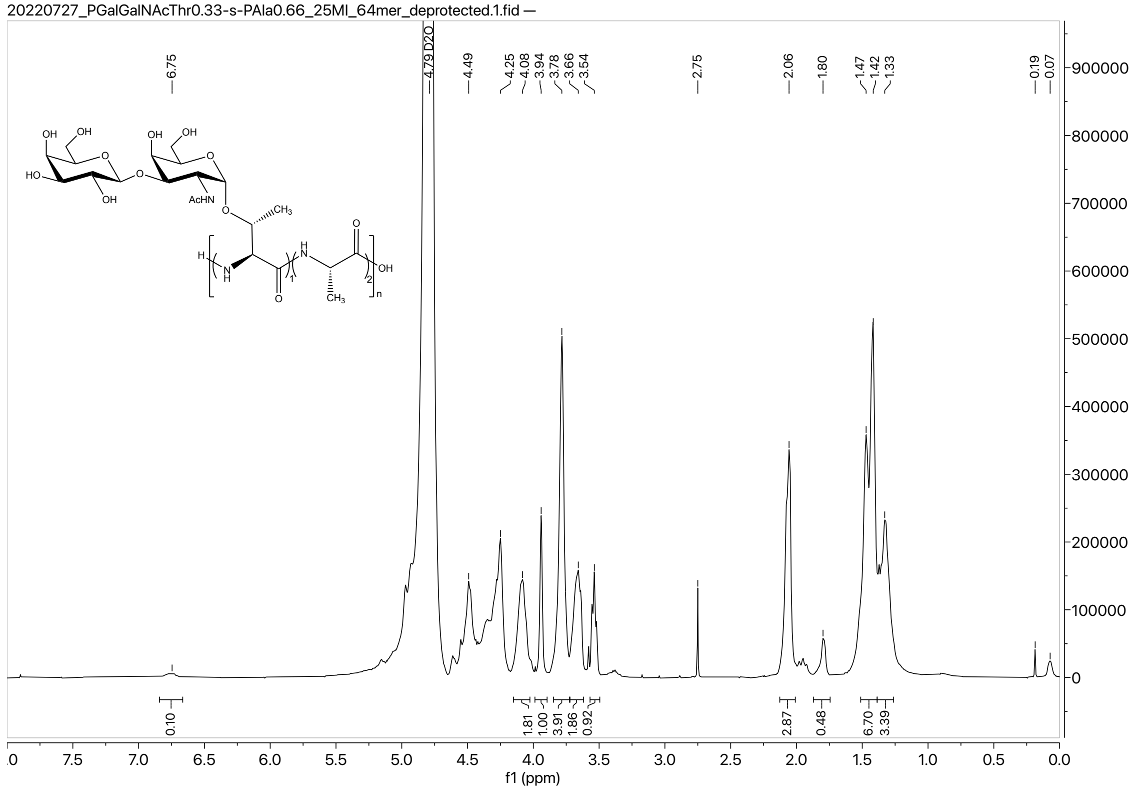

#### ($\alpha$GalNAcT$_{0.33}$-co-A$_{0.66}$)$_{93}$

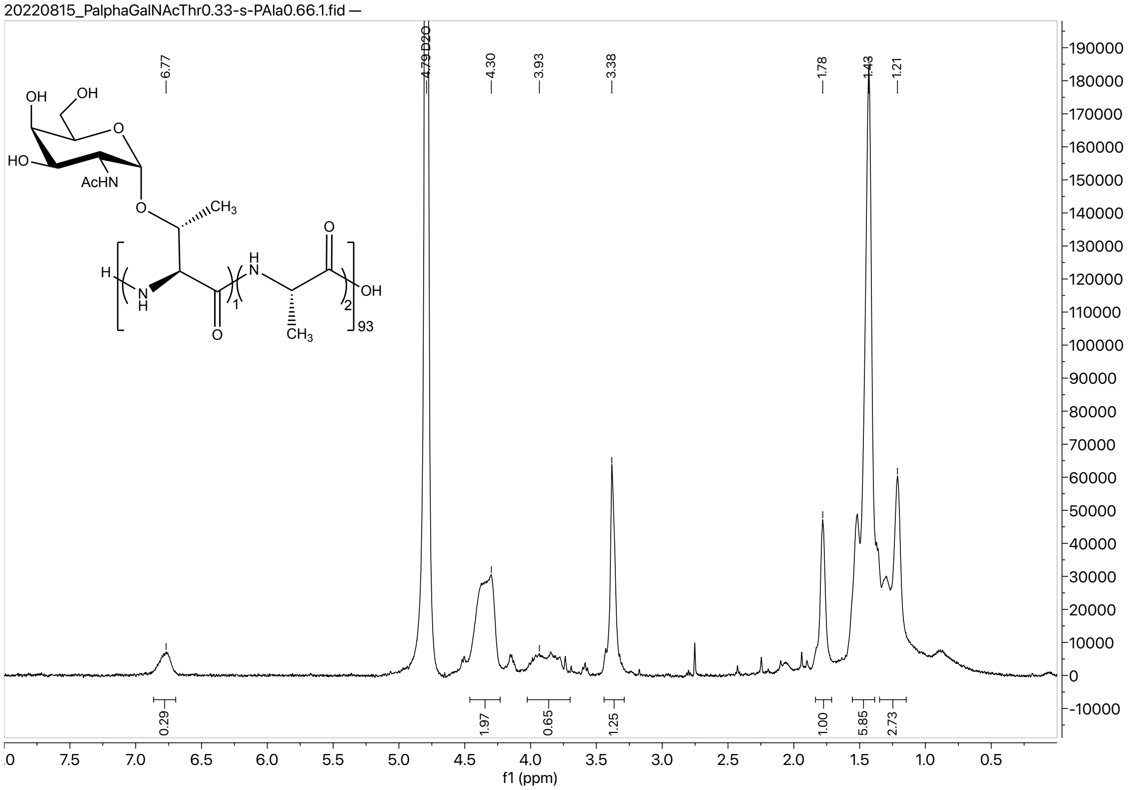

#### ($\beta$GalNAcT$_{0.33}$-co-A$_{0.66}$)$_{93}$

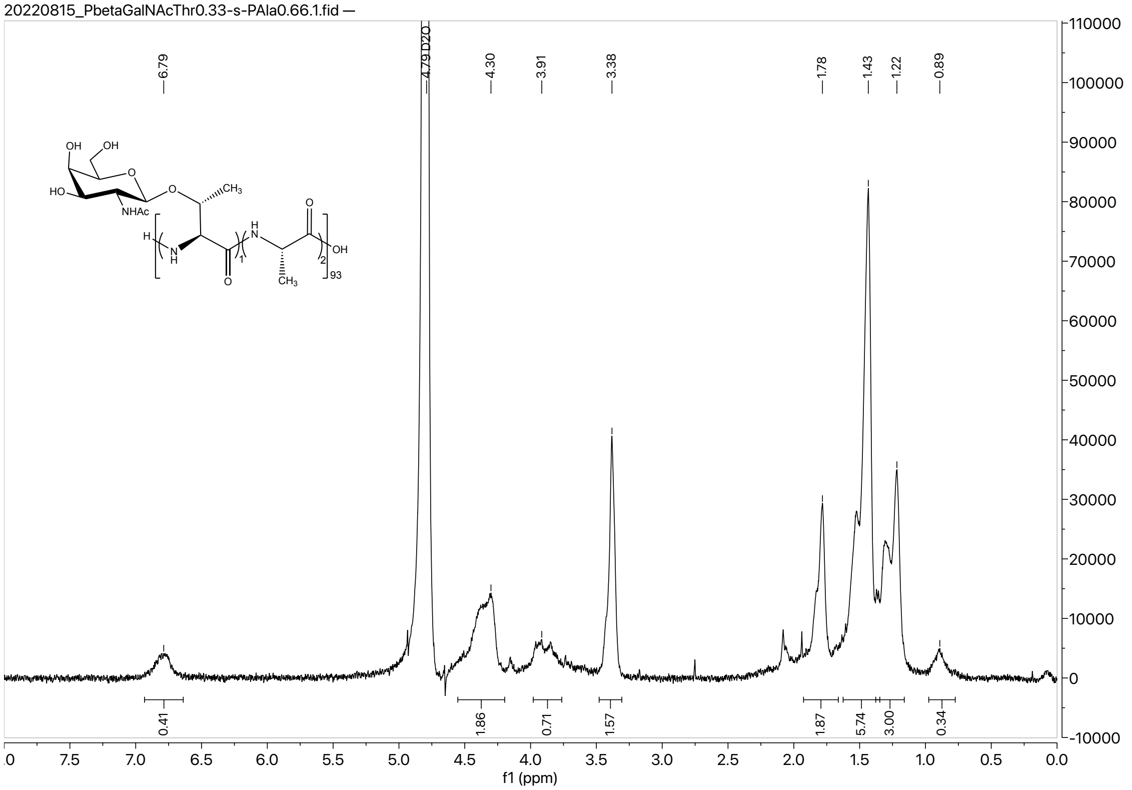

**($\alpha$GalT$_{0.33}$-co-A$_{0.66}$)$_{93}$**

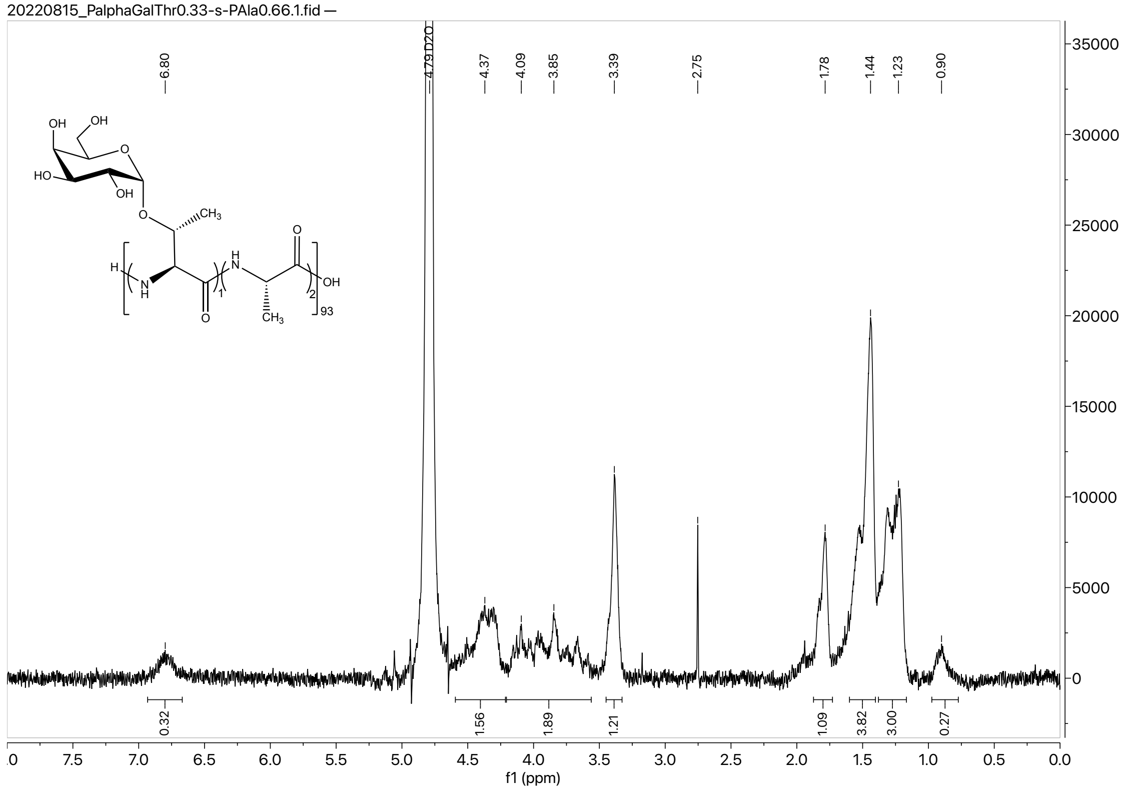

**($\beta$GalT$_{0.33}$-co-A$_{0.66}$)$_{93}$**

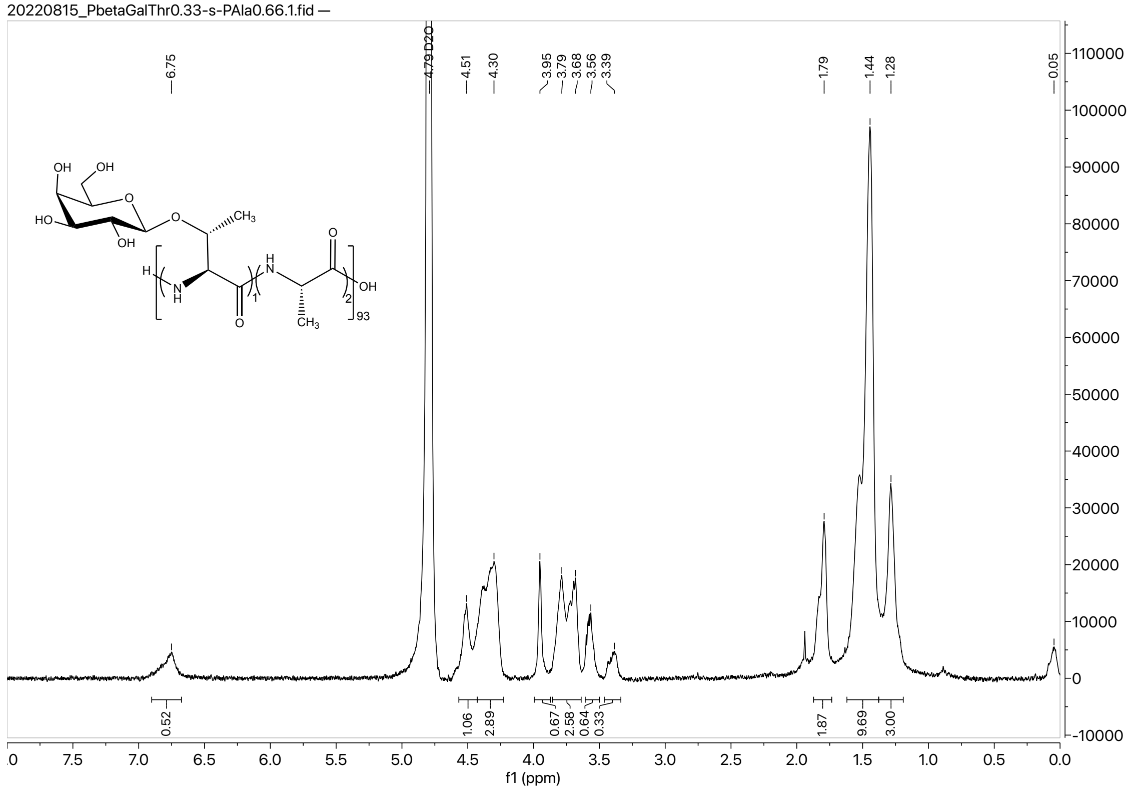

#### VII. ATR-FTIR Spectra

#### Ala NCA

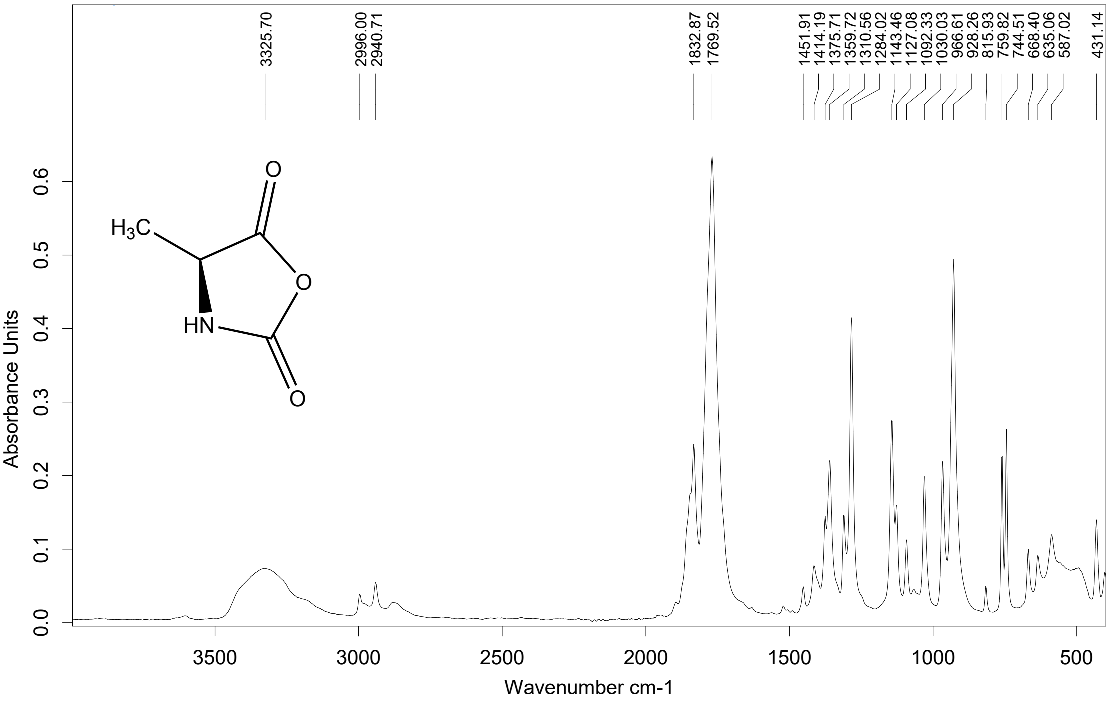

#### Boc- Thr(βGal(OAc)$_4$-(1,3)-αGalNAc(OAc)$_3$)-OH

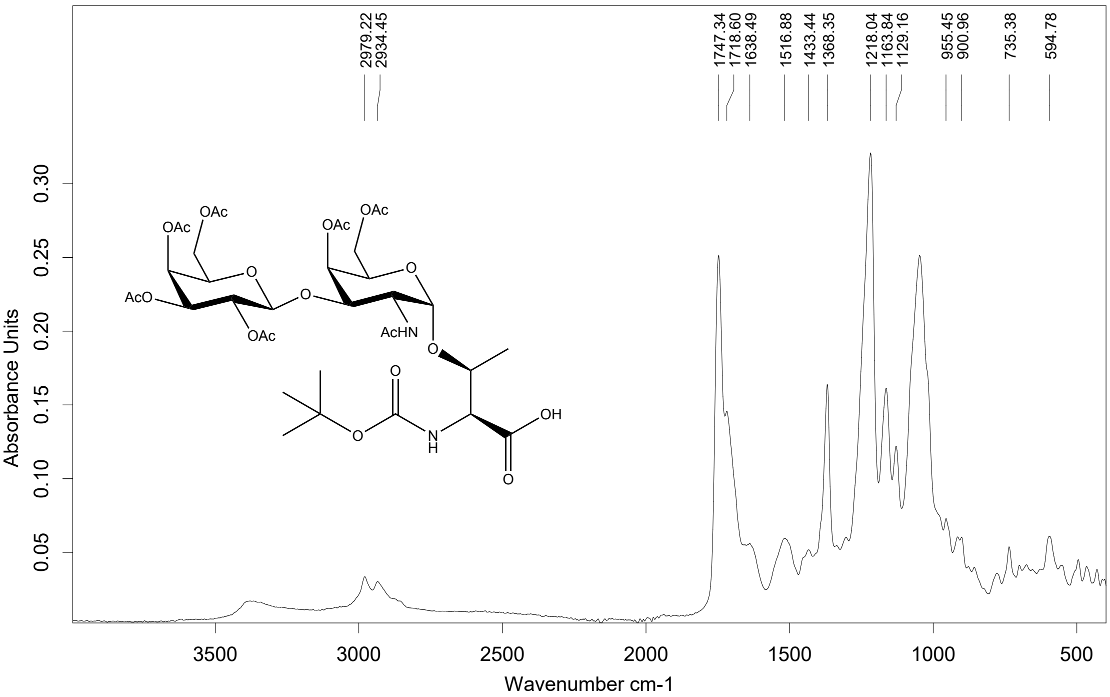

#### Thr(βGal(OAc)$_4$-(1,3)-αGalNAc(OAc)$_3$) NCA

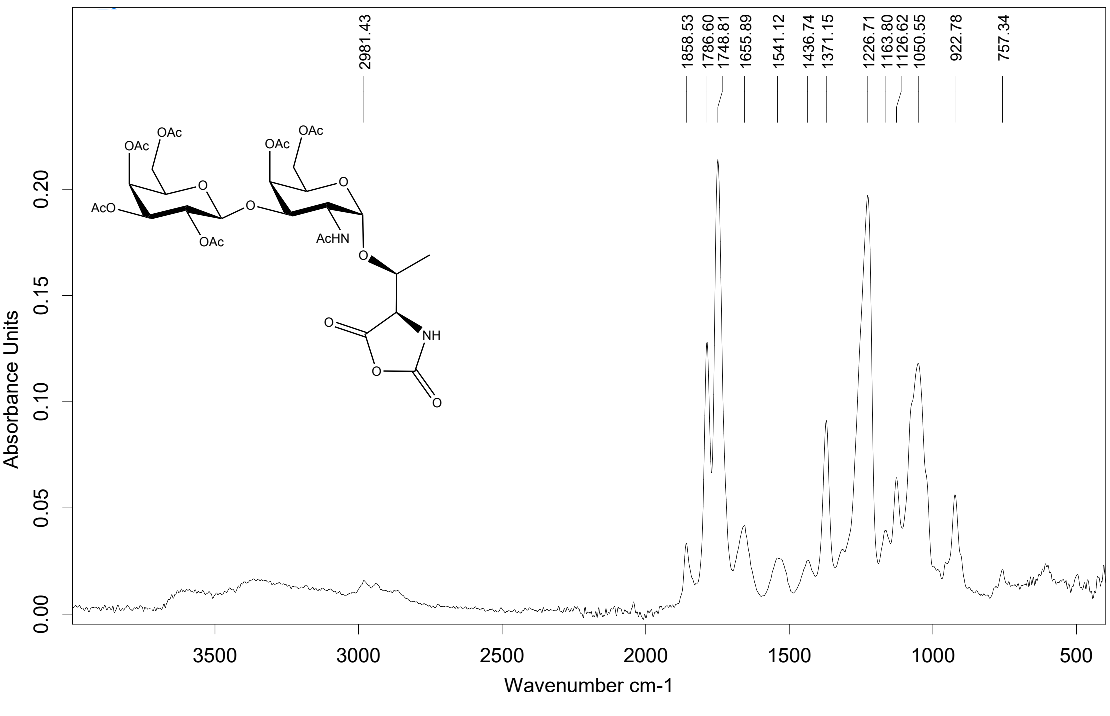

**(βGalαGalNAcT)$_{37}$**

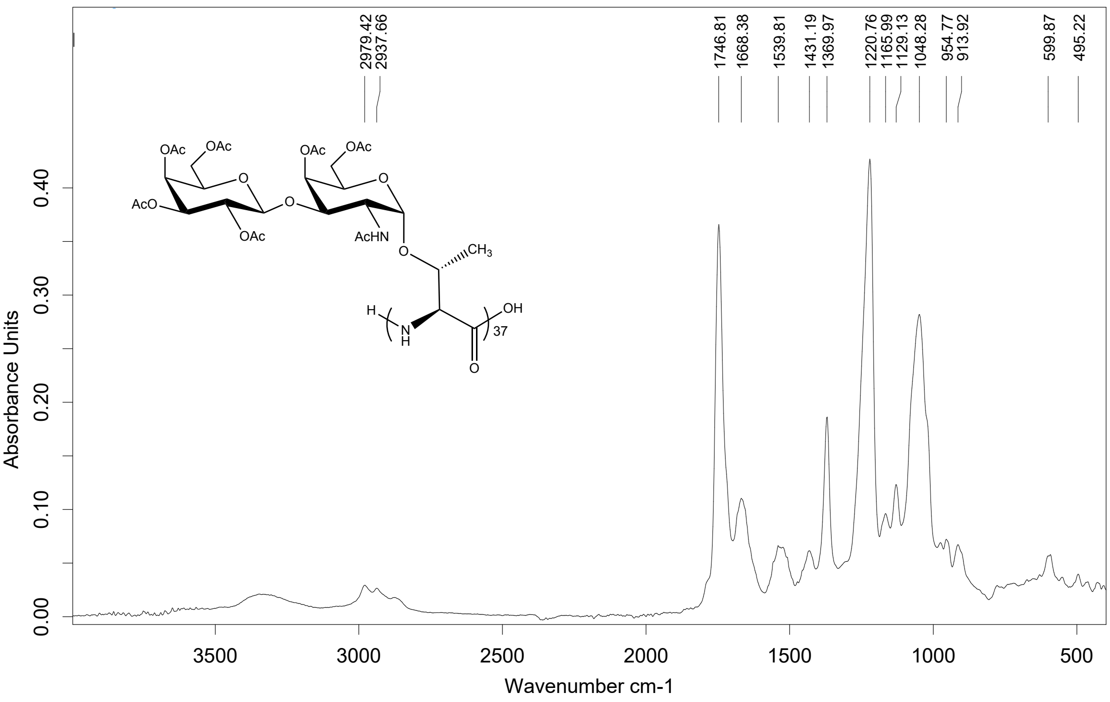

**Boc-Thr(βGalNAc(OAc)$_3$)-OH**

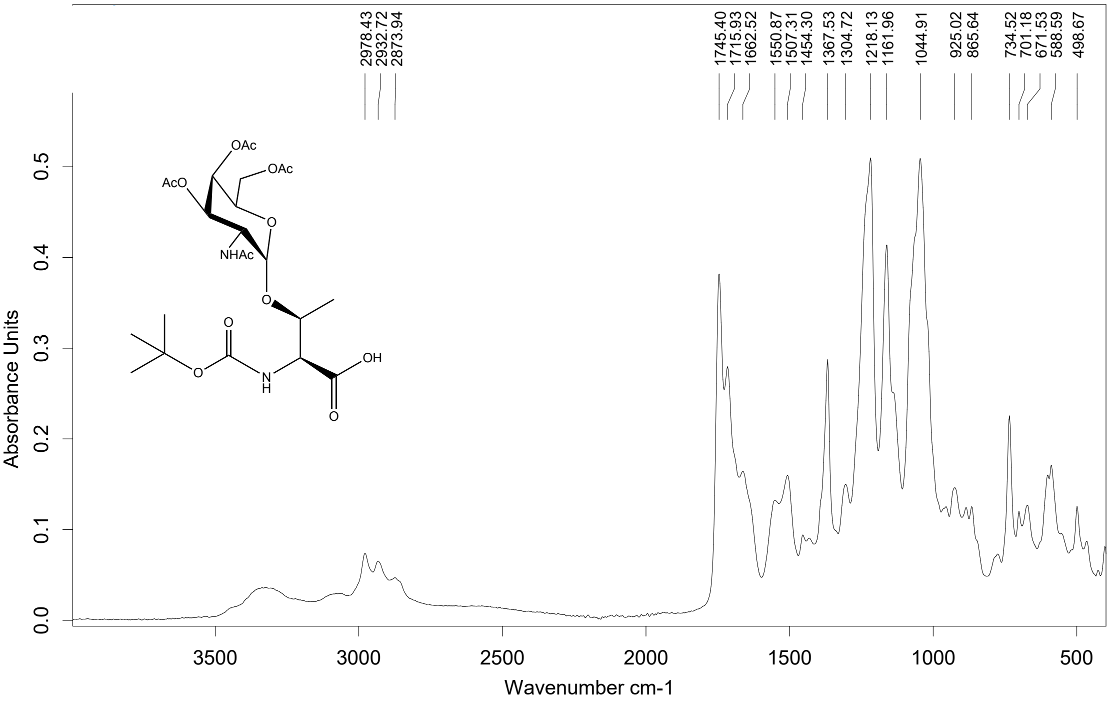

**Thr(βGalNAc(OAc)₃) NCA**

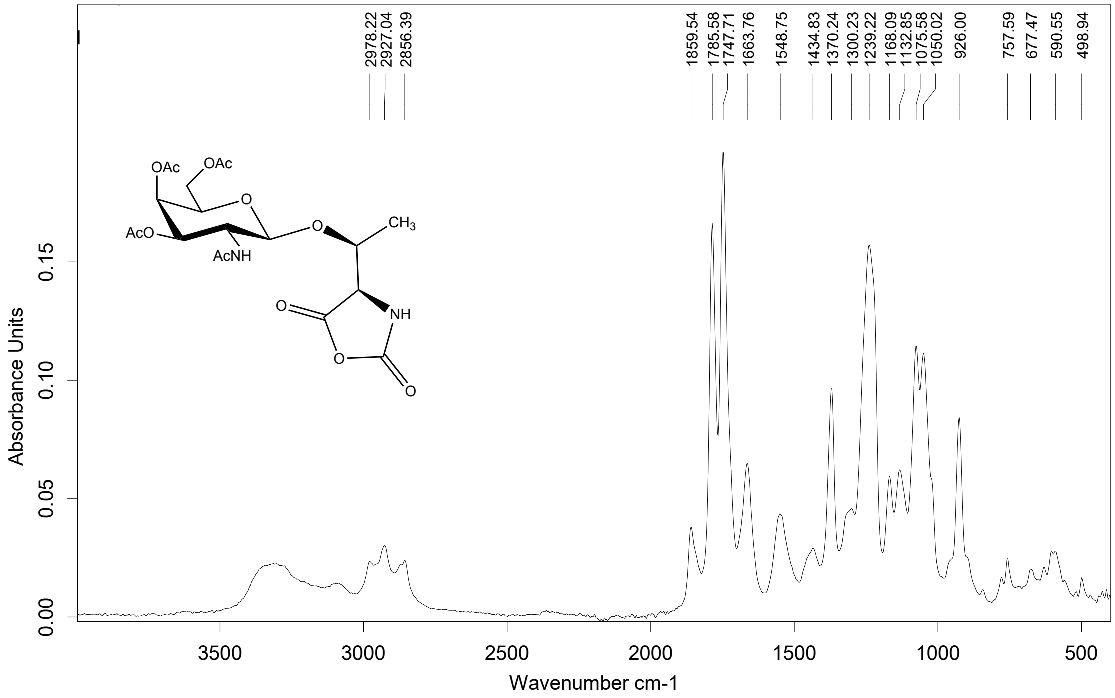

**(βGalNAc(OAc)₃T)₆₈**

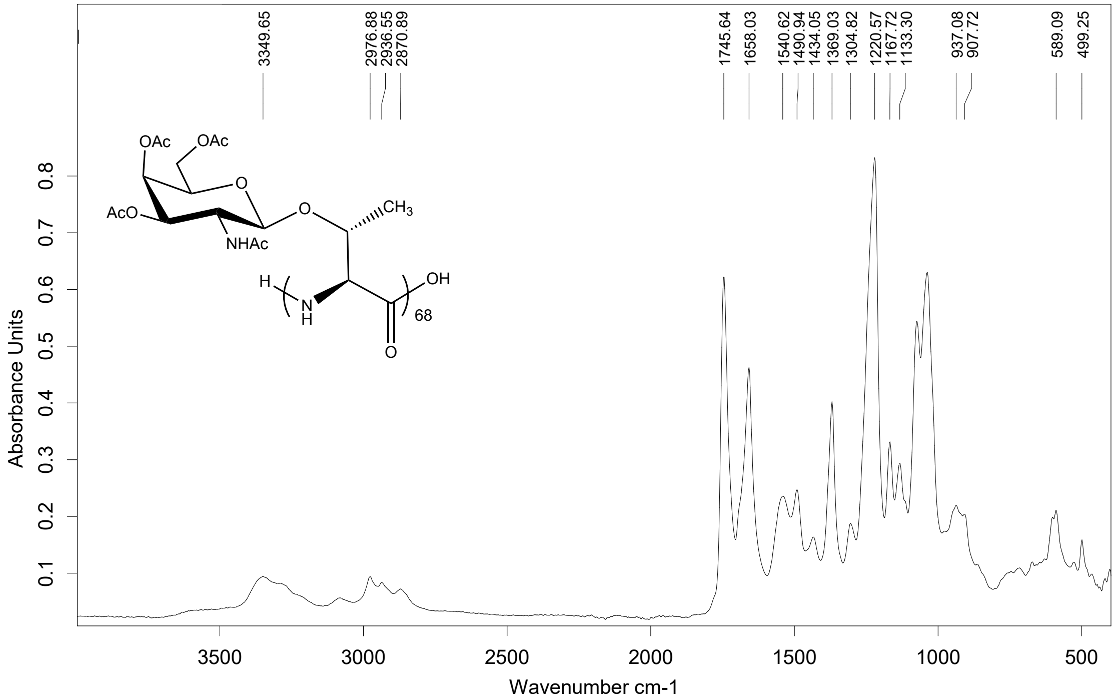

**(βGal(OAc)$_4$αGalNAc(OAc)$_3$T$_{0.33}$-co-A$_{0.66}$)$_n$**

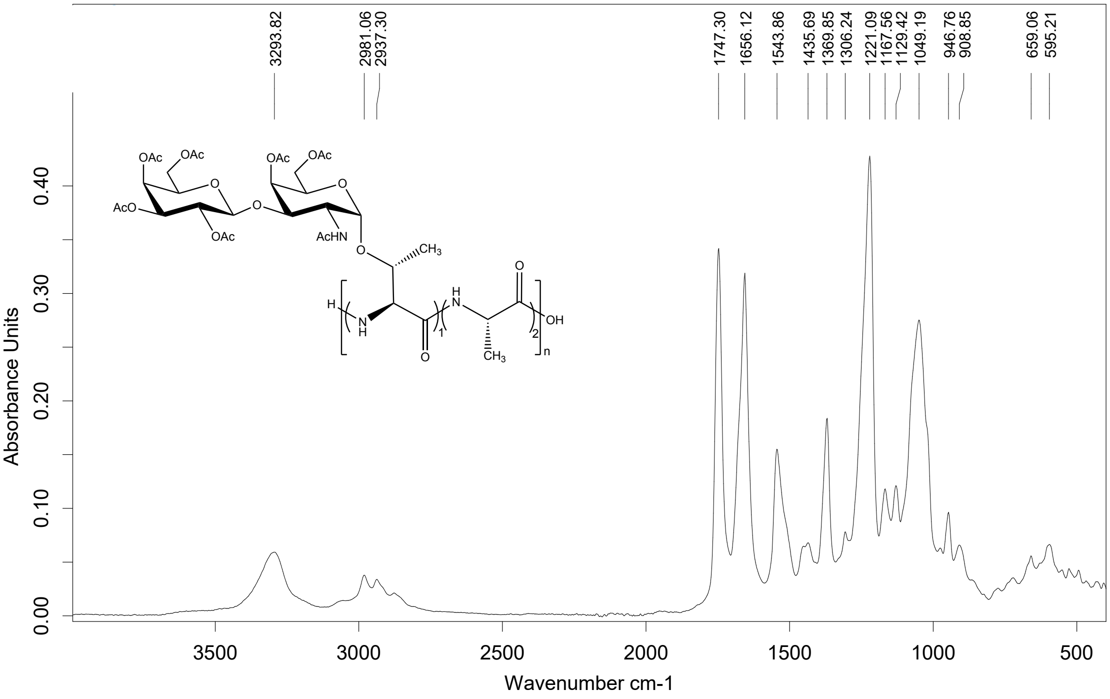

**(αGalNAc(OAc)$_3$T$_{0.33}$-co-A$_{0.66}$)$_{93}$**

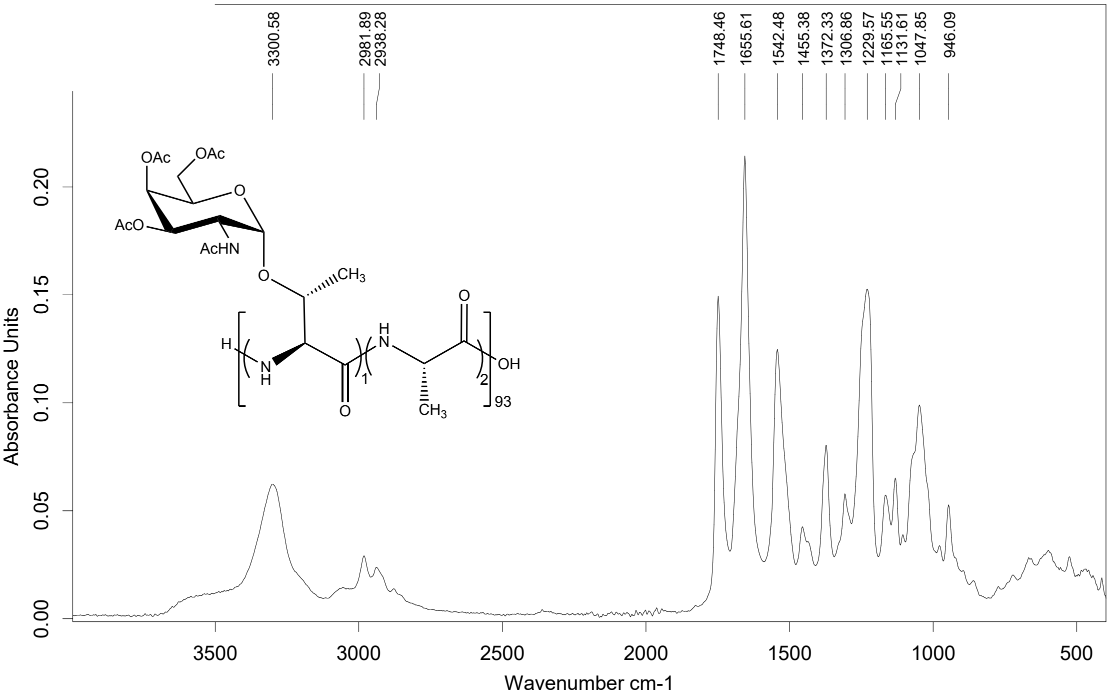

**($\beta$GalNAc(OAc)$_3$T$_{0.33}$-co-A$_{0.66}$)$_{93}$**

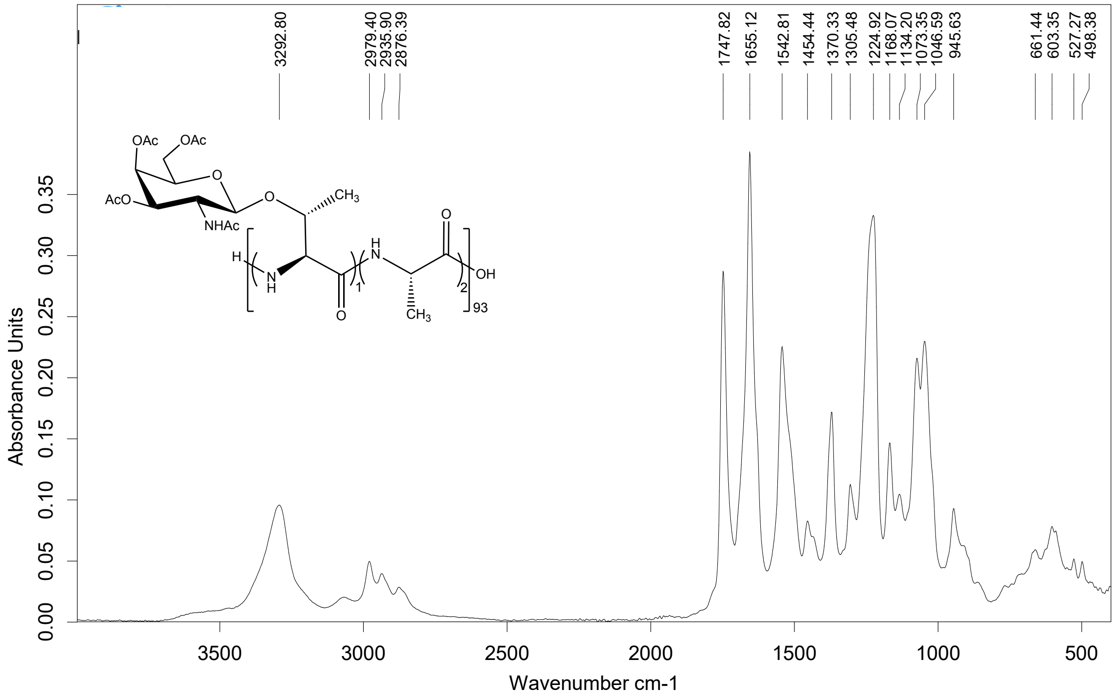

**($\alpha$Gal(OAc)$_4$T$_{0.33}$-co-A$_{0.66}$)$_{93}$**

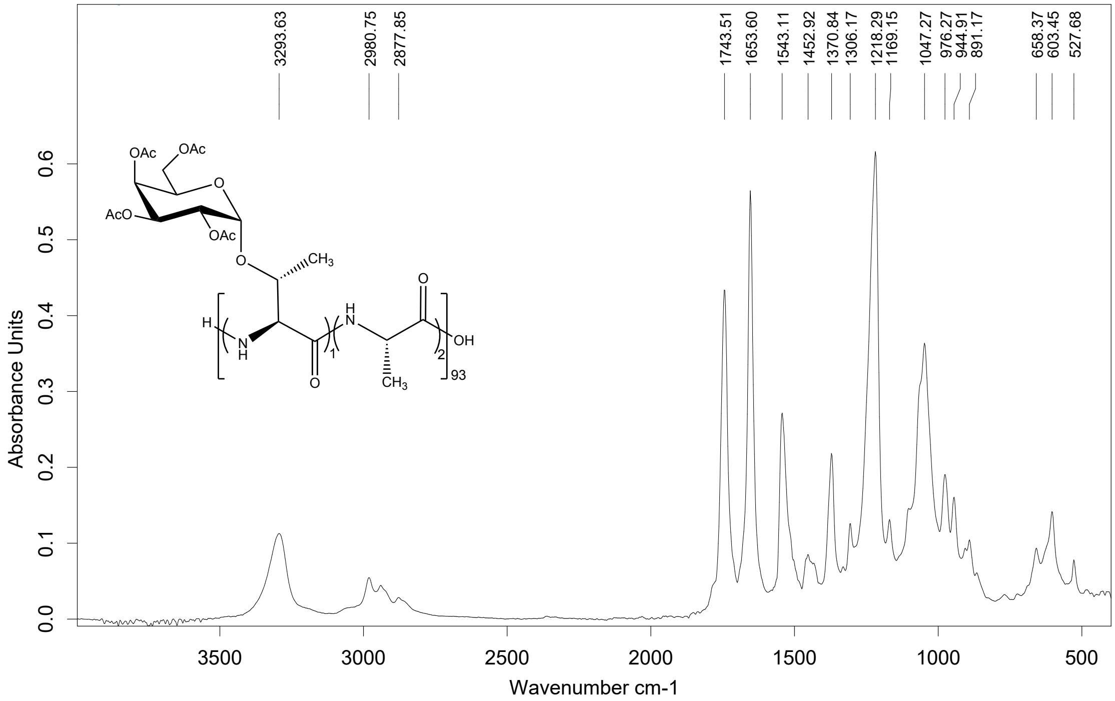

**(βGal(OAc)$_4$T$_{0.33}$-co-PA$_{0.66}$)$_{93}$**

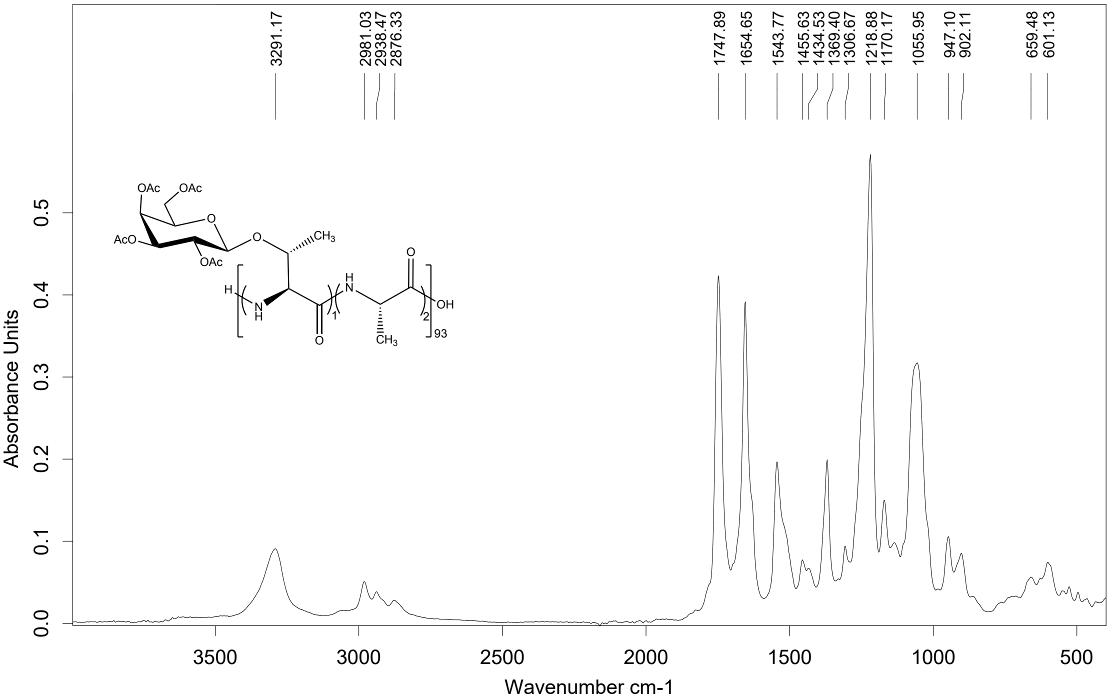

**VIII. References**

(1) Fulmer, G. R.; M Miller, A. J.; Sherden, N. H.; Gottlieb, H. E.; Nudelman, A.; Stoltz, B. M.; Bercaw, J. E.; Goldberg, K. I.; Beckman, M. NMR Chemical Shifts of Trace Impurities: Common Laboratory Solvents, Organics, and Gases in Deuterated Solvents Relevant to the Organometallic Chemist. *Organometallics*. https://doi.org/10.1021/om100106e.
